# Fast timescale reparameterization of stable neural manifolds

**DOI:** 10.64898/2026.01.26.701671

**Authors:** Stephen E. Clarke, Paul Nuyujukian

## Abstract

The prevailing opinion in systems neuroscience is that motor cortex generates consistent behavior through a stable neural manifold—a low-dimensional mapping from neuron population activity to kinematics. However, standard population analyses group many repeated trials to reduce estimation noise, obscuring the time-varying activity of the network. Here, a block-by-block analysis of task trials in two species revealed that motor cortex rotates its principal subspaces as the contribution of individual neurons fluctuates, maintaining stable dynamics that are coupled to the neural state. While static decoders performed inconsistently due to non-stationary neuron activity, we confirmed that decoding performance could be rescued through subspace alignment, finding that roughly half of this geometric correction was driven by the internal rank-swapping of dominant state eigenvectors. Rather than relying on a static mapping to execute reaching behavior, motor cortex maintains a library of redundant state eigenvectors that generalize over different reach directions. Model simulations demonstrated that strong excitatory coupling combined with spike-rate adaptation is sufficient to produce a latent library whose elements can swap rank during continuous reparameterization of the network state. Together, these results refine our understanding of motor control: stable neural manifolds drift and jump within bounded limits, continuously outrunning neuron fatigue during repetitive activation.

## Introduction

The current state of systems neuroscience provides a complex view of how representations in neuron populations change over time. Representations in cortical neuron populations are known to slowly drift between repeated experimental sessions, despite the associated task behavior remaining stable for days, even years.^1–7^ While the average response of these individual neurons can be stable over long timescales,^8–11^ particularly in the motor system, they can also change quickly with respect to task behavior during single experimental sessions.^12–18^ Therefore, it’s reasonable to expect that correlations among a population of neurons could also change quickly over time,^19^ especially since they can shift rapidly under different behavioral task contexts.^20,21^ Recent theoretical modeling demonstrates that topologically distinct circuit structures and single-neuron functional properties can produce equivalent low-dimensional population dynamics.^22^ This degeneracy suggests that stable collective behavior does not require static connectivity or fixed single-neuron loadings for neural dimensions, consistent with earlier theories of motor learning.^17^

Understanding representational drift over faster timescales is central; if the neural manifold is structurally stable during an experimental session, it presents a paradox as to why brain-computer interface decoder performance degrades during continuous operation.^23,24^ To resolve this tension between the long-term stability of neural manifolds and the underlying churn of individual neurons, we tracked the mapping of neuron contributions to the population state dimensions over contiguous blocks of task trials during single experimental sessions. A recent breakthrough in motor cortex has demonstrated that the neural manifold is not entirely fixed, but acts as a parameterized dynamical system where the classic plane of rotation^25^ shifts based on a target’s spatial location.^21^ Rather than viewing the evolution of neural activity as random gradual drift or noisy sampled neurons, we hypothesize that motor cortex extends this flexibility by utilizing redundant, shifting encoding patterns parameterized by time to maintain a stable behavioral output.

## Results

### Non-stationary neuron population activity underlies consistent reaching behavior

To explore whether there are fast-timescale changes of individual neuron contributions to population state dynamics, we analyzed chronic neuron activity recorded from layer 2/3 of the motor cortex while rhesus monkeys performed a stereotyped center-out reaching task with their contralateral arm (Figure 1a; see Methods for full detail). Kinematics were derived from the 2D task cursor and the associated neuronal activity was analyzed in 25ms bins over the interval [-50,400]ms around the estimated reaction time, where the variability of the behavior and the neuron population activity is known to decrease significantly.^26^ Data shown for an exemplar session from Monkey U, 2020-11-30, used to illustrate results in this article. Instead of concatenating trials or computing trial averages under the assumption of a stationary process, recording sessions were broken into sequential blocks of trials to independently evaluate the temporal evolution of the reach representation in motor cortex (16 trials for Monkey U, 2 trials per 8 targets; 12 trials for Monkey H, 2 trials per 6 targets).

**Figure 1:**
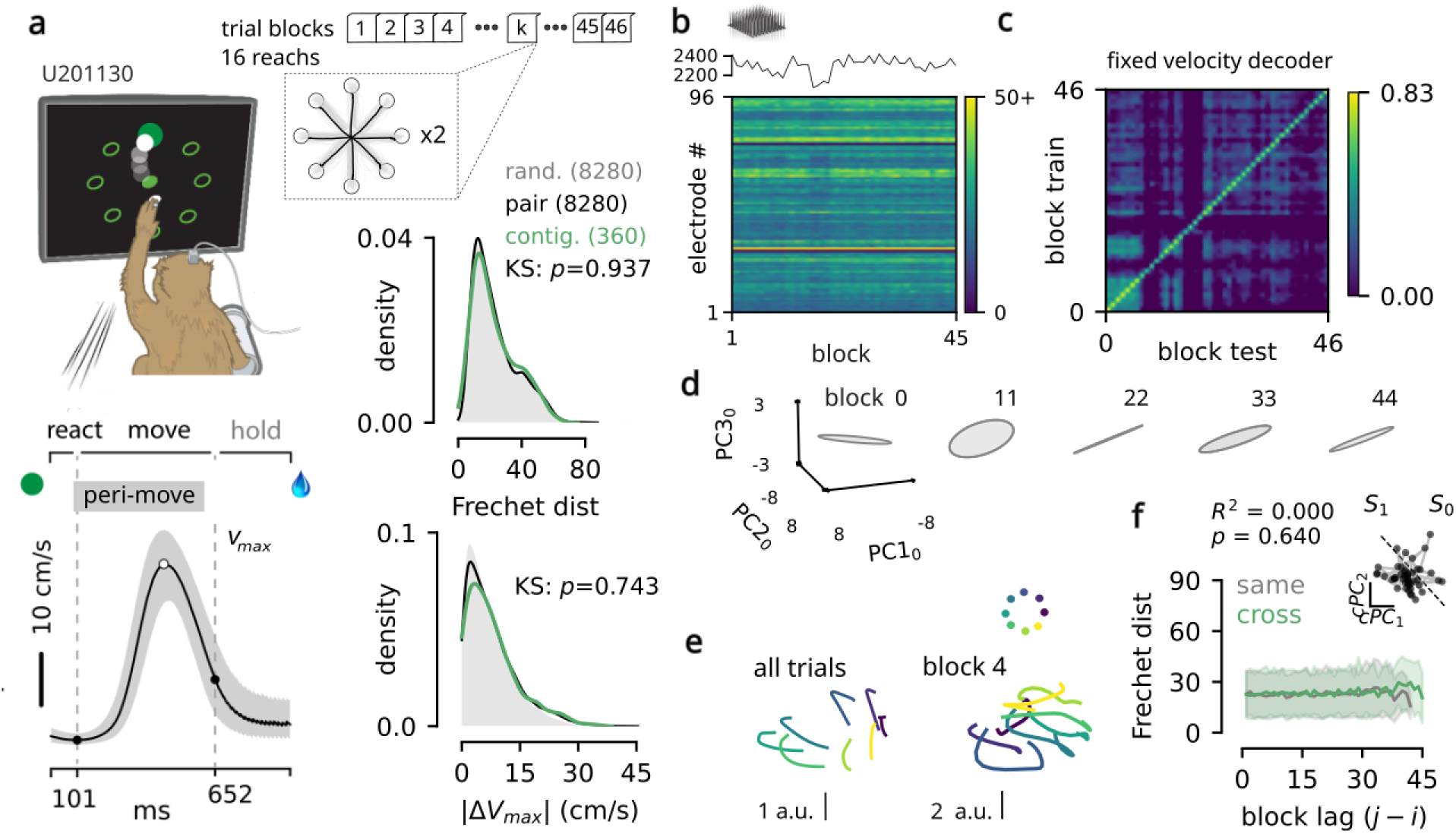
Non-stationary neuron population activity underlies consistent reaching behavior. **[a]** During single experimental sessions, two experienced rhesus macaques performed hundreds of continuous center-out reach trials while neuron populations were recorded from motor cortex. Trial-averaged reach speed is shown for all 750 successful trials in an example 20 minute session (Monkey U, 2020-11-30). For each session, trials were grouped into sequential, target-balanced blocks, where each trial was aligned to movement onset to isolate a stereotyped move phase, minimizing the influence of kinematic variability on the observed population response^25,27^ (see Methods). There were negligible changes to *v_max_* and the Fréchet distance between reach trajectories measured across successive blocks and those sampled randomly throughout the recording session (see also Supplementary Figures S1-S4). **[b]** Mean population spike count per block recorded by the 96-channel array (*top*), and mean spike counts for individual electrodes (*bottom*). Population activity remains largely stable but exhibits variability throughout the session (Supplementary Figure S5). **[c]** Despite the behavioral invariance, a kinematic ridge-regression decoder trained on a given block of trials exhibited patchy cross-block generalization. **[d]** When projected into the active subspace defined during block 0, the neural trajectories of subsequent blocks dynamically rotate and scale (visualized as covariance ellipses capturing a snapshot of the neural manifold at each block *t*). **[e]** Comparing conventional state-space trajectories pooled across the entire session against block-specific dimensionality reduction confirms that the underlying task geometry remains intact; although individual blocks are inherently noisier due to lower trial counts, the network successfully separates the 8 targets. **[f]** Kinematics were compared between pairs of blocks that had rotated into opposing principal halves of the neural state space, revealing no significant differences over time.

First, we verified that any putative changes in neural activity were not simply artifacts of changing kinematics.^27^ All trials were performed with the same posture and the ipsilateral limb held in a steady position.^28^ We quantified the behavioral similarity between trial blocks using the maximum trial velocity (*v_max_*) and the discrete Fréchet distance measured between the 2D cursor trajectories. Despite the behavioral consistency (Figure 1a; Supplementary Figures S1-S4), population spiking activity showed significant variability: while it was fairly stable in terms of which specific neurons were activated on each electrode, their median spike counts varied across blocks (Figure 1b; Supplementary Figure S5). The shifts in the total population spike count occurred far too rapidly to be the result of physical electrode drift, which is known to be quite stable over a few consecutive days for chronically implanted Utah electrode arrays.^3^

If the mapping between individual neurons and motor output was rigidly fixed, a static linear decoder trained at the beginning of a session should maintain stable performance. Instead, we observed that static linear decoders failed in a patchy manner, with readout performance dropping in and out over the course of the session or failing to generalize all together (Figure 1c; Supplementary Figures S6-S9). This decoding failure makes sense if the underlying manifold is systematically twisting out of alignment with the static 96D decoding weights. Indeed, projecting the block-by-block trial-averaged firing rates into the baseline coordinates extracted from the first block revealed that the neural trajectories continuously rotate out of their original alignment (trajectories bounded by ellipses in Figure 1d for Monkey U, 2020-11-30). Despite the small number of trials per individual block, reasonably separable trajectories were observed relative to the trajectories obtained via low-dimensional projection of the data into the subspace determined from all the session’s trials (Figure 1e). To verify that these rotations occurred without altering the animal’s behavior, we categorized pairs of blocks based on their positions within the bisected 3D neural state space, comparing periods where the manifold remained within the same half to periods where the network had rotated into the opposing half. Comparing the pairwise Fréchet distances for these two conditions revealed no significant difference between blocks over the course of the session (Figure 1f; Supplementary Figures S1-S4). This demonstrates that whether the neural manifold remained structurally stable or underwent a large spatial rotation, the resulting kinematic output was not significantly different.

### Reparameterization of the neural state mitigates non-stationary neuron activity and preserves population dynamics

To quantify the observed reorganization at the population level, we tracked the transport of the neural manifold using principal subspace angles. For each pair of sequential blocks, we computed the transport angles for both the population state subspace (the structure of the neuron population spike count covariance) and the instantaneous population dynamics subspace (the underlying vector field). These two subspaces are bound by the continuous Lyapunov equation (Box 1); measuring them simultaneously allows us to test whether their geometric relationship remains stable over time.

These geometric shifts revealed continuous rotations of the network’s coordinate frame over the course of single experimental sessions. We visualized this reorganization using 3D spherical projections, where sequential trial blocks are mapped as trajectories along a sphere to explicitly capture both the block-to-block and cumulative angular divergence of the manifold (Figure 2a, Supplementary Figures S10, S11). Rather than clustering tightly near 0^○^ as predicted by a static manifold, the angular histograms demonstrated that the state and dynamics subspaces routinely underwent block-to-block rotations of 20 40^○^. While the manifold exhibited continuous rotations, the transport remained strictly bounded. Despite accumulating dozens of local block-to-block shifts, the cumulative transport measured from the start of the session rapidly saturated between 20 40^○^ and never approached the geometric limit of 90^○^. This saturation confirms that the population is not undergoing random drift; rather, it is exploring a bounded cone of highly degenerate coordinate frames. To evaluate whether these internal rotations exceeded estimation noise, we computed the excess mass of the block-to-block angular distributions relative to trial sequence shuffle controls (Figure 2b). This analysis confirmed significant excess mass, proving that the observed coordinate frame rotations represent genuine, bounded structural reorganization (Supplementary Figures S12, S13).

**Figure 2:**
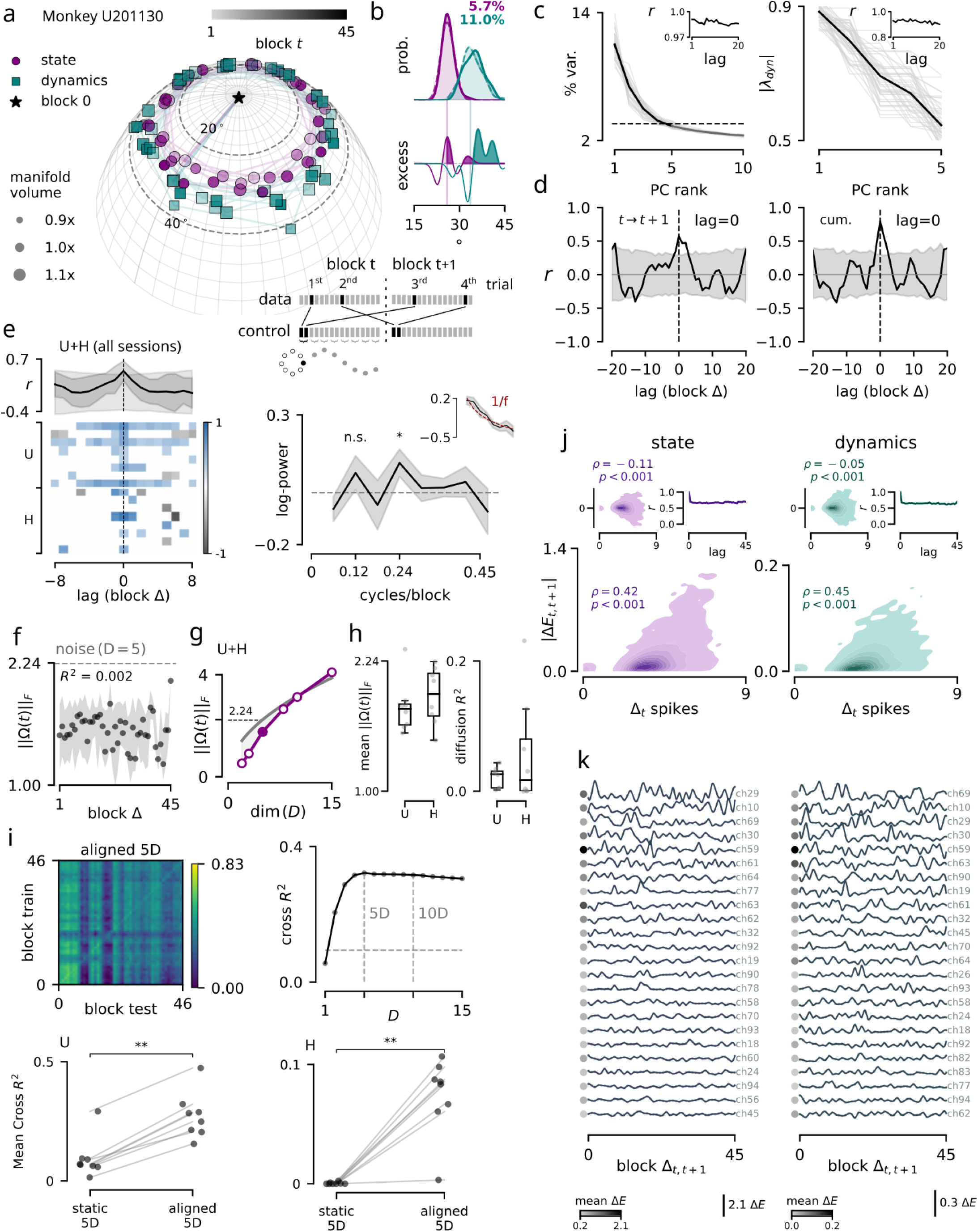
Reparameterization of the neural state mitigates non-stationary neuron activity and preserves population dynamics. **[a]** Principal subspace angles for the top five dimensions of the neural state (purple) and the linear dynamical system (teal) across example session U2020-11-30. In the spherical projection, contiguous block-to-block angular deviations are mapped azimuthally, while cumulative subspace deviations with respect to block 0 extend radially from the origin and remain bounded. Rotational transport occurred without deformation of the network geometry: the absolute manifold volume was strictly preserved (±0.05 relative to block 0) and the mean translation of the manifold in the high-dimensional space remained minimal (0.1-0.2; Supplementary Figures S10,S11). **[b]** Block-to-block subspace angle distributions for the empirical data and a trial-shuffled control. As shown in the schematic, when comparing two sequential blocks (*t* and *t* 1), successive trials were interleaved to abolish temporal structure between the blocks, better isolating finite-sample estimation noise using the exact same trial sets (see also Supplementary Figures S12,S13 for two additional shuffle controls). The empirical data exhibited significant excess mass at both extremes compared to the null: angular deviations smaller than the control reflect hyper-stability of the neural manifold (light shading), while the rightward tails reveal structural rotations that physically exceed the unconstrained noise bounds (darker shading). **[c]** State and dynamics eigenvalues remain stable across blocks as the system undergoes geometric reparameterization (*insets*: Pearson correlation between eigenvalue traces). **[d]** Cross-correlation between the rotations of the state space and the dynamics matrix revealed a highly significant peak at a temporal lag of 0, indicating synchronous geometric coupling (left, block-to-block angular deviation; right, cumulative deviation from block 0; gray shading shows 95% CI). **[e]** (*Top*) The average cross-correlation between state and dynamics transport for Monkey U and H sessions was significant at a lag of 0. (*Bottom*) Individual session cross correlations, where values less than the 95% confidence interval have been zeroed out. (*Right*) Detrended cross-spectral density revealed a significant oscillation at 0.24 cycles/block (*p* 0.05), alongside a subharmonic at 0.12 cycles/block (n.s.; *p* 0.13). The inset displays the log-power spectrum overlaid with the extracted aperiodic 1 *f* fit used in the detrending step (red). **[f]** Averaged unweighted rotational transport (Ω *t*; Equation 5, Box 1, Supplementary Figure S14) as a function of block separation, reflecting a bounded diffusion process. **[g]** Averaged temporal transport for all sessions and animals as a function of the number of retained dimensions. By 10D, the transport rate meets the null boundary √*D* (the geometric limit of orthogonalization). **[h]** Global summary of the mean (Ω) and *R*^2^ values across all sessions for Monkeys U and H. **[i]** Procrustes subspace alignment using the top 5 dimensions partially rescues the decoder from the failure observed in Figure 1c. This kinematic recovery saturates at approximately 5 dimensions (occasionally requiring up to 10; see Supplementary Figures S6-S9 for all sessions). Summary *R*^2^ values are significantly higher for Monkey U and H after 5D subspace alignment. **[j]** Single-neuron contributions to subspace energy. The absolute change in subspace energy scales with each electrode’s dynamic range (Δspikes), demonstrating that the most active neurons in a given block undergo the most extreme shifts in subspace energy in the following block. The total subspace energy was highly conserved across blocks, as evidenced by the flat autocorrelation function (insets; Supplementary Figures S15-S18). The population-wide distribution of subspace energy is robust, with the temporal autocorrelation of the population energy laying flat at ~ 0.7 (insets; Supplementary Figures S15-S18). **[k]** The top quartile of neurons exhibiting the greatest variability in energy contribution across the session. The mean-centered subspace energy for each electrode demonstrated the compensatory load-shifting among neurons that physically drive the macroscopic rotation of the manifold (scale is uniform across all rows).

Despite the animal’s consistent behavior and the stability of the population eigenspectra for state and dynamics (Figure 2c, Supplementary Figures S10,S11), the eigenvectors of the linear dynamics transition matrix shifted in lockstep with the state covariance eigenvectors (Figure 2d,e). This simultaneous preservation of the eigenspectra alongside the synchronized rotation of the spatial axes empirically verifies the conditions for continuous Lyapunov stability (Box 1). By rotating together, the network successfully preserves the global energy landscape and behavioral output despite a completely reconfigured underlying coordinate frame. To further quantify the temporal structure of these rotations, we analyzed the cross-spectral density of the transport between the state and dynamics subspaces. After detrending the aperiodic 1 *f* background component,^29^ the spectral density revealed a significant oscillatory peak at a frequency corresponding to approximately 4 trial blocks, with a non-significant but visually identified sub-harmonic at approximately 2 trial blocks (Figure 2e). This further indicates that the geometric drift is not an unconstrained random walk, but rather a process of bounded, confined diffusion with some periodic structure.

As established by the continuous Lyapunov equation (Box 1), the network must be continuously rotating its spatial eigenvectors to preserve dynamic stability. We explicitly quantified the magnitude of this structural reparameterization using the unweighted rotational generator, Ω *t* (Figure 2f, Supplementary Figure S14). This global operator acts upon the existing state covariance (*dV* Ω *t V*), such that the displacement of any given dimension is strictly proportional to its variance, forcing the highly active dominant axes to swing the furthest in physical space. Next, sweeping 1 to 15 active dimensions revealed a clear gradient of hierarchical stability (Figure 2g). From 1-5D, the transport rate is substantial but remains anchored below the orthogonal ceiling. However, by 10D, the transport rate hits the analytical null boundary (the geometric limit of complete orthogonalization, scaling proportionally with √*D*). The temporal degradation of this transport as a function of block separation (*R*^2^) is summarized for all sessions in Figure 2h and Supplementary Figure S14.

Since the observed rotational transport causes static linear readouts to fail, we applied orthogonal Procrustes alignment—a standard technique frequently utilized to correct for representational drift over much longer timescales (e.g., across days or weeks). Re-aligning the drifting active subspace back into the baseline coordinate frame significantly rescued the cross-block decoding performance. Furthermore, this rescued accuracy reliably saturated when restricted to just 5-10 dimensions, proving that despite the geometric rotation of the axes, the fundamental information necessary for kinematic control remains preserved within a low-dimensional volume (Figure 2i, Supplementary Figures S6-S9).

Having established that the population geometry continuously rotates to maintain isospectral dynamics, we sought to determine what drives this structural reorganization at the level of individual neurons. To explore this, we quantified the subspace energy (variance) of the network over time. By summing the variance-weighted squared loading weights of single electrodes for the top 5 state dimensions, we observed that individual neurons continuously waxed and waned in their energetic contributions to the active subspace, which scaled proportionally to modulation depth (Figure 2j,k, Supplementary Figures S15-S18). Although the contributions of neurons fluctuated dramatically about their average baseline, they typically did not exhibit any monotonic trend across the session. Despite the continuous internal turnover, the population-wide distribution of subspace energy was robustly maintained across session blocks (Figure 2j, inset; Supplementary Figures S15-S18). Specifically, the temporal autocorrelation of this population energy pattern was 0.7 across all block lags: the global energy landscape is neither perfectly frozen (ACF of 1.0), nor is it collapsing (ACF near 0.0). Instead, this indicates that the observed manifold rotations are a direct consequence of the network continuously redistributing its computational load— recruiting and shedding individual neurons in an activity-dependent manner without disrupting the animal’s kinematic output.

### Latent libraries in M1 neuron populations during repeated reaches

To determine whether this geometric drift represents a continuous random walk or a structured transition between discrete representational states, we applied unsupervised clustering (K-Means) to the high-dimensional state eigenvectors appearing within the top 10 ranks of each block. The results of this high-dimensional separation are visualized via three clustering principal components (cPC1-3; Figure 3a,b). Across all experimental sessions from Monkeys U and H, we observed a redundant ‘latent library’ of discrete spatial eigenvectors, typically organized into four to six distinct encoding clusters (with five being the most common; Figure 3a-d, Supplementary Figures S19,S20). To ensure these clusters were not merely artifacts of the algorithm parsing high-dimensional noise, we compared the ratio of across-cluster to within-cluster variance of the empirical state eigenvectors against an equivalent number of uniformly random orthonormal vectors sampled from the Haar distribution. The variance ratios for the empirical data were significantly larger than the random null (Figure 3b,c), confirming the existence of discrete execution channels rather than continuous unconstrained drift.

**Figure 3:**
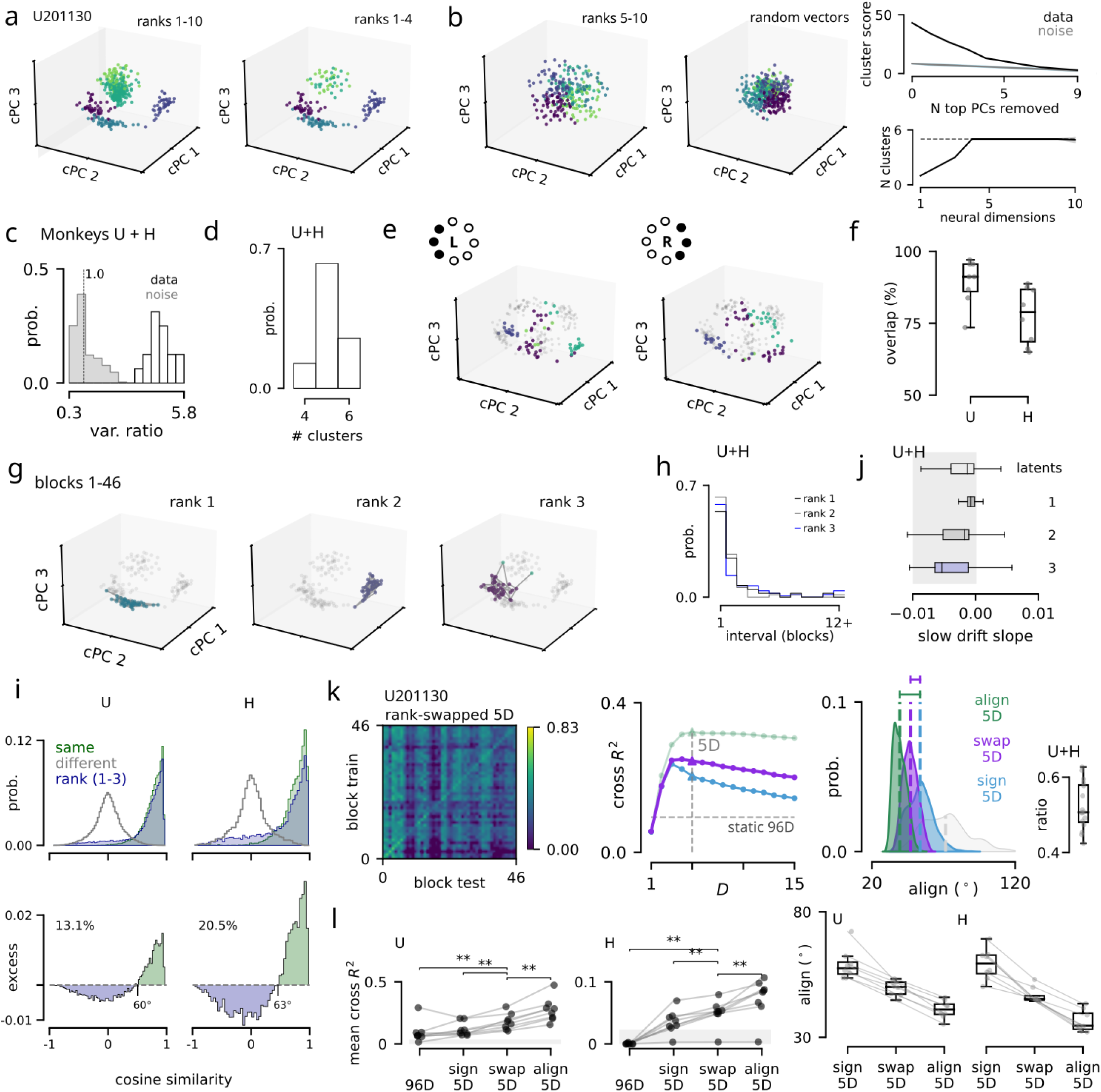
Latent libraries in M1 neuron populations during repeated reaches. For each daily session, state eigenvectors were determined independently within each block of trials. The top 10 ranked eigenvectors from each block were then pooled to form a universal set for clustering analysis. [a] Low-dimensional visualization of discrete geometric states extracted from ambient 96D eigenvectors for ranks 1–10 (*left*) and ranks 1–4 (*right*) in example session U2020-11-30 (cluster principal components, cPCs, range [-1,1]; see Supplementary Figures S19,S20 for each individual session). [b] The same analysis was repeated on sample-matched, uniformly random orthonormal matrices drawn from the Haar distribution. Progressively removing the top-ranked eigenvectors from the data and re-calculating the between– to within-cluster variance score (Calinski-Harabasz) showed that clusters start to resemble unconstrained noise by rank 5. [c] The ratio of between– to within-cluster variance of the empirical data was computed for all sessions and compared to the variance ratios derived from 1,000 realizations of the null distribution. [d] Number of discrete encoding clusters identified across all sessions. [e] Active subspaces constructed exclusively from leftward versus rightward reaching trials show largely preserved eigenvectors. Grey dots are the universal block eigenvectors determined from trials across all target directions. [f] Summary of the shared spatial occupancy (histogram intersection) of left and right eigenvectors across all sessions (see Supplementary Figures S21,S22). [g] Temporal evolution of eigenvector activation across consecutive trial blocks for example session U2020-11-30. While the cluster labels remain largely stable in this session, the underlying rank 1 eigenvectors experience continuous structural transport without breaking the macroscopic K-Means boundary, while ranks 3-5 frequently shifted clusters (see Supplementary Videos S1-16 for all sessions). [h] Summary of rank-swap intervals for all sessions, broken down by individual eigenvector rank. [i] (*Top*) Pairwise cosine similarity between state eigenvectors grouped by the same identified cluster, different identified clusters, and specific eigenvector rank (1–3). The rank-matched distribution shifts leftward, proving that distinct eigenvectors dynamically swap variance ranks over a session. Including ranks 4 and 5 led to much larger discrepancies (Supplementary Figure S23). (*Bottom*) Excess probability mass of cluster-matched versus rank-matched distributions suggested a 60^○^ spatial separation is required to maintain distinct representations. [j] Slow geometric drift was computed by measuring the cosine similarity between the latent encoding patterns, or ranks, as a function of block separation and averaging across all pairwise comparisons. The slope of continuous degradation is minimal across all datasets. [k] Kinematic decoder performance improves markedly when applying a discrete Hungarian rank-swap permutation to the top 5 dimensions. Principal deviation angles between temporal blocks are shown for continuous Procrustes alignment (align 5D), discrete Hungarian rank-swapping (swap 5D), sign-corrected PCA (sign 5D), and a scrambled rotational control (rand. 5D). The inset box plot illustrates the rotation recovery ratio, quantifying the proportion of total angular axis error that is successfully corrected by discrete rank-swapping only, relative to the sign-corrected baseline. Sign-uncorrected eigenvectors illustrate PCA’s artifactual sign-flipping error (grey). [l] Summary of cross-block decoding performance and the rotation recovery ratio for sign 5D, swap 5D, and align 5D, compared across all sessions for Monkeys U and H. The grey shading bounds the maximum observed decoder performance for the random 5D control.

We next investigated whether different kinematic behaviors (e.g., leftward versus rightward reaches) utilized exclusive, target-specific subspaces, or if they drew from a shared library of latent state dimensions. To characterize this overlap, we assigned global cluster labels to eigenvectors extracted independently from leftward versus rightward reaching trials. By computing the histogram intersection of relative cluster visits, we observed a remarkably high degree of structural overlap between conditions (Figure 3e,f, Supplementary Figures S21,S22). This confirms that the *mean-centered* neural activity does not partition into exclusive, behavior-specific attractors. While condition-dependent offsets serve to spatially translate the manifold to distinct operating points and oriented rotational dynamics,^21^ stripping these static offsets revealed that the residual dynamics themselves ultimately route through a highly conserved, shared latent library.

Having established that the dominant execution channels are drawn from a discrete latent library, we next observed that their variance ranking could swap across blocks (Figure 3g,h; Supplementary Videos S1-16). To evaluate the extent of rank swaps, the cosine similarity between identified encoding patterns in the latent library were compared pairwise both within and across clusters, as well as by population state dimension (ranks 1-3); see Supplementary Figure S23 for ranks 1-5). The choice to focus on the top 3 ranks here had the following rationale: first, by rank 5 the eigenvalues are approaching the Marchenko-Pastur theoretical limit and might experience meaningless changes in rank; second, after about 7 ranks the state’s spatial eigenvectors hit the isotropic noise floor; and third, because decoder performance saturates around 5D, which suggests lower ranks are mostly pure null space dimensions. As expected, eigenvectors belonging to the same cluster displayed a high degree of similarity, while patterns from different clusters were effectively orthogonal. However, when patterns were compared strictly by their associated rank, the distribution shifted leftward relative to the within-cluster comparisons (Figure 3i). This confirms that specific execution channels will frequently drop in variance rank, causing a discrete “rank swap” as a redundant cluster assumes the computational load. Despite these fast-timescale swaps, only small amounts of slow drift were observed across the entire session, indicating that the network reliably returns to the same stable library rather than drifting continuously away from the baseline state (Figure 3j, Supplementary Videos 1-16).

Finally, we tested whether tracking these discrete rank-swaps could functionally restore decoder accuracy. While full subspace alignment successfully rescued decoding by treating the manifold as undergoing a continuous geometric rotation, we hypothesized that a significant portion of this drift is actually driven by the abrupt hand-off of control between redundant latent states. To test this, we implemented a Hungarian-matching swap decoder that leverages the absolute cosine similarity matrix to geometrically pair the baseline state eigenvectors with another block’s eigenvectors, even if they had dropped in variance rank. This discrete rank-swapping algorithm significantly rescued decoder performance compared to a baseline sign-corrected PCA decoder (Figure 3k,l, Supplementary Figures S6-S9). While it did not fully reach the performance ceiling of the continuous 5D subspace alignment, the discrete rank-swap correction successfully accounted for approximately half of the total angular axis error for Monkeys U and H. This provides robust evidence that the motor cortex utilizes a discrete library of redundant execution channels, continuously swapping their variance hierarchy to preserve stable kinematic output.

### Discrete latent libraries in mouse ALM populations during sensory discrimination

To determine whether a fluid neural manifold represents a conserved mechanism of cortical computation rather than a primate– or task-specific artifact, we extended our analysis to a distinct mammalian dataset: open-source extracellular electrophysiology recordings from the anterior lateral motor cortex (ALM) of awake, behaving mice (*N* = 6 subjects) performing a discrete delay-response lick task.^30^ By applying our identical block-wise computational framework to this recorded population activity (Figure 4a, Supplementary Figure S24), we found that the mouse dynamics largely mirrored the primate findings. Despite fundamental differences in species, neural sampling technology, and task modality, the ALM representations exhibited similar continuous rotations in both the state and dynamics subspaces (Figure 4a-c, Supplementary Figures S25-S27), activity-dependent fluctuations in single-neuron energetic contributions (Figure 4d, Supplementary Figures S28,S29), as well as discrete rank-swapping among a flexible latent library of state eigenvectors (Figure 4e-h, Supplementary Figures S30,S31).

**Figure 4:**
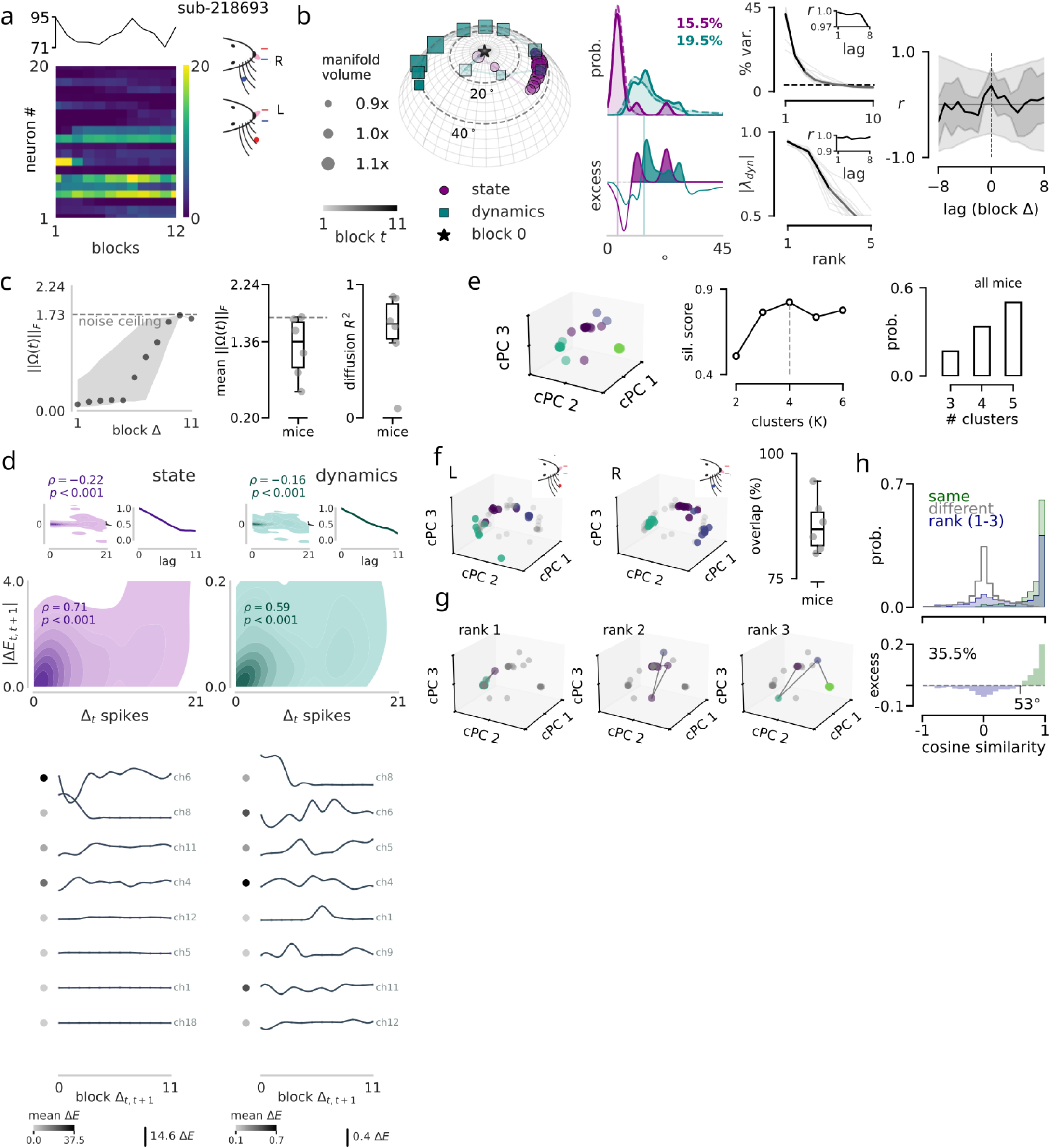
Discrete latent libraries in mouse ALM populations during sensory discrimination. [**a**] The same analyses were extended to multi-channel recordings from the anterior lateral motor cortex (ALM) of six mice performing a tactile discrimination task, which required a directional lick response for reward (Dandiset:000010).^30^ Spike counts for individual electrodes across trial blocks are shown for an example session (sub-218693). [b] Significant subspace rotations were observed for both state and dynamics eigenvectors (ranks 1-3 analyzed). Consistent with the primate motor cortex, block-to-block angular deviations exhibited significant excess probability mass at both extremes relative to the temporal shuffle control. State and dynamics eigenvalues were conserved and cross-correlation revealed a zero-lag peak in the average across mice, although it was not significant (see also Supplementary Figures S25,S26). Mean translation of the manifold in the high-dimensional space remained minimal (0.37-0.65). *R*^2^ values were variable across sessions, reflecting both steady and evolving transport rates. [c] Unweighted rotational transport (Ω *t*) as a function of block separation showed periods of confined diffusion among sharper transitions where the active subspace abruptly jumped further away from the original baseline state (block 0; see also Supplementary Figure S27). ALM transport frequently shifted into a high-rate drift that approached the analytical null boundary for complete orthogonalization (scaling with √*D*; 1.73 for ranks 1-3). [d] Single-neuron contributions to subspace energy. (*Top*) The absolute change in subspace energy scaled with each electrode’s dynamic range, confirming that highly active neurons drive the macroscopic manifold rotation. The linear decay of the autocorrelation indicates that the total subspace energy captured by the probe steadily dissipated as a function of block separation (Supplementary Figures S28,S29). (*Bottom*) Raster plot for the top quartile of ALM neurons exhibiting the greatest variability in subspace energy contribution across the session blocks. [e] State eigenvectors were clustered to form a discrete latent library containing 3 to 5 stable geometric states across all mice (Supplementary Figure S30). [f] Active subspaces constructed exclusively from leftward versus rightward lick trials show largely preserved eigenvectors. Summary of shared occupancy across all sessions (see also Supplementary Figure S31). [g] Temporal evolution of eigenvector activation across consecutive trial blocks for the example session reveals frequent rank swaps. [h] Grouped pairwise cosine similarity confirmed dynamic rank-swapping among the dominant state eigenvectors, with a geometric separation of 53^○^ required to maintain distinct representations (see also Supplementary Figure S30).

While the core computational mechanics were conserved, the distinct nature of the mouse experiment uncovered notable differences in the timecourse of the manifold’s structural drift. Specifically, both the variance weighted principal subspace angles and the unweighted Ω *t* transport (Figure 4b,c) revealed a tendency for the mouse ALM to linger longer near local dynamical solutions (evidenced by increased density near 0^○^ transport) before abruptly jumping into the 20 40^○^ rotational band characteristic of the primate data. In some sessions, these rotations exceeded 40^○^, with Ω *t* transport occasionally approaching the theoretical limit. Furthermore, unlike much of the primate data, the total subspace energy in the mouse ALM steadily dissipated as a function of block separation (Figure 4d inset; Supplementary Figures S28,S29). This dissipation likely reflects the larger unconstrained null space of a simple 1D licking task compared to a precision 2D primate reach, compounded by the much smaller neural sample size of the silicon probes. While the Utah array recordings in the primate successfully captured stable subspace energy in the local population, the subspace energy dissipation in the mouse recordings suggested that the active execution channels were frequently transported outside the limited sampling volume of the silicon probe.

### Spike-rate adaptation builds a latent library through reparameterization of the neural state

To test the hypothesis that activity-dependent fatigue acts as a causal engine for the observed manifold rotations, we constructed a biophysically plausible spiking neural network model. The network featured a balanced excitatory-inhibitory architecture^31,32^ with five distinct excitatory co-ensembles (E1-E5) configured to represent the empirical latent library. Crucially, these neurons were equipped with spike rate adaptation (SRA)—a pervasive, activity-dependent hyperpolarizing current in cortical neurons that forces highly active units to gradually reduce their firing rate over time.^33^ By driving the network with stable, low-dimensional sinusoidal target signals, we found that the inclusion of SRA was biophysically sufficient to reproduce the empirical transport dynamics and latent library observed *in vivo*. Conversely, when SRA was disabled, the network became trapped within a single structural cluster, resulting in a static, rigid manifold (Figure 5a,b). Across multiple random initializations of the network, the critical crossover boundary separating the eigenvectors was remarkably consistent, occurring at an intersection angle of 55.6^○^ 6.5^○^ (Figure 5c,d). This closely matched the 50 60^○^ observed empirically in the mouse and primate data.

**Figure 5:**
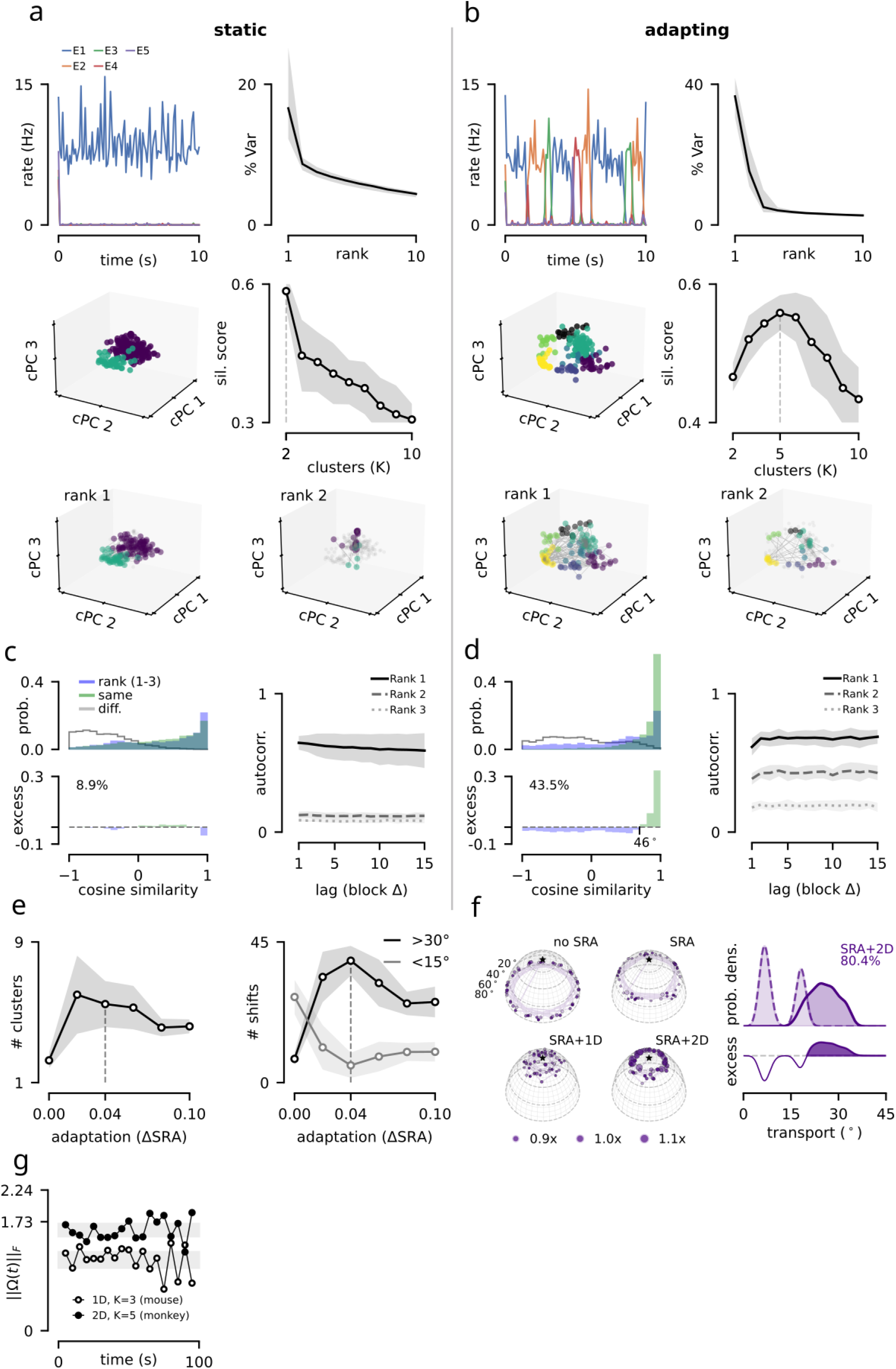
Adaptation builds a latent library through reparameterization of the neural state. **[a]** Simulations of a balanced excitatory-inhibitory spiking network that consisted of *N* = 1, 000 leaky integrate-and-fire neurons, evaluated with and without spike rate adaptation (SRA). Sparse baseline connectivity was augmented by embedding five excitatory co-ensembles (E1–E5). These co-ensembles were parameterized with varied sizes (*N* 100, 220), generating hierarchical activation probabilities and yielding an eigenspectrum that better replicates the *in vivo* data (see Methods for further detail). Without SRA, population state eigenvectors remained trapped in a static, rigid manifold. **[b]** When SRA was added, a latent library with five clusters emerged and rank swaps occurred during repetitive activation of the neuron ensembles (ranks 1-4 used for cluster visualization; activation sequences are shown for ranks 1 and 2). **[c]** Without SRA, individual rank stability far exceeds baseline within-cluster cosine similarity. **[d]** The addition of SRA to the model evoked the continuous geometric transport observed *in vivo*, generating the characteristic excess probability mass. **[e]** Sweeping the magnitude of SRA conductance. (*Left*) Biologically plausible values (~0.04) maximize the number of active clusters in the latent library. (*Right*) Operating in this same regime preferentially drives large macroscopic rotations (30^○^) over the small rotations caused by slow drift and estimation noise (15^○^). **[f]** (*Left*) With SRA active (using either a 1D or 2D continuous external stimulus), structural transport exhibits bounded diffusion. Removing SRA from the model artificially inflated block-to-block angular shifts to 80^○^, approaching the analytical null boundary for complete orthogonalization. (*Right*) A temporal shuffle control was performed across successive model blocks, yielding a distinctly bimodal distribution with peaks at 10^○^ and 20 25^○^. **[g]** Averaged unweighted rotational transport (Ω *t*) rates were reproduced simply by matching the dimensionality of the behavioral stimulus and the number of top ranks analyzed (1D/3 ranks for mice; 2D/5 ranks for non-human primates).

By sweeping the magnitude of the SRA conductance, we found that it directly influenced both the number of active clusters in the latent library and the frequency of macroscopic jumps (30^○^). Both of these values peaked within the biologically plausible range (Δ*_SRA_* 0.02, 0.05; Figure 5e; see Methods). When operating within this narrow physiological regime, the model successfully reproduced the distinct dynamics of both species by matching the dimensionality of the behavioral stimulus and species-specific adaptation strengths (Δ*_SRA_* = 0.02 for the mouse 1D lick; Δ*_SRA_* = 0.04 for the primate 2D reach; and Δ*_SRA_* 0.05 for spontaneous network transport simulations). When driving the network with a continuous 2D oscillatory drive (0.7 and 2.0 Hz), the model generated the 20 40^○^ continuous transport distribution observed in non-human primates. Applying a 1D sinusoidal stimulus (1.5Hz) reproduced the metastable local dwell times characteristic of the mouse ALM, as well as large angular transport (>20^○^; Figure 5f). The model was also able to reproduce the species-specific rotational generator magnitudes observed *in vivo* (Figure 5g).

The causal link between neuron adaptation and macroscopic rotation yielded a reinterpretation of neural variance. In standard trial-averaged analyses, continuous rotations of the manifold are often characterized as unstructured trial-to-trial noise. However, when we applied the temporal shuffle control to the model (Supplementary Figure S32), we observed a distinct bimodal distribution in the transport null distribution. This bimodality can be explained as a direct artifact of discrete cluster hand-offs: when comparing blocks that do not span a state transition, baseline estimation noise sits narrowly near 10^○^. However, when the artificial time boundaries of a block happen to straddle a change in state, the resulting temporal mixture drives the apparent ‘noise’ estimate up to the 20 25^○^ range, which matches the empirical transport observed *in vivo*. This variance is not isotropic error, as supported by the excess mass analysis across all our empirical datasets. Rather, what has traditionally been discarded as noise may actually be the signature of a structured, autonomic load-balancing mechanism.

## Discussion

When producing prolonged and repeated muscle contractions, the motor neurons of the spinal cord flexibly cycle their contributions to avoid fatigue and meet task demands.^34–37^ Our findings reveal that upstream cortical networks employ a conceptually similar strategy – rotational homeostasis – to maintain consistent motor output during repeated activation. Viewed through a classic computational framework,^38^ individual neurons serve as the biological hardware, while the discrete execution channels of a latent library act as the algorithmic engine that implement a goal-oriented computation. Rather than relying on a rigid, static mapping between the hardware and algorithmic level, the motor cortex appears to continuously shift the computational load across a redundant set of execution channels.

This dynamic cycling forces a re-evaluation of how neural stability is measured. Significant focus has been directed toward representational drift across days or months, driven by synaptic turnover, neurogenesis, slow fluctuations in excitability, and our sampling of neuron populations.^2,3,39–42^ It is well known that after affine transformation, manifolds are highly conserved and can be useful for decoding over long timescales.^3,43,44^ However, there has been comparatively little focus on reconciling these findings with the fast-timescale fluctuations of individual neuron responses^12,13,16^ and their task-driven covariance.^20^ Much of the literature studying cortical dynamics achieves a highly consistent estimate of an averaged neural manifold by pooling activity over hundreds of repeated trials. However, what is gained in statistical stability is lost in temporal resolution; this approach inherently blurs away the fast-timescale, non-stationary changes in correlated population activity. Recent work in the hippocampus demonstrates that slow drift over time is governed by distinct mechanisms compared to the fast drift evoked by rapid iterations of a behavior;^14,45^ our results firmly establish a similar behavior-driven algorithmic drift in the motor cortex, decomposable into continuous and discrete transport.

If this faster timescale drift exists at the algorithmic level, what biological mechanisms are responsible? Our analysis of subspace energy revealed that individual neurons continuously sub in and out to maintain the population state. As demonstrated by the spiking neural network model, activity-dependent spike rate adaptation (SRA) provides a sufficient causal driver of this continuous population transport. Furthermore, theoretical work demonstrates that cortical networks with balanced excitation and inhibition naturally produce metastable transitions between structured clusters of active neurons,^31,32^ and that rapid, sub-second shifts in this balance can dynamically compensate for localized neuron loss.^46^ It is highly probable that the discrete switches among the latent library are heavily regulated by similar fast-timescale shifts in global inhibitory scaling, effectively connecting localized cellular mechanisms to abstract computations performed at the population level. In fact, recent computational modeling has demonstrated that such abrupt, discrete jumps in neural representations are mathematically easier for downstream circuits to stably decode than slow, continuous diffusion.^47^ The rotational homeostasis observed here provides a concrete biophysical mechanism for this phenomenon: activity-dependent fatigue eventually forces the network to execute abrupt, discrete rank-swaps to preserve downstream kinematic readouts.

This physiological cycling provides a substrate for theories of redundant motor learning.^17^ Seminal brain-computer interface experiments^48,49^ have demonstrated that learning novel neuron-to-latent mappings (OFF manifold) is slow, whereas animals can quickly learn new associations within a few hundred trials, provided the new solution lies within the existing intrinsic manifold (ON manifold).^48^ The rotational homeostasis we observe operates over similar timescales. Furthermore, ON manifold adaptation relies on the flexible re-association of existing latent patterns to new behavioral conditions.^49^ Our observation that latent library patterns broadly overlap for both left and right target sets provides evidence for this principle during natural behavior. While our block-wise K-Means may coarse-grain over finer, target-specific cluster structure, the equivalence is clear. The motor cortex does not generate entirely novel subspaces (OFF manifold) to execute distinct reaches or combat fatigue; instead, it avoids interference by continuously re-associating a degenerate reservoir of ON manifold latents that already have established mappings to behavior. By restricting this internal drift to the intrinsic manifold, downstream networks can maintain a stable behavioral readout—a decoding mechanism supported by recent computational models^47,50^

In the non-human primate datasets, action potentials were identified by thresholding high-pass filtered voltage traces; as such, the spike trains recorded on some electrodes represent an informative but unresolvable multi-unit hash.^51^ Looking forward, fully resolving continuous and discrete transport dynamics will require transitioning from multi-neuron pooling, which inherently obscures the multi-phasic activity of individual neurons,^51^ to chronic high-density silicon probes capable of precisely identifying individual spiking responses.^52–54^ Furthermore, the validation of these results in the rodent ALM opens the door for targeted optogenetic perturbations^30^ and specific cell-type identification.^55^ Ultimately, anchoring these drifting neural states directly to continuous behavior will be essential to decipher how the brain maps dynamic neural circuits to consistent real-time action.^56,57^

### Box 1: Covariance decomposition in linear dynamical systems

Consider a linear stochastic dynamical system of the form

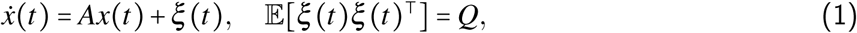

where *A* defines the recurrent dynamics and *Q* defines the covariance of external and/or intrinsic fluctuations. Models of this form are commonly used to model neuronal dynamics from recorded populations of neurons.^58^ We do not discuss nonlinear models – linear models have been used extensively in the field and are the basis of the methods used here to detect reparameterizations of the neural activity in state space. The following discussion generalizes to non-autonomous input provided it’s bounded.^59^

In the stationary regime, the covariance matrix of this dynamical system, Σ = Cov (*x*), satisfies the continuous Lyapunov equation:

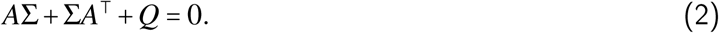

This classic equation, with applications in neuroscience,^60^ establishes that Σ is not an independent object, but is fully determined by the interaction between dynamics (*A*) and the structure of the system noise and input (*Q*). It is expected that the noise structure and its covariance Q is static, especially for the heavily practiced behavior and short-timescales considered here. Thus, it’s not considered a principal contributor to changes in the high-dimensional spike count covariance. Under these assumptions, changes in Σ can occur from two distinct mechanisms.

**Mechanism 1** Changes in the recurrent dynamics, *A*→*A*^′^, modify the propagator *e^At^*, which governs how activity evolves over time. Since

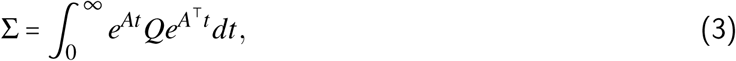

changes in *A* alter how noise and external input are filtered through the system. Mechanistically, this corresponds to: (i) changes in stability (real parts of eigenvalues of *A*), (ii) changes in oscillatory structure (imaginary parts of *A*), and (iii) the re-organization of dynamical modes. The latter necessitates the rotational transformations performed by algorithms such as canonical correlation analysis to recover consistent manifolds across different recording sessions, different animals, and even distinct behaviors.^3,61,62^

Changes in the underlying dynamics matrix *A* could manifest in the spatial covariance Σ as either shifts in eigenvalues (variance redistribution) or structured rotations of the state eigenvectors to align with the new dynamical modes of *A*^′^. The eigenvalues (*λ_dyn_*) dictate the temporal decay rates and frequencies of the macroscopic movement; these must remain stable across the session to preserve the behavior. Along with certain changes to the eigenvectors of *A*, this is the only class of change that corresponds to an actual modification of the underlying *computation* and would likely only occur under conditions of learning a new skill or re-learning after perturbation (e.g., physical injury, neuron loss, or electrical/pharmacological interventions). However, the eigenvectors of *A* could change if confined within a degenerate equivalence class, where effective macroscopic dynamics and behavior remain invariant (isospectral flow).

**Mechanism 2** Representation-driven changes through invertible changes of coordinate embeddings in the neuron population.

Consider 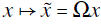, which induces the transformations,

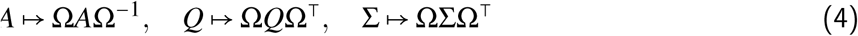

These transformations do not affect physical dynamics or behavior but change the observed representation of state space. In particular, as eigenvectors of Σ rotate, eigenvalues remain invariant provided Ω is orthogonal (e.g., permutation, reflection and rotation matrices). This corresponds to switching within an equivalence class of a behavior-preserving quotient space.

Combining the above, the temporal evolution of covariance in the linear dynamical system can be expressed as:

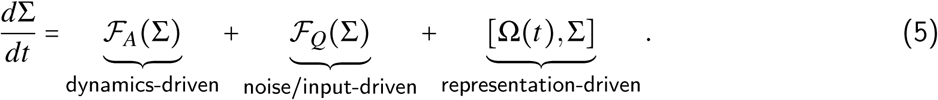

While their form is unknown, *F_A_* captures hypothetical changes in covariance induced by modifications in *A*, *F_Q_*captures changes induced by modifications in *Q*, and [Ω, Σ] denotes reparameterizations of the state space observed in our experimental data. Under conditions of a heavily practiced, stereotyped behavior, the rate of change of covariance with respect to isospectral population dynamics and input driven terms should be near 0, leaving representation as the dominant source of changes observed in the state space eigenvectors as the system reparametrizes itself in the high-dimensional neuron population. In theory, this invertible transformation (*A→*Ω*A*Ω^−1^ and Σ→ΩΣΩ^⊺^) guarantees that drift is mathematically isospectral. The spatial hardware (Σ) and the algorithmic engine (*A*) are rotated in lockstep by Ω(*t*) to cycle fresh neurons into the potent space without altering the underlying behavioral computations. For further detail about the connection between high-dimensional covariance and low-dimensional covariance with respect to the above Lyapunov stability analysis, see Methods.

## Detailed Methods

### Experimental Design

Two adult male rhesus macaques, Monkeys U (11 years of age) and H (14), were presented with a center-out reaching task during recordings from primary motor cortex using 96-electrode Utah arrays (Blackrock Microsystems, Salt Lake City, Utah, USA). Monkey U was implanted in 2017 with three arrays that had 1.0mm electrodes coated in iridium oxide; one placed in the left medial primary motor cortex (not used for data collection), the left lateral primary motor cortex (data reported here), as well as left premotor cortex (data not reported here). Monkey H was implanted in 2012 with arrays that had 1.0mm platinum coated electrodes in the right primary motor cortex (data reported here) and premotor cortex (data not reported here). He performed reaches with his left arm and the primary motor cortex array was used for the lesion. Importantly, all Utah arrays utilized in this study were chronically implanted for several years prior to data collection. This extensive multi-year timeline ensured that any acute mechanical instability, tissue encapsulation, or initial hardware drift had long since reached a stable equilibrium.^39,63^ Following the initial four consecutive days of data collection for Monkeys U and H, micro-scale electrolytic lesions were performed through adjacent electrodes. A subsequent set of four recording sessions was then gathered one month later, well after the initial post-lesion recovery period (approximately 2 days). Notably, the delivery of this highly localized electrolytic current did not disrupt the structural integrity of the hardware, nor did it impair the capacity to record robust neural population activity across the array during these later sessions.^64^ All animal procedures were reviewed and approved by Stanford University’s Institutional Animal Care and Use Committee.

Monkeys performed a simple reaching task during daily recording sessions from motor cortex over the span of four to five days to minimize any potential changes in the animals’ behavior and associated neural data streams. The animals were highly engaged and limited to 750 successful trials, ensuring minimal confounds from fatigue or waning interest, which can occur after initial bouts of focused play. Participation in the reaching task was entirely voluntary: when the animals actively engaged with the task, the room lights were automatically dimmed and the task began. If presented with continued inactivity from the subject, the task would automatically time out, remove the target, and turn the lights back on. The task re-presented itself according to a variable schedule, offering animals subsequent opportunities to play. If multiple re-presentation cycles elapsed without any play, the monkey was returned home. To track reach kinematics, a reflective bead was comfortably secured to the third and fourth digits of the hand (proximal phalanges) and optically tracked at 60Hz in three-dimensions with infrared cameras (Northern Digital Inc., Waterloo, Canada). The bead’s position was projected onto a computer screen as a task cursor and aligned in a virtual work space so that when the animal extended its arm straight out, the cursor mapped to the origin (0,0). This ensured centered visual alignment without having to reach across the body, which can hamper reach performance in that direction. The task also enforced a maximum permitted distance along the axis orthogonal to the screen’s surface, requiring the monkeys to maintain an extended reach and perform stereotyped movements within a defined work space of approximately 0.25m^3^. Failure to do so on a few consecutive trials would lead to a task lockout and subsequent representation a few minutes later. This helped ensure that consistent trials occurred in the largest possible continuous batches. A single reach trial was structured as follows: a green target (12mm diameter) appeared at the origin of the workspace (0,0) and the monkey moved its arm to place the cursor over the target, holding it within target bounds for 350ms (Monkey H), 250ms (Monkey U). Next, one of eight radial targets appeared at a fixed interval of *π* 4 with a radius of 100mm. Monkeys then reacted to the target’s appearance and moved their arm to guide the cursor accordingly. Successful navigation of the cursor to the target and holding it within target-bounds earned a juice reward. The target then reset to the center and the cycle was repeated for hundreds of trials at the animal’s discretion. Note, sessions with Monkey H omitted the bottom and bottom-left targets due to an apparent visual field defect that presented during initial task training. In this case, the block size was two repeated trials for six different targets or 12 trials instead of 16 (Monkey U; Figure 1a).

To avoid potential prediction of target location and ensure robust, unbiased temporal comparisons across blocks, the experimental task presented target direction pseudo-randomly; only after each target in the set had been successfully reached would the task move on to the next set of trial reach directions. As an additional safe-guard, trial blocks were uniformly balanced by stratifying them into balanced blocks at the analysis stage – here, a block was strictly defined as the shortest chronological sequence containing exactly two successful reach trials for every unique target direction. If there were any irregular trial sequences that arose due to an interrupted experiment or any rejection of trials during pre-processing (e.g., noise contamination), this step would correctly balance the block. Although the task was designed to present balanced blocks in the first place, this additional programmatic stratification guarantees that all latent temporal comparisons are perfectly balanced for reach kinematics and are not confounded by uneven target sampling.

Data streams and control signals for the infrared camera, experimental rig lights, juice reward system, and the behavioral task were collected, synchronized, and controlled at 1kHz using the Brain Interfacing Lab’s open-source platform for realtime processing, LiCoRICE.^65^ Behavioral measures presented in this study were obtained from the time varying two-dimensional task cursor position data, differenced and smoothed using a Savitzky-Golay filter of order 2 and window length 100ms in order to estimate the second temporal derivative, cursor acceleration. During individual reach trials (Figure 1a), the derived acceleration profile was used to identify reaction time as the last positive acceleration event that occurred after the appearance of the radial target and preceded a large movement. The maximum velocity was extracted from the period spanning the reaction time to when the cursor reached the target.

### Behavioral Consistency

To ensure temporal drift in the neural manifold represented genuine internal reorganization rather than simple behavioral degradation (e.g., the subject adopting different reach kinematics over time due to fatigue), behavioral consistency was quantified across trial blocks. For each target direction, continuous 2D cursor trajectories (*x, y*) were isolated from a fixed window [-50, 400]ms relative to the trial’s reaction time. The condition-averaged kinematic trajectory for each target’s two trials were then computed within each block.

The structural similarity between trajectories from blocks (*P*) and (*Q*) was evaluated using the discrete Fréchet distance:

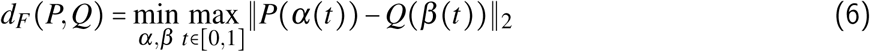

where *α*(*t*)and *β*(*t*) represent non-decreasing monotonic reparameterizations of the trajectory sequences, optimized for the data (Eiter and Mannila, 1994). This ensures that the metric is strictly sensitive to the spatial path of the reach while remaining invariant to exact velocity fluctuations within a given trial. Fréchet distances between contiguous trial blocks (*P* and *Q P* 1; Figure 1, contig.) were compared against a temporally-shuffled null distribution where *Q* is any other block randomly selected from the session. Statistical significance between the contiguous block-to-block drift and this null distribution was assessed using a two-sample Kolmogorov-Smirnov test. This ensured that neural reorganization across the blocks could not be trivially attributed to changing kinematics. An additional visual control (rand.) was performed by randomly selecting target-matched trials from across the entire session, independent of their block membership.

To directly assess whether shifts in the neuron population correlated with shifts in behavior, the empirical Fréchet distance across discrete manifold boundaries was evaluated. The sequence of blocks was subdivided into two distinct groups based on the structure of the neural manifold (determined via *K*-means clustering into two discrete states on the collective top three eigenvectors; more detail below). Block pairs were categorized as either *intra-cluster*, where both blocks possessed more similar eigenbases, or *cross-cluster*, where the blocks were associated with more distinct eigenbases. Pairwise group comparisons of the trial blocks were performed as a function of the block lag and linear regression was applied (Supplementary Figure S1-S4).

### Neuronal Data Collection and Pre-processing

Neuronal activity was sampled at 30kHz in primary motor cortex using 96-electrode Utah arrays (Blackrock Microsystems, Salt Lake City, Utah, USA). For Monkeys U and H, a Blackrock CerePlex E headstage was connected to the array’s external CerePort to transmit a digitized signal to a Blackrock Digital Hub, which relays the signal over fiber optic cable to a Blackrock Neural Signal Processor (firmware version 6.05.02.00). The NSP transmits neural data over Ethernet as UDP packets. The recorded data was then saved to a local computer, backed up to the cloud, and analyzed offline. The recorded 30kHz signals were high-pass filtered using a fourth-order zero-phase Butterworth filter with a 250Hz cutoff frequency. The root mean square voltage was computed for each electrodes’ high-pass filtered data from different time points during the session, and, if any of these RMS blocks displayed values greater than 40µV, they were discarded as noisy or corrupted channels. Action potentials were identified by thresholding the filtered membrane potential at 4 the root mean square voltage for Monkey H and –3.5× for Monkey U, measured over the first minute of each day’s recording session. The binary spike trains were determined from identified crossings of the threshold by the membrane potential. Data snippets were generated to analyze the action potential waveforms by taking the 16 sampled data points (0.53ms) before each identified threshold crossing and the 32 sampled points (1.07ms) afterward. Waveforms whose peak-to-trough amplitude exceeded 300µV are atypical of extracellular recordings from cortical neurons and were excluded as noise.^39^ The width of the waveform was calculated as the difference between the two prominent peaks in the temporal derivative of the extracellular potential. Waveforms whose widths were longer than 1ms were also excluded as non-physiological for fast– and regular-spiking cortical neurons.^66^ Next, each electrode’s action potential waveform snippets were pooled and principal component analysis was performed to identify any remaining potential artifacts present in the recordings. All binary spike rasters were then downsampled to a 1kHz raster and aligned to the behavioral data using a serial timestamp issued every 1ms by LiCoRICE and the 30kHz timestamp issued from the Neural Signal Processor (Blackrock Microsystems). This approach corrects for any potential temporal slip arising from the two data streams with their independent system clocks.

### Neuronal Data Analysis: Dimensionality Reduction

The neuron population activity analyzed in this article was aligned with the peri-movement component of the reach (Figure 1a,b). For each successful trial, the reaction time was identified from the behavioral cursor data and the spikes for each electrode were counted in the time interval 50ms before this value and 400ms after (Figure 1b; Supplemental Figures S1-3). For each trial, binary spike rasters in this 450ms reach-associated window were partitioned into eighteen non-overlapping temporal bins (*t_bin_* = 25ms) and summed to generate a raw spike count matrix. Notably, to preserve the high-frequency dynamics, the spike trains were not subjected to Gaussian smoothing or continuous rate estimation. The binned spike counts were then concatenated across all trials within the block to form a *T ×N* matrix, where *T* is the total number of temporal bins and *N* is the number of recorded electrodes. Depending on the analysis step, *T* = *n_trial_* ⋅ *n_bin_* uses just one block’s worth of trials or all the trials from a single session.

To pre-process the data for PCA, the activity of each electrode was mean-centered by subtracting its average spike count across all concatenated time bins within the trial block. Unlike standard scaling procedures, the electrode activities were explicitly not scaled to unit variance. This preserves the natural amplitude differences between high-firing and low-firing units, ensuring that the resulting covariance matrix accurately reflects the magnitude of co-modulation in the network. PCA was then applied directly to this mean-centered, un-smoothed spike count matrix to extract the spatial eigenvectors (loading weights), and the resulting eigenvalues were expressed as a percentage of the total population variance explained. The 48 largest eigenvalues and associated eigenvectors were kept for subsequent analysis, and the smallest 48 were dropped as a standard de-noising step. As mentioned above, recording sessions were split into contiguous blocks of 16 trials for Monkey U and blocks of 12 trials for Monkey H. These group sizes were selected as a minimum block size that contained two trials for each target direction, yielding reliable results from PCA and sensible power law fits to block eigenspectra. The all trial session decomposition, typical of many studies, was used as a means to resolve artifactual sign ambiguities when studying the eigenvectors of the population state and dynamics within individual session blocks (Supplementary Figure S23). Any eigen-decomposition solver carries a mathematical ±1 sign ambiguity. We evaluated the dot product of each local block eigenvector against the closest global true-north reference eigenvector computed from the entire session. If the dot product was negative, the block-level eigenvector was multiplied by 1. This locked the orientation and guaranteed that transport measurements captured real structural drift free of sign-flipping artifacts.

For illustration, trial-averaged, target-aligned population firing rates for the example session (Monkey U 2020-11-30) were projected into the fixed, static 3D PCA subspace defined by all trials in the session (Figure 1e). Next, individual block activity was projected into the static 3D subspace defined by just the first chronological block of trials (block 0). To quantify the position of the 8-target reach activity within this fixed reference frame, we computed the 2D elliptical plane of best fit for each set of 3D trajectories using singular value decomposition (SVD). These 2D elliptical planes were then plotted sequentially for example blocks to show the progressive structural drift of the state space, verifying that the internal manifold systematically rotated in-and-out of the original block’s execution channels (Figure 1e,f).

*Neuronal Data Analysis: State Eigenvector Clustering* Rather than continuously diffusing through the high-dimensional ambient space, we hypothesized that the network organizes into a finite vocabulary of discrete execution channels. To map this latent library, we pooled the spatial eigenvectors from every individual trial block across the session. We then clustered these high-dimensional encoding vectors using K-Means.

To prevent artificial clustering artifacts, the clustering was performed strictly on the raw 96D ambient neural weights, purposefully avoiding pre-mature PCA dimensionality reduction. This guarantees that the resulting discrete library states represent genuinely orthogonal, functionally distinct spatial axes in the cortex. The optimal number of discrete states (*k*, typically 4-6) was determined objectively by identifying the inflection point (“elbow”) of the K-Means inertia curve (sum of squared distances). By analyzing the frequency distribution of these assigned cluster labels for specific behavioral sub-tasks (e.g., left vs. right reaches), we quantified the degree of representational overlap using histogram intersection.

Each population state dimension within each block of trials was then assigned a label based on the clustering analysis. From here, switches between the labeled encoding patterns were determined across blocks for the largest five state dimensions. When assessing the cosine similarity between encoding patterns for both fast and slow timescales, the same low-dimensional encoding patterns were compared based on which the cluster labels were derived. Note, only the largest three to five population state dimensions were included in the analysis of cosine similarity. Inclusion of ten state dimensions resulted in a large shift of mass to be centered around a cosine similarity of 0 when patterns were compared by rank. This could artificially inflate estimates of switching, further suggesting that beyond rank 5, state dimensions may be weakly task constrained or effectively random.

To determine if different kinematic sub-tasks rely on a shared structural architecture, we analyzed the overlap of the latent library between distinct behavioral targets (e.g., leftward versus rightward reaches). K-Means clustering was first fit universally across the entire session to define the global basis set. We then assigned global cluster labels to the block-by-block eigenvectors extracted independently from each specific target group. The structural overlap between the sub-tasks was quantified by calculating the histogram intersection of the relative frequency distributions of cluster visits, measuring the proportional utilization of shared spatial motifs between the distinct behaviors.

### Neuronal Data Analysis: State and Dynamics Transport

We quantified the rotational transport of the neural manifold across blocks using two parallel geometric definitions: state subspaces and dynamics subspaces (Box 1). To extract the state subspace, we applied standard PCA directly to the concatenated single-trial spike counts of each block. The top *K* principal components represent the static spatial covariance axes capturing the majority of the population variance. To prevent the inclusion of low-variance isotropic noise, the boundary of this active signal subspace was analytically derived using the Marchenko-Pastur distribution. By adjusting the theoretical upper bound for the specific finite sampling ratios (*γ* = *N*_neuron√s_/*N*_samples_) for the block trial counts, we computed the exact eigenvalue noise floor limit 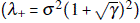. Only spatial eigenvectors possessing eigenvalues at or above the M-P threshold were included in downstream transport analyses (approximately 5 across all sessions; Supplementary Figures S10-S11).

To extract the dynamics subspace, we sought to identify the functional execution channels driving the temporal evolution of the reach. In this case, to minimize single-trial noise for the dynamical fit, spike counts were first trial-averaged across identical targets within the same block, yielding less noisy target-specific population spike counts. We then projected these block-level responses into an intermediate, mildly compressed 30D PCA subspace. This intermediate projection compresses the rank-deficient 96D ambient space to ensure the subsequent dynamical system is mathematically well-conditioned. We then fit an unconstrained discrete-time linear dynamical system (LDS), whose transition matrix **A** (*X_t_*_+1_ = **A***X_t_*) was calculated using L2-regularized Ridge regression (*α* = 1.0). We opted for L2 regularization on the full block data rather than cross-validation, as the latter is more susceptible to sampling volatility.^67^ By forcing an analytical constraint on the best fit weights, the ridge penalty prevents the dynamical system from overfitting the noise.

The fitted transition matrix **A** captures rotational dynamics, yielding complex conjugate pairs of eigenvalues and eigenvectors. However, complex eigenvectors only describe abstract rotational phases. To express these dynamics as a physical neuron-to-latent relationship, we extracted the real and imaginary components of the dominant eigenvectors and orthogonalized them. This converts the abstract complex vectors into a sensible, real-valued spatial basis where individual neuron contributions could be meaningfully assessed. Since orthogonalization preserves the hyperplanes of the original subspace, we could then reliably quantify the macroscopic transport of the dynamics bases across blocks using the subspace alignment index. Note, because subspace metrics inherently track hyperplanes rather than directed vectors, this calculation is immune to sign-flip ambiguities. Consequently, true-north sign resolution was strictly reserved for tracking the explicit spatial embedding of individual state eigenvectors (Supplementary Figure S23).

### Neuronal Data Analysis: Cross-Correlation of State and Dynamics Transport

For each consecutive block step, the scalar angular transport magnitudes for both the state subspace and dynamics subspace were extracted. We computed the Pearson cross-correlation between these two temporal sequences across a range of block lags for each session (up to a maximum lag of 20 blocks for sessions with as many as 750 trials). To establish strict statistical significance bounds, we generated 95% confidence intervals using non-parametric permutation testing (*N* = 500 permutations), systematically shuffling the transport time-series. For each session, we measured the cross-correlation on both the instantaneous block-to-block rotational shifts and cumulative rotational shifts from block 0. For the aggregate analysis across subjects in Figures 2 and 4, we focused specifically on the block-to-block angular deviations. By examining the temporal lag that maximized this cross-correlation, the analysis allowed us to verify whether the changes in the population spike count covariance (state) moved in lockstep with the functional dynamics at a zero-lag delay as expected for a Lyapunov-stable dynamical system (Box 1). Finally, we applied the Wiener-Khinchin theorem, computing the fast Fourier transform of the cross-correlation sequence to extract the cross-spectral density. We then detrended the 1 *f* component of the power spectrum^29^ to extract any dominant periodic modes of manifold rotation from the aperiodic continuous background transport.

### Neuronal Data Analysis: Ω(t)Transport

While the state and dynamics transport described above utilized variance-weighted alignment to track the dominant subspace dimensions, we quantified the total angular transport of the manifold using the unweighted Frobenius norm of the block-to-block change in the state eigenvectors (*dV* Ω(*t*)*V*). This empirical measure corresponds to the rotational generator predicted by the continuous Lyapunov equation under conditions of stereotyped behavior (Box 1). Thus, tracking the unweighted Ω-transport provides a direct, empirical measurement of the *isospectral* rotation required to preserve stable network dynamics despite continuous physiological changes at the single-neuron level.

To benchmark the rate of this transport, the empirical Ω was compared to a geometric limit of transport if two network states shared no structure. An exhaustive null boundary was analytically derived by calculating the theoretical limit of the Frobenius norm (√*D*) for the rotation generator between completely uncoupled orthogonal subspaces in the ambient space. By comparing empirical Ω-transport against this boundary, we quantified how closely the continuous reparameterization of the neural manifold approached total geometric orthogonalization over the course of the session.

Next, to evaluate how rotational transport scaled hierarchically, we analyzed it as a function of the active subspace dimensionality, *D*. At each dimensional step, we recalculated the block-to-block Ω-transport via the Frobenius norm of the respective *D*-dimensional rotation generator. By tracking the dimension at which the empirical Ω-transport rate intersected the null, the sweep was designed to identify the critical dimensionality threshold where the manifold’s continuous reparameterization ultimately becomes dominated by noisy rotation.

### Neuronal Data Analysis: Single Neuron Modulation and Subspace Energy ΔE

To directly link the geometric rotation of the manifold to the underlying biophysical fatigue of the neural population, we quantified the subspace energy (variance) for each block of trials. Since we strictly preserved the amplitude of the firing rate variance during pre-processing, the absolute magnitude of the active *K*-dimensional spatial eigenvectors serves as a direct, physical proxy for the accumulated spike rate adaptation (SRA) within the local neuron population.

For each electrode, we computed its subspace energy by extracting its spatial loading weight for each of the top *K* principal components, squaring it, scaling it by the corresponding eigenvalue (variance explained), and summing these “energy” contributions across the entire *K*-dimensional execution channel. We then tracked the energetic drops (Δ*E*) across consecutive blocks for the top 25% most energetically modulated units in the population. By analyzing Δ*E* across blocks *t* and *t* + 1, as a function of neurons’ corresponding spike count modulation depth in block *t*, we could map the structural rotation of the manifold to the shedding or accepting of activation load. When a dominant execution channel rotates, the subspace energy is forcibly redistributed; this allows the most exhausted, highly-weighted neurons to dramatically drop their modulation depth (Δ*E* 0), while fresh neurons are simultaneously recruited to preserve the stable kinematic readout.

### Neuronal Data Analysis: Cross-Block Kinematic Decoder

Block-specific memoryless linear decoders were trained to predict 2D cursor kinematics (X and Y velocities) from the instantaneous 25 ms binned neural state. The kinematic target variables for the decoder were computed as raw, non-smoothed finite differences of the cursor position. Smoothing the kinematic trajectories was deliberately avoided in this analysis, since it artificially inflates *R*^2^ performance metrics, supporting the claim that the neural population genuinely encodes instantaneous velocity on a moment-to-moment basis. A biologically plausible 100 ms sensorimotor lag was applied, such that neural activity at time *t* predicted kinematics at time *t* + 100 ms. The baseline decoder was trained exclusively on block *t* using L2-regularized ridge regression (*α* = 1.0) to predict velocity from the full 96D neural state. This static 96D decoder was then tested iteratively on all other blocks to measure the uncorrected degradation of readout accuracy (*R*^2^).

To isolate how much information was preserved within the drifting execution channels, we restricted the decoder to a 5-dimensional active subspace. The “align 5D” decoder applied orthogonal Procrustes alignment to optimally rotate the test block’s 5D subspace back into the training block’s frame before decoding. To test the hypothesis that the functional axes swap ranks, we introduced a Hungarian-matching “swap 5D” decoder. Using the scipy.optimize.linear_sum_assignment algorithm on the absolute cosine similarity matrix, the swap decoder geometrically matched the five baseline eigenvectors to the five newly rotated eigenvectors, even if they had dropped in variance rank. The decoder was allowed to search the ambient state space up to an encompassing 15 ranks deep to hunt for eigenvectors that may have dropped entirely out of the top 5D active subspace. This was particularly useful when sweeping through D=1-5 of the active subspace.

Finally, the aligned and swap decoders were compared to a sign-corrected 5D PCA decoder that controlled for trivial axis-flipping. To control for trivial, algorithmically induced axis inversions, we applied the same true north sign resolution function. This mathematically prevents artificial decoder collapse stemming from simple sign inversion between the train and test manifolds. The *recovery ratio* was geometrically quantified as the fractional reduction in angular axis error achieved by the swap decoder, measured relative to the sign-corrected baseline and the optimal geometric lower-bound of the strictly aligned (Procrustes) decoder.

To establish a statistical floor for cross-block performance, we also introduced a random rotation control. For this control, the test block was rotated using a scrambled alignment matrix (**R**_rand_ = **QRQ***^T^*), where **R** is the optimal Procrustes matrix determined from the data and **Q** is a random orthonormal matrix sampled from the Haar measure. This provided a conservative null-distribution representing the expected decoding accuracy of a completely random, volume-preserving 5D subspace rotation.

### Mouse Electrophysiology and Analysis

To validate whether the results in primate reflect a conserved cortical mechanism, the entire analysis pipeline was replicated on electrophysiological recordings from the Anterior Lateral Motor cortex (ALM) of awake, behaving mice using extracellular multi-electrode silicon probes (*N* = 6 subjects; DANDI Archive ID: 000010).^30^ Mice performed a simple 1D sensorimotor licking task with two conditions rather than a continuous 2D reaching task with 8 conditions. While the core mathematical analysis and controls remained identical to the primate analysis, a few specific pre-processing parameters were adapted to account for differences in species, neuron population sampling, and task structure.

First, free-licking in mice is inherently more variable and less kinematically constrained than an expert targeted reach; therefore, our mouse analysis focuses primarily on the geometric stability of the neural representations rather than enforcing the stringent trial-by-trial kinematic consistency controls applied to the primate data. Since the mouse task involved only two discrete targets (left and right) compared to the primate’s eight, trial blocks were strictly fixed at 16 trials, deliberately stratified to contain exactly 8 left and 8 right trials. This guaranteed perfect spatial balancing for the target-averaged responses. The recorded spikes were binned at 25 ms, identical to the non-human primate pipeline. PCA was performed on the flattened, target-averaged block spike counts, and LDS transition matrices were fit to a 15-dimensional active subspace instead of 30.

A key distinction in the mouse pipeline occurred for the state eigenvector clustering. While clustering in the primate data operated on the high-dimensional 96D ambient coordinate space, the smaller recording samples and distinct noise profile of the mouse recordings benefited from intermediate dimensionality reduction. Pooled block eigenvectors were first projected into a 3D subspace (cPC1-3), and K-Means clustering was swept from *K* 2 to *K* 6 directly within this subspace, consistently revealing 3 to 5 discrete spatial motifs.

### Spiking Neural Network Model

The model network consisted of *N* = 1000 leaky integrate-and-fire (LIF) neurons, partitioned into an excitatory population (*N_e_* = 800) and an inhibitory population (*N_i_* = 200). The integration time step was set to *dt* = 1.0 ms, with membrane time constants *τ_e_* = 15.0 ms and *τ_i_*= 10.0 ms, and a refractory period of *t*_ref_ = 5.0 ms.

To replicate the latent library observed in the empirical non-human primate and mouse data, the excitatory population was anatomically partitioned into five distinct ensembles (E1-5: 220, 190, 160, 130, and 100 neurons). The excitatory-to-excitatory connectivity was highly structured to promote balanced, degenerate dynamics. Specifically, these ensembles were embedded via a log-normal within-group excitatory coupling (mean weight ≈ 0.08). To maintain a sparse global average excitatory connectivity of 15%, the intra-cluster connection probability (56.3%) was scaled significantly higher than the baseline inter-cluster probability (4.7%, which was further penalized to an effective 1.4% by a factor of 0.3 for cross-ensemble connections). This established a strict structural clustering ratio (*R_EE_*= 12.0). All other cross-population connections (E→I, I→E, I→I) were assigned using a uniform random connection probability (50%), with fixed synaptic weights scaled by connection type (E→I: 0.10, I→E: −0.20, I→I: −0.15).

To simulate the hypothesized metabolic fatigue driving the observed homeostatic rotations, we introduced an activity-dependent spike rate adaptation (SRA) mechanism. SRA was modeled as a slow inhibitory adaptation current that increments by a fixed Δ after each action potential. Importantly, this load decays over three biological timescales (*τ_k_* 10, 100, 1000 ms), accurately reflecting the complex accumulation of biophysical and metabolic fatigue displayed in cortical neurons.^33^ This mechanism forces heavily active ensembles to progressively suppress their own spiking activity, transferring computational load to rested clusters within the latent library.

Finally, to simulate continuous reaching behavior, a low-dimensional (2D) oscillatory common drive (comprising of offset sin and cos waves) with randomized neuronal gains was continuously injected into the excitatory population. To compensate for the smaller network size and higher relative spiking noise of the simulation, population spike trains were binned at 100 ms and evaluated in sequential 5-second blocks using the same PCA and K-Means analytical pipelines applied to the empirical primate recordings. This computational result proves that single neuron fatigue (SRA) operating within a structurally clustered recurrent network is biophysically sufficient to generate the empirical rank-swapping and geometric transport observed *in vivo*.

Importantly, the model results do not rely on an artificially strong adaptation force. In the normalized leaky integrate-and-fire formulation—where the subthreshold voltage distance from reset (*V*_re_ = 0.0) to spike threshold (*V*_th_ = 1.0) is strictly unity—the total adaptation increment is set to Δ_SRA_ = 0.04. Therefore, each action potential generates an outward adaptation current equivalent to just 4% of the net threshold. Because this increment is distributed across three independent timescales,^33^ the immediate per-spike contribution to any individual decay timescale is only ~ 1.3% of threshold (Δ_SRA_ *K ≈* 0.0133). In physical units, assuming a canonical cortical pyramidal neuron threshold of Δ*V ≈* 20 mV, our parameterization maps to an effective spike-triggered hyperpolarizing deflection of roughly 0.8 mV per spike. This is in agreement with empirical multi-timescale adaptation currents fitted directly to patch-clamp recordings of regular-spiking (RS) excitatory cortical neurons^33,68^ and aligns with typical parameterizations of the adaptive exponential integrate-and-fire (AdEx) model, where spike-triggered adaptation typically contributes 2% to 5% of rheobase per spike.^69^ Furthermore, within the theoretical framework of clustered balanced E-I networks, adaptation strength must remain bounded within a narrow physiological regime (*≈* 2% to 10% of threshold) to support metastable ensemble activation.^31,32,70^ Values below this range fail to overcome recurrent attractor trapping, whereas excessive adaptation (≥15% of threshold) acts as an unphysiological forcing mechanism that degrades E-I balance and induces synchronous population bursting. Thus, our results demonstrate that biologically plausible adaptation is sufficient to gently destabilize local attractor basins and drive fluid, low-dimensional manifold transport without compromising macroscopic network stability.

### Connection between high– and low-dimensional covariance

In conventional population state space analysis, PCA is applied to extract a low-dimensional state 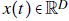 from a high-dimensional population state 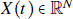, where *D ≪ N*. Although the results we describe in this article operate on these low-dimensional latent dynamics to preserve stable behavior, the underlying physiological constraints (e.g., spike rate adaptation) occur at the level of individual neurons in the high-dimensional ambient space. To mathematically justify our approach of tracking individual neuron contributions to the state eigenvectors, we demonstrate below that the low-dimensional Lyapunov equation is an exact orthogonal projection of the high-dimensional population dynamics. This algebraic connection proves that the eigenvectors serve as the explicit biophysical bridge mapping macroscopic stability to microscopic fluctuations.

Let 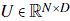 be the matrix containing the top *D* orthogonal eigenvectors (*U^T^U I_D_*) of the high-dimensional data covariance. The projection and reconstruction mappings are thus 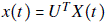 and 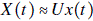. Assuming the high-dimensional state evolves according to Equation 1 (Box 1), the low-dimensional dynamics estimated in the PCA subspace are:

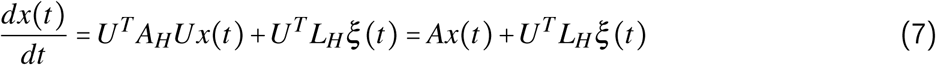

Thus, the estimated low-dimensional dynamics matrix is defined as 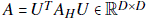. The high-dimensional covariance 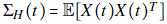 is mapped to the low-dimensional covariance, 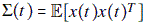, as 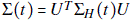. Substituting the projected matrices into the reduced steady-state Lyapunov equation yields:

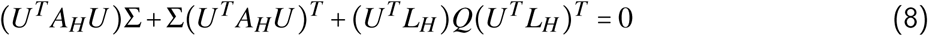

Assuming the behavioral signal is predominantly captured within the active subspace (such that *UU^T^* acts as the identity for the signal covariance), we can factor out *U^T^* on the left and *U* on the right:

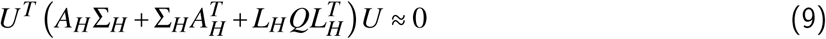

The low-dimensional Lyapunov equation is the exact orthogonal projection of the original high-dimensional Lyapunov equation onto the PCA subspace, forming a direct algebraic connection between the high– and low-dimensional interpretations of the data.

## Author Contributions

Conceptualization – SEC, PN Data collection, SEC Data curation, SEC Formal analysis, SEC Funding acquisition PN, SEC Investigation SEC, PN Methodology SEC, PN Project administration, SEC Resources SEC, PN Software, SEC Supervision, PN Validation, SEC, PN Visualization SEC, PN Writing, original draft, SEC Writing, review and editing, SEC, PN

## Funding

SEC was supported by a Stanford School of Medicine Dean’s Postdoctoral Fellowship, and Stanford’s Human-Centered Artificial Intelligence Seed Grant. Funding for this project was provided to PN by the Wu Tsai Neuroscience Institute and supplemented by NIH grants R01NS130789, R01NS123517, and U19NS118284 awarded to PN.

## Competing Interests

The authors declare that they have no competing interests.

## Data and Code Availability

Data and code used in this study are available from the corresponding author upon reasonable request.

## Inclusion and Ethics

The authors declare that they have no inclusion or ethics concerns for this study. The experiments were carried out under the supervision of Dr. Paul Nuyujukian and in accordance with the three Rs of ethical animal research. Independent analysis of the data was performed entirely using the resources and physical space allocated to the Brain Interfacing Lab, Stanford University.

## Acknowledgements

We thank Michelle S. Wechsler, Alex Paraskevopoulou, Mackenzie J. Risch, Sam W. Baker and Kristina Lebedev for animal care, and Kimberly Chin and Margaret Truong for administrative support. In addition to the authors, the members of the Brain Interfacing Lab who contributed to animal care during experimental data collection include Michelle S. Wechsler, Iliana E. Bray, Michael P. Silvernagel, Alissa S. Ling, and Elizabeth Jun.

## Supplementary Figures

**Figure S1:**
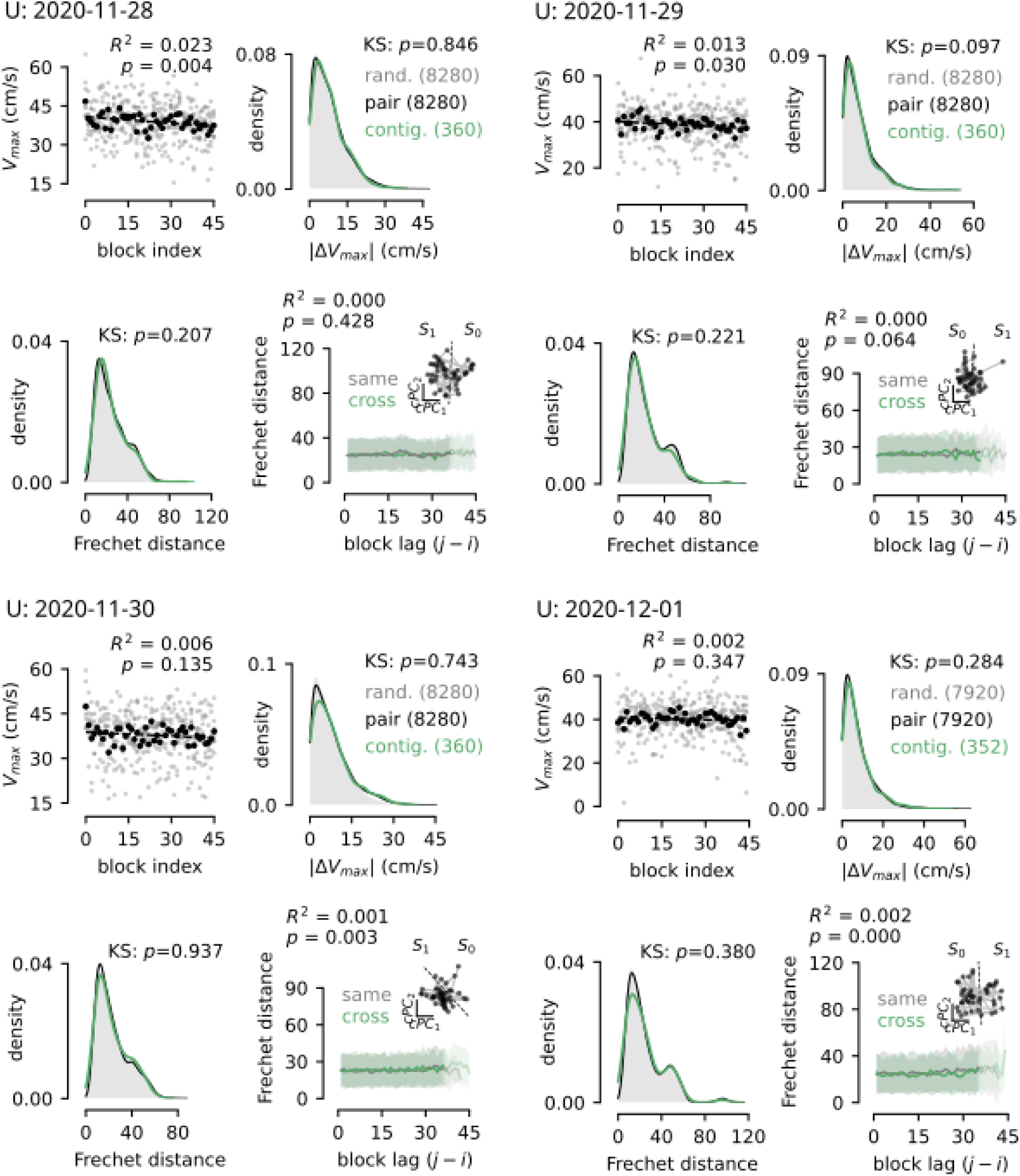
Behavioral consistency plots for Monkey U, sessions 1 through 4.

**Figure S2:**
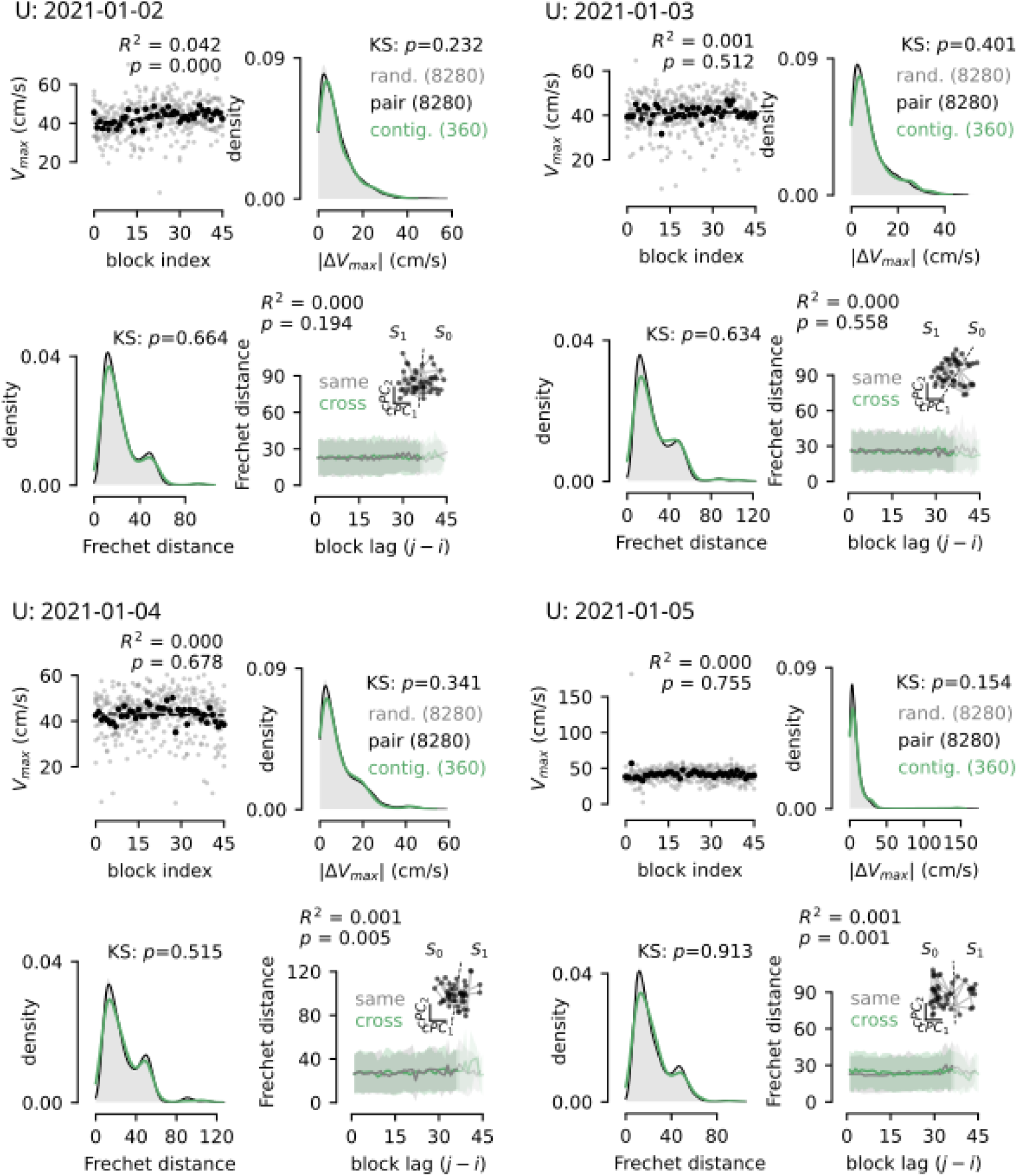
Behavioral consistency plots for Monkey U, sessions 5 through 8.

**Figure S3:**
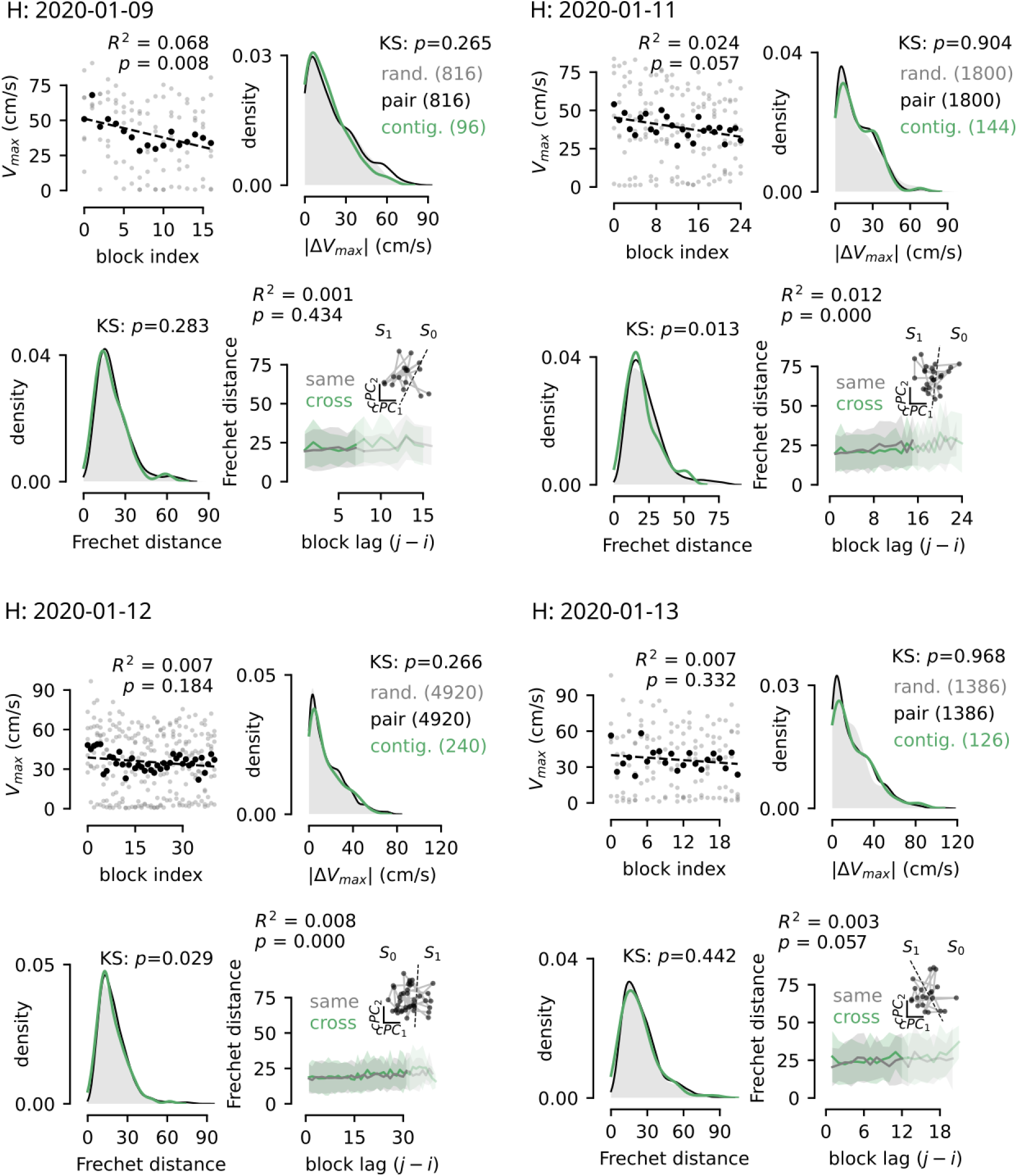
Behavioral consistency plots for Monkey H, sessions 1 through 4.

**Figure S4:**
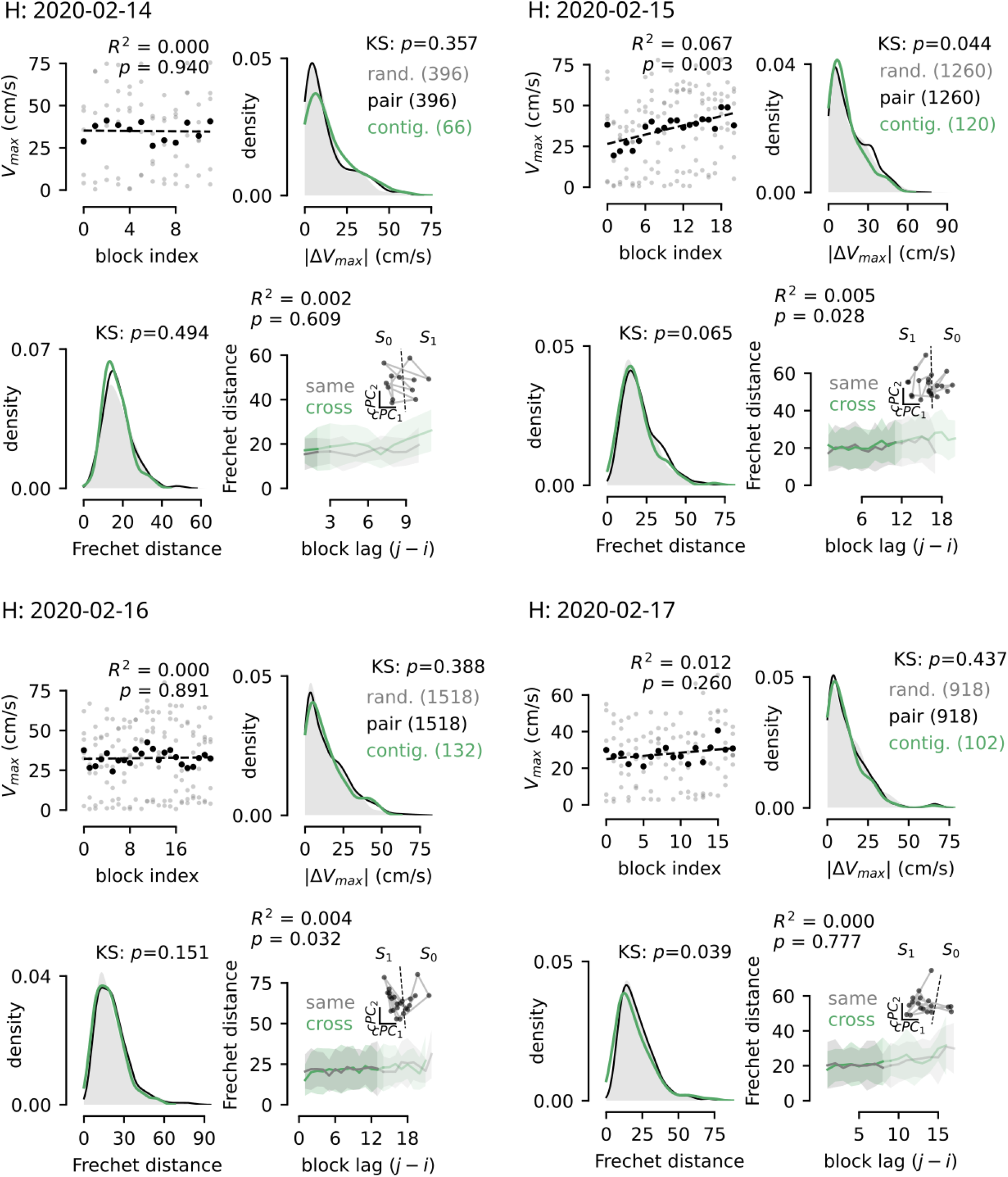
Behavioral consistency plots for Monkey H, sessions 5 through 8.

**Figure S5:**
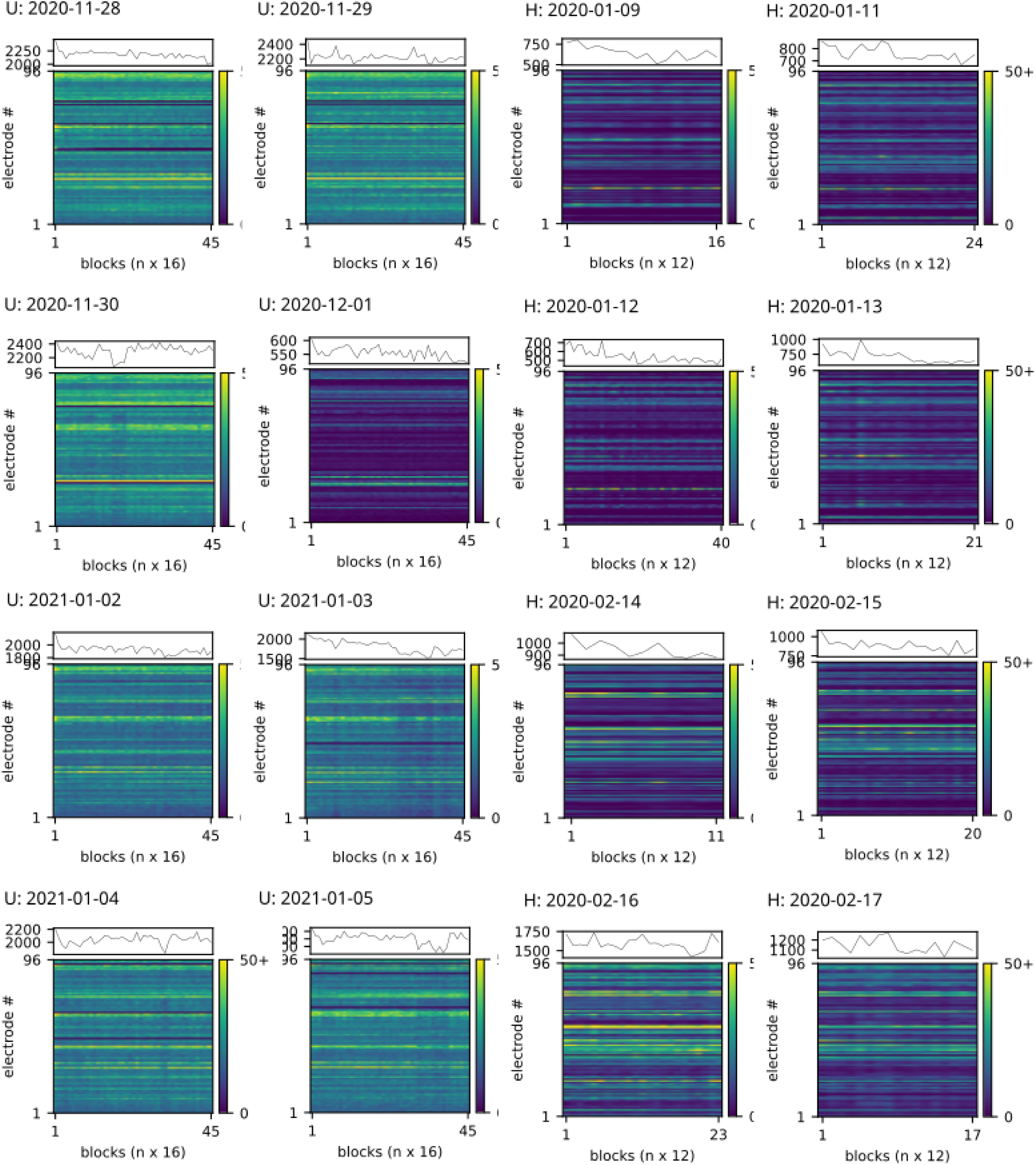
Averaged trial spike counts of individual electrodes and the total population for successive trial blocks, sessions 1 through 8 for both Monkey U and H.

**Figure S6:**
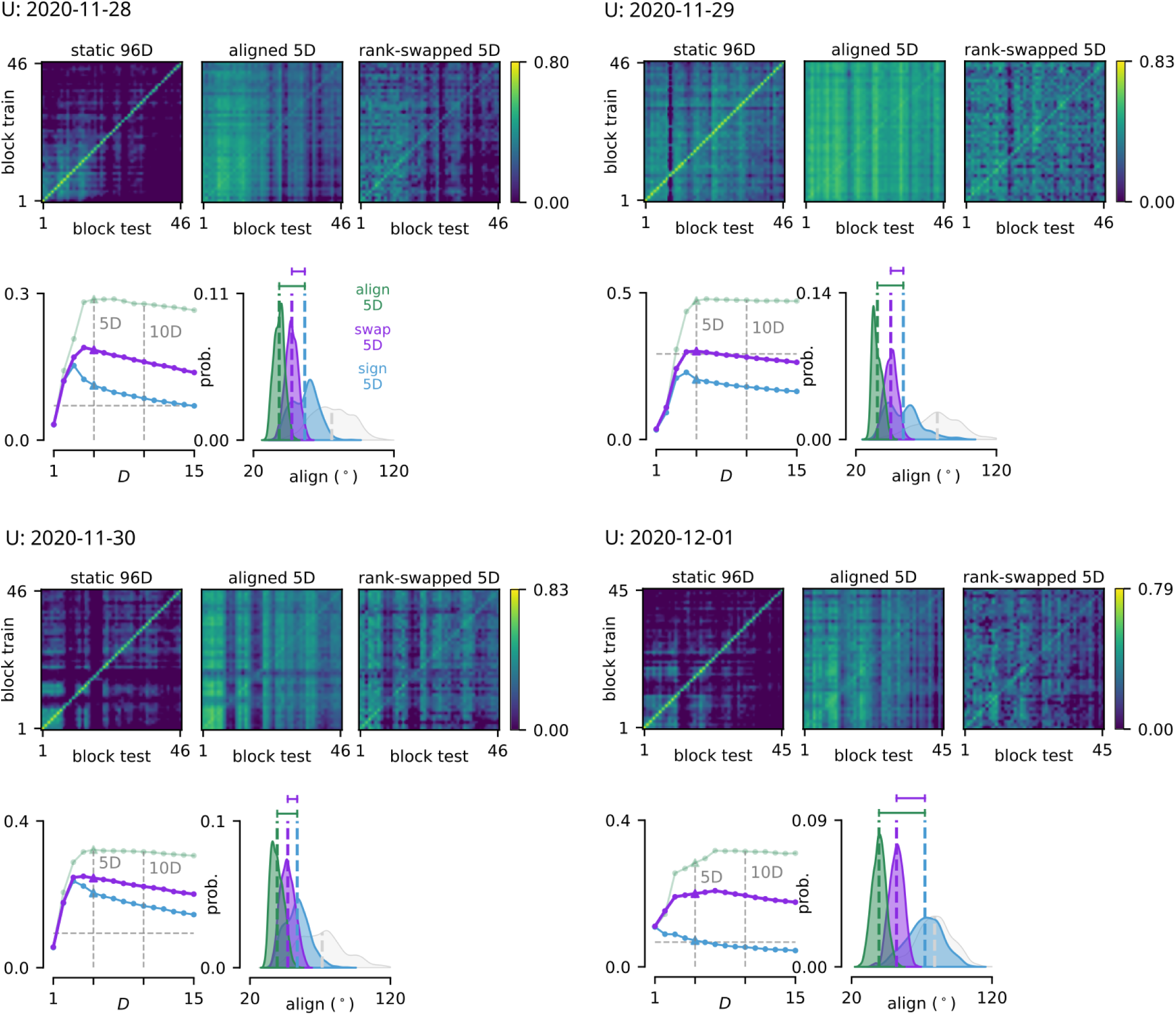
Cross-block kinematic decoding results for Monkey U, sessions 1 through 4.

**Figure S7:**
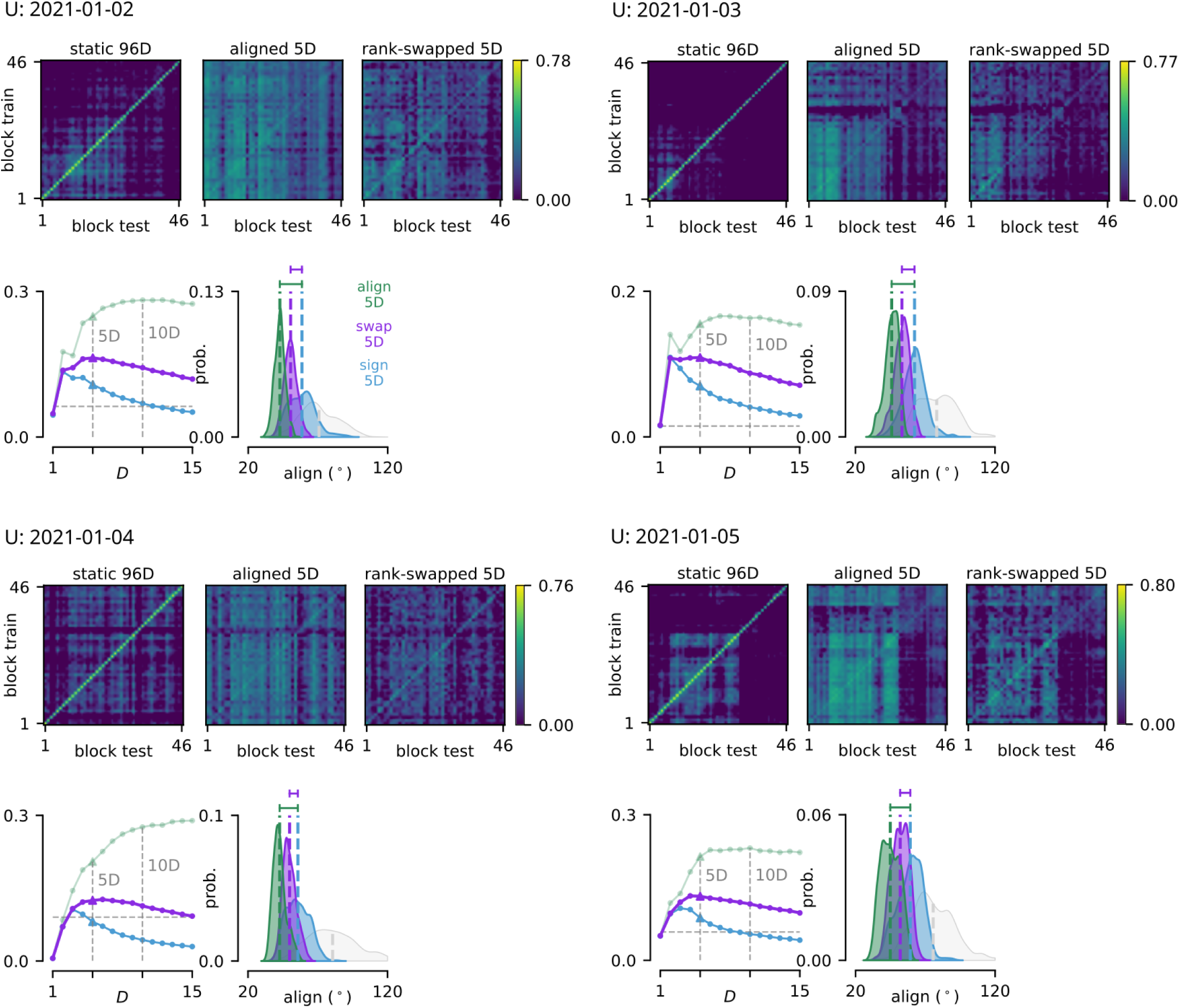
Cross-block kinematic decoding results for Monkey U, sessions 5 through 8.

**Figure S8:**
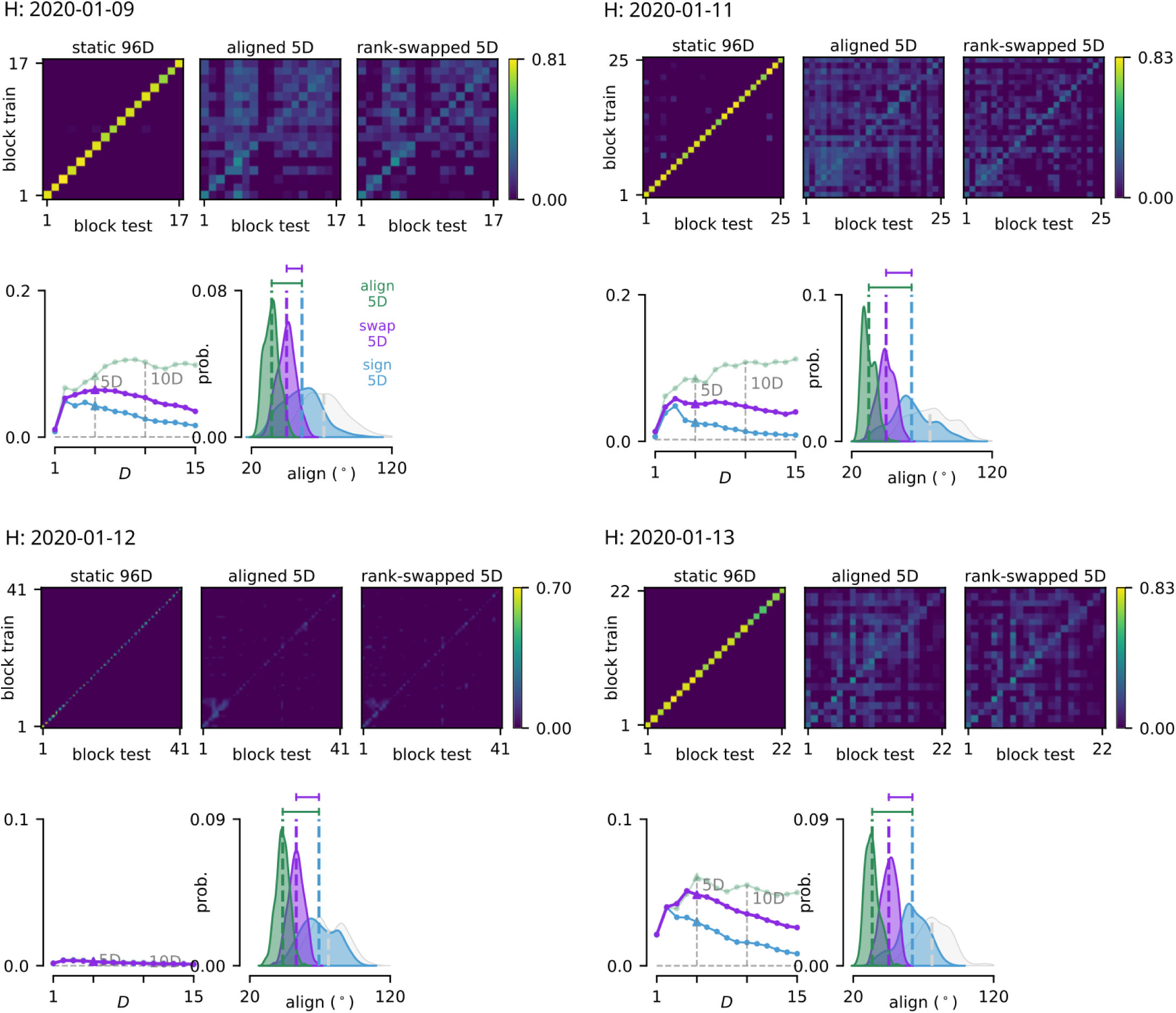
Cross-block kinematic decoding results for Monkey H, sessions 1 through 4. Decoder performance was inexplicably poor on 2020-01-12, however, everything else appeared normal that day so it was not excluded from the dataset.

**Figure S9:**
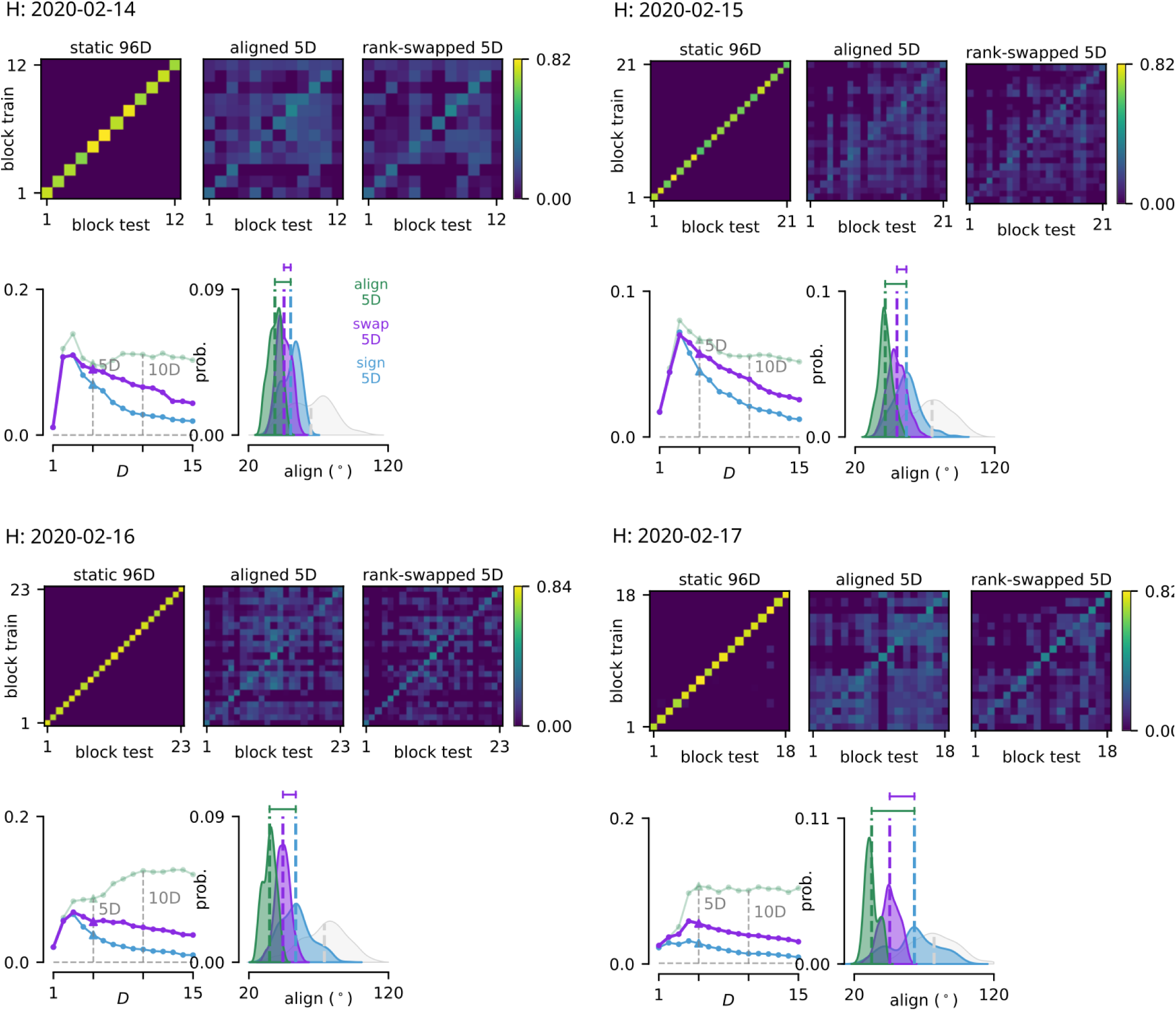
Cross-block kinematic decoding results for Monkey H, sessions 5 through 8.

**Figure S10:**
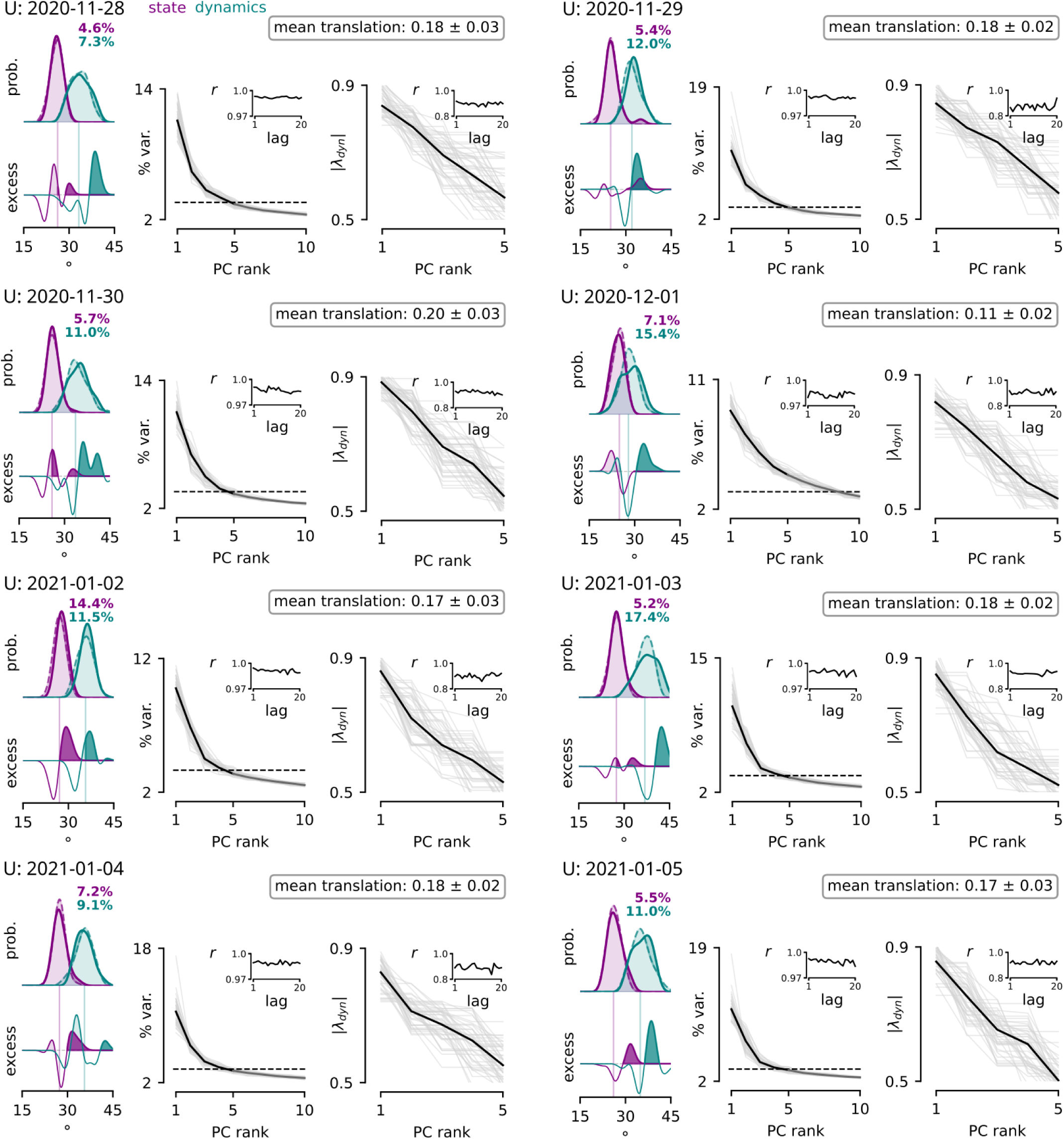
State and dynamics subspace transport for Monkey U, sessions 1 through 8.

**Figure S11:**
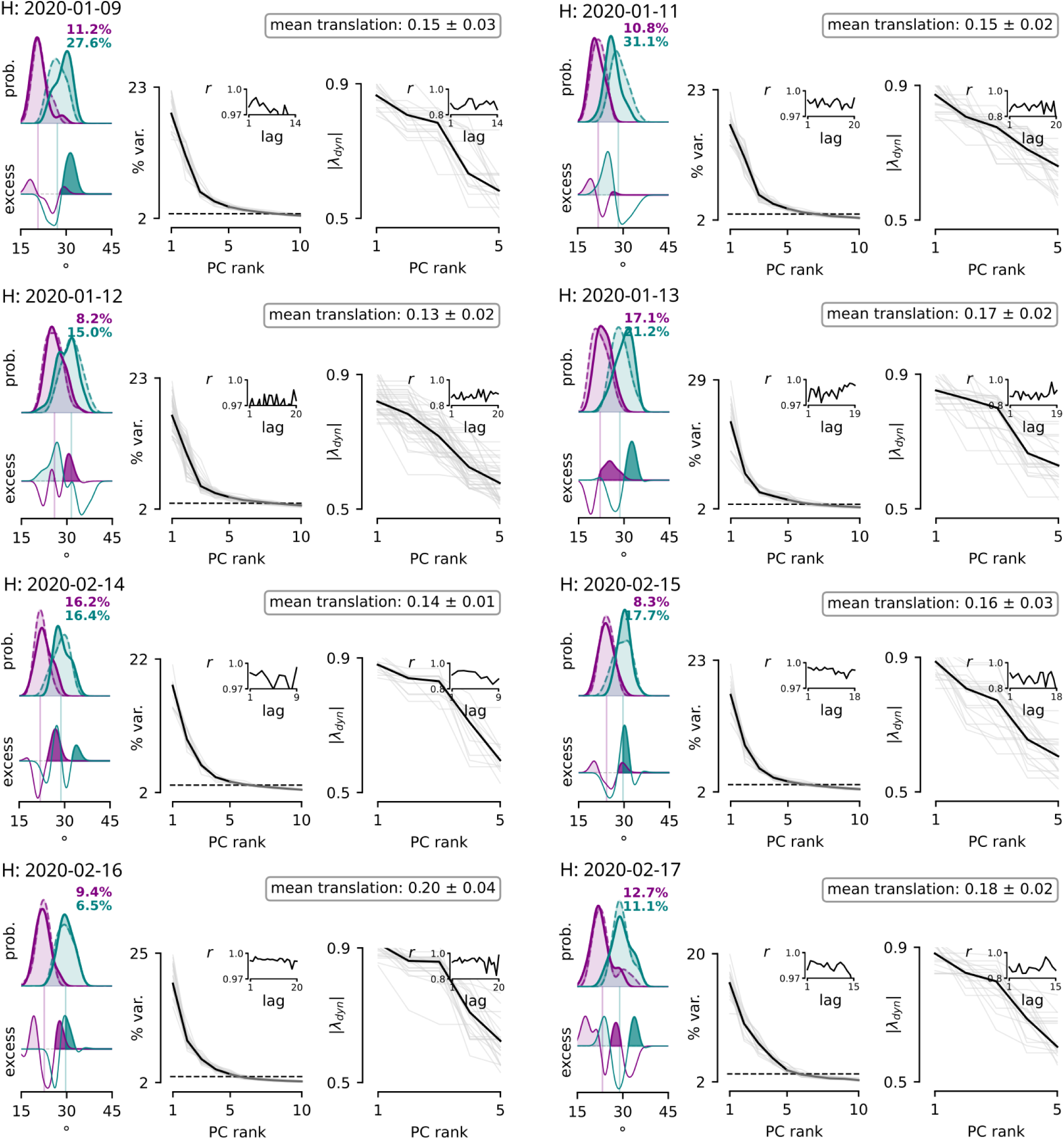
State and dynamics subspace transport for Monkey H, sessions 1 through 8.

**Figure S12:**
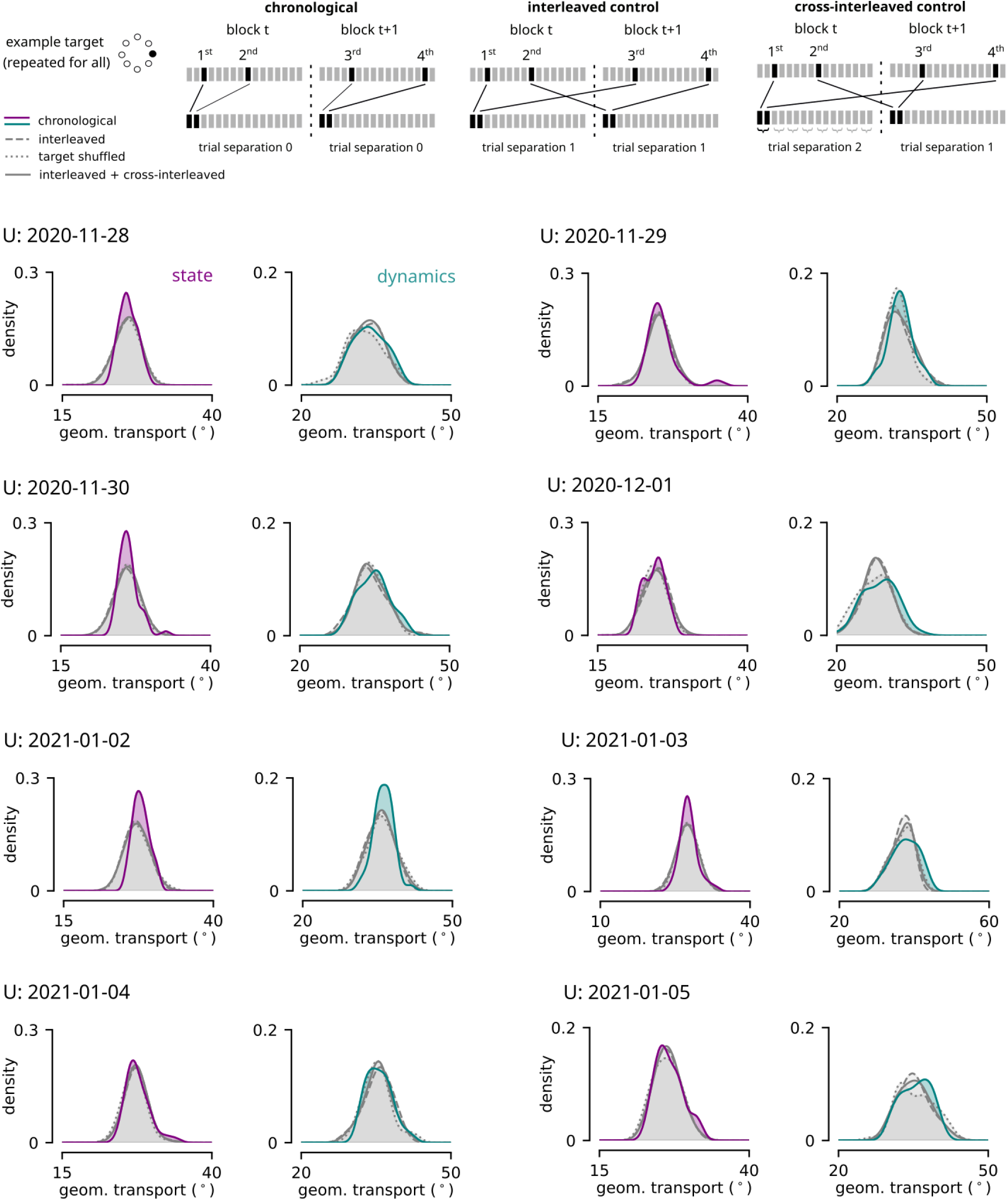
Three shuffle controls for state and dynamics subspace transport, Monkey U sessions 1 through 8. To distinguish true drift from estimation noise, block-to-block transport was compared to three non-chronological controls. If block A and block B were different states, then re-ordering their trials should create a more blended average state (like pooling trials across an entire session). The first control interleaved target-specific trials and was exactly sample-size matched (main text). The second control, which pooled both interleaved and cross-interleaved trial orderings, used the exact same data but boosted the sample size two-fold for better estimation of the blended state. The third asynchronous control distribution was sample-size matched, but shuffled trial sequence randomly across blocks.

**Figure S13:**
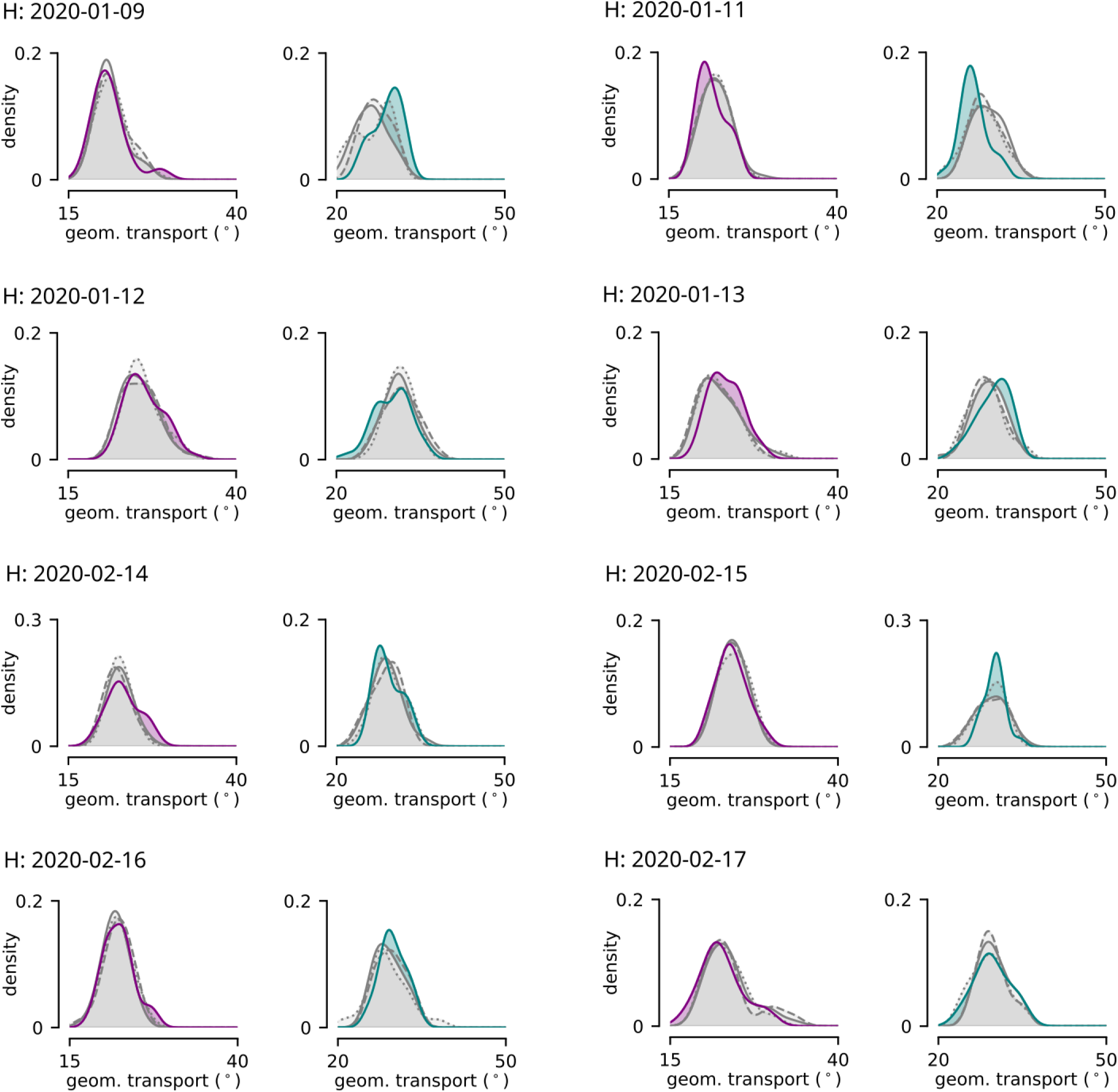
Three shuffle controls for state and dynamics subspace transport, Monkey H sessions 1 through 8.

**Figure S14:**
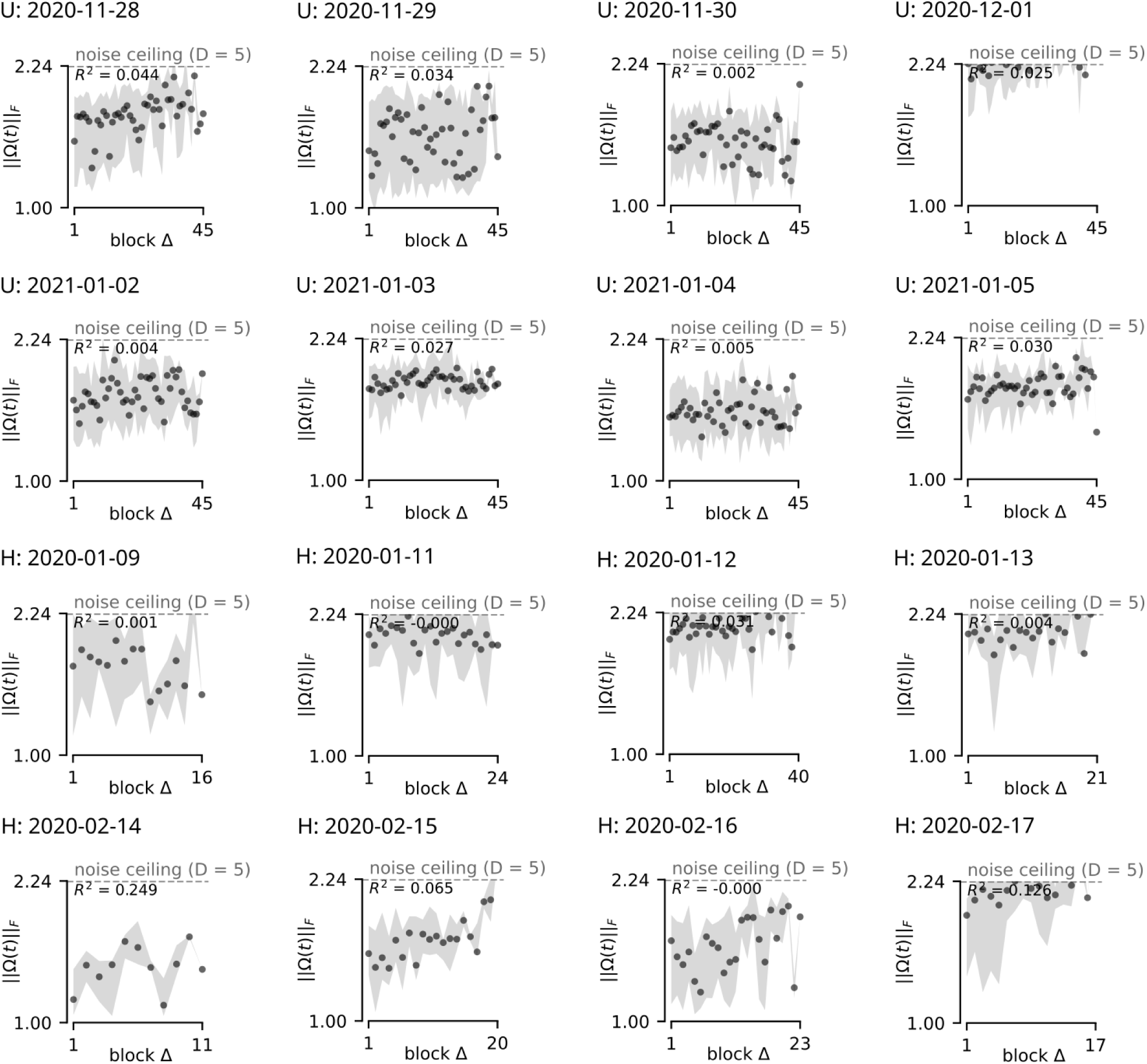
The transport for sessions 1-8, Monkeys U and H.

**Figure S15:**
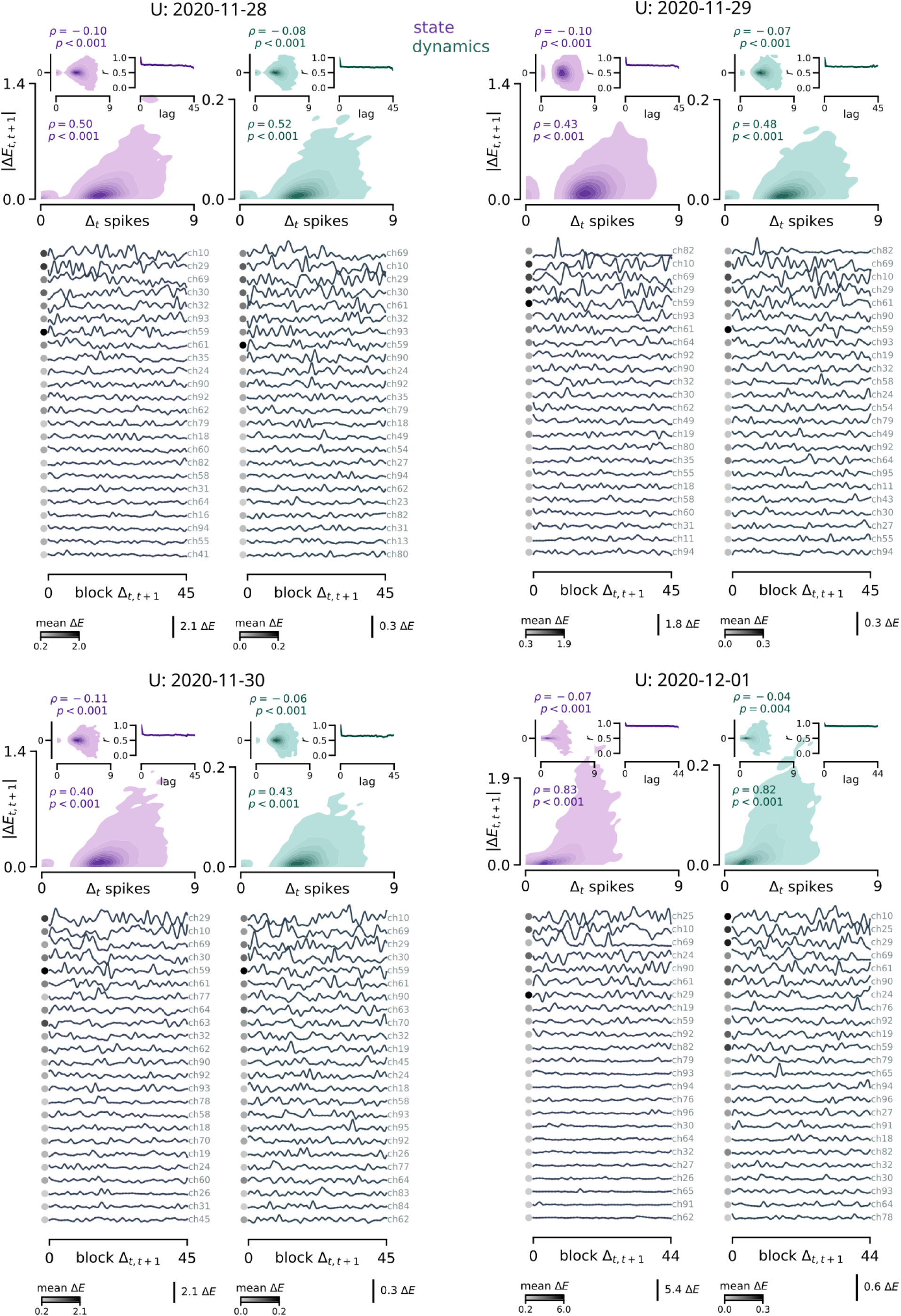
Subspace energy plots for Monkey U, sessions 1 through 4.

**Figure S16:**
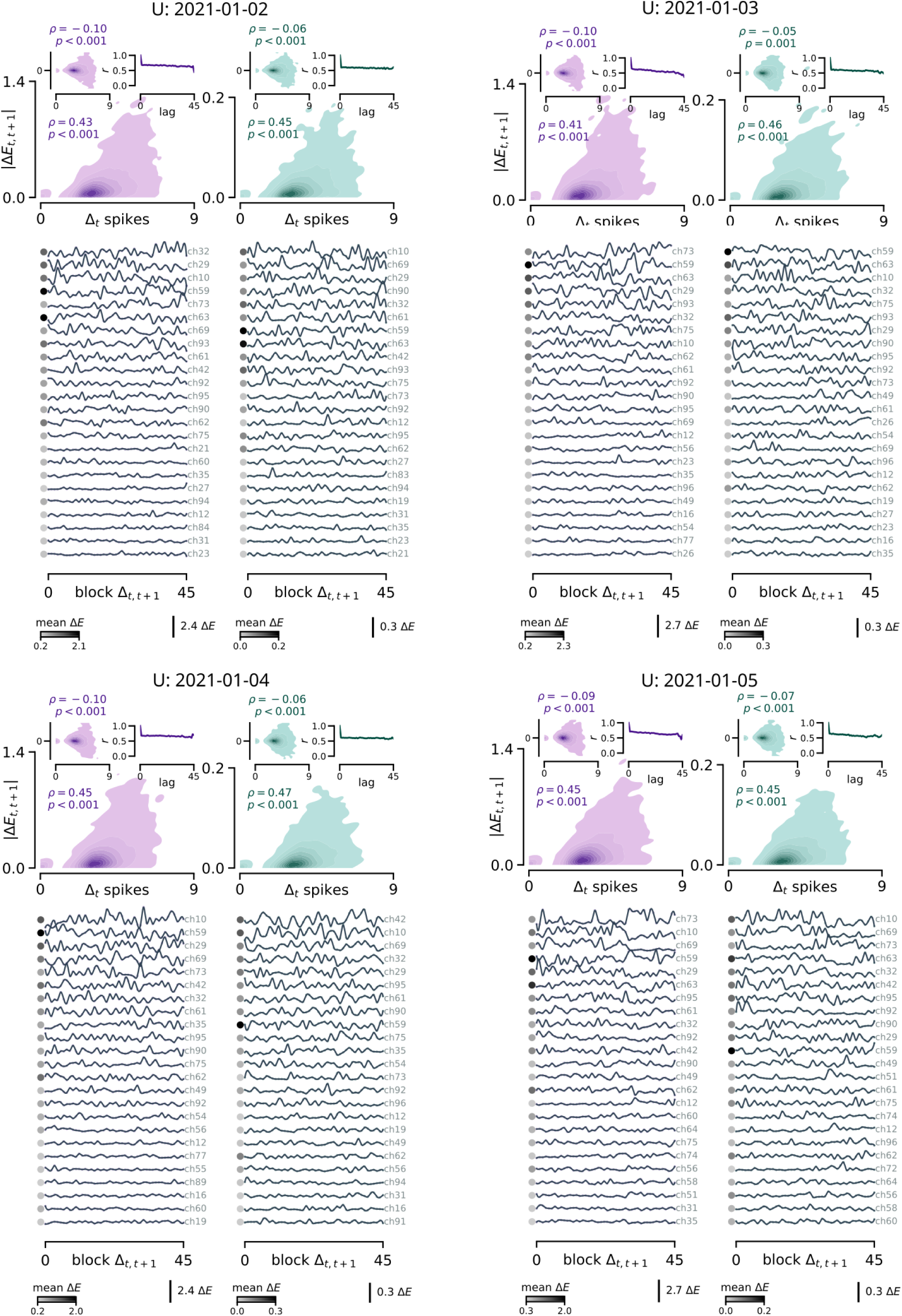
Subspace energy plots for Monkey U, sessions 5 through 8.

**Figure S17:**
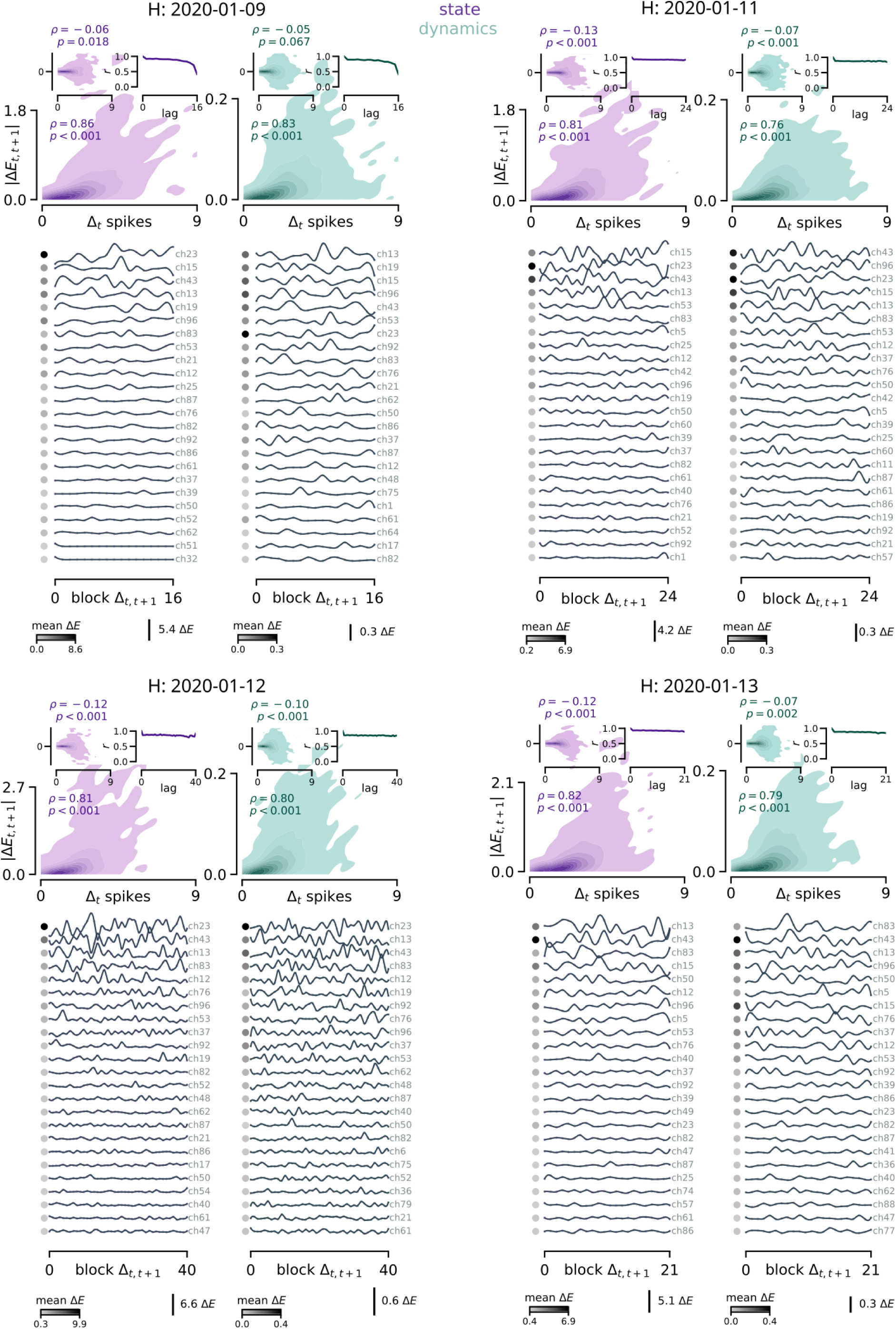
Subspace energy plots for Monkey H, sessions 1 through 4.

**Figure S18:**
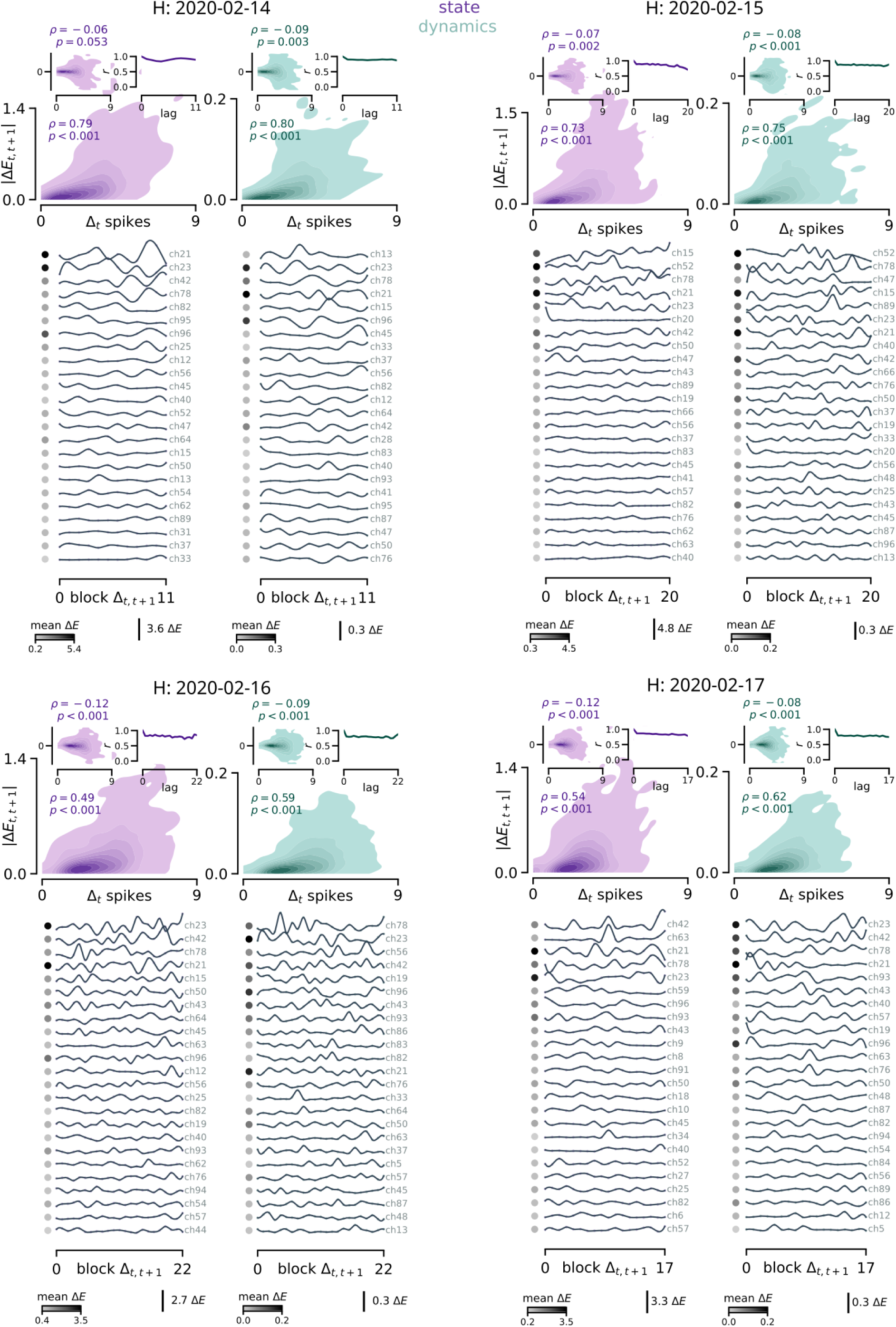
Subspace energy plots for Monkey H, sessions 5 through 8.

**Figure S19:**
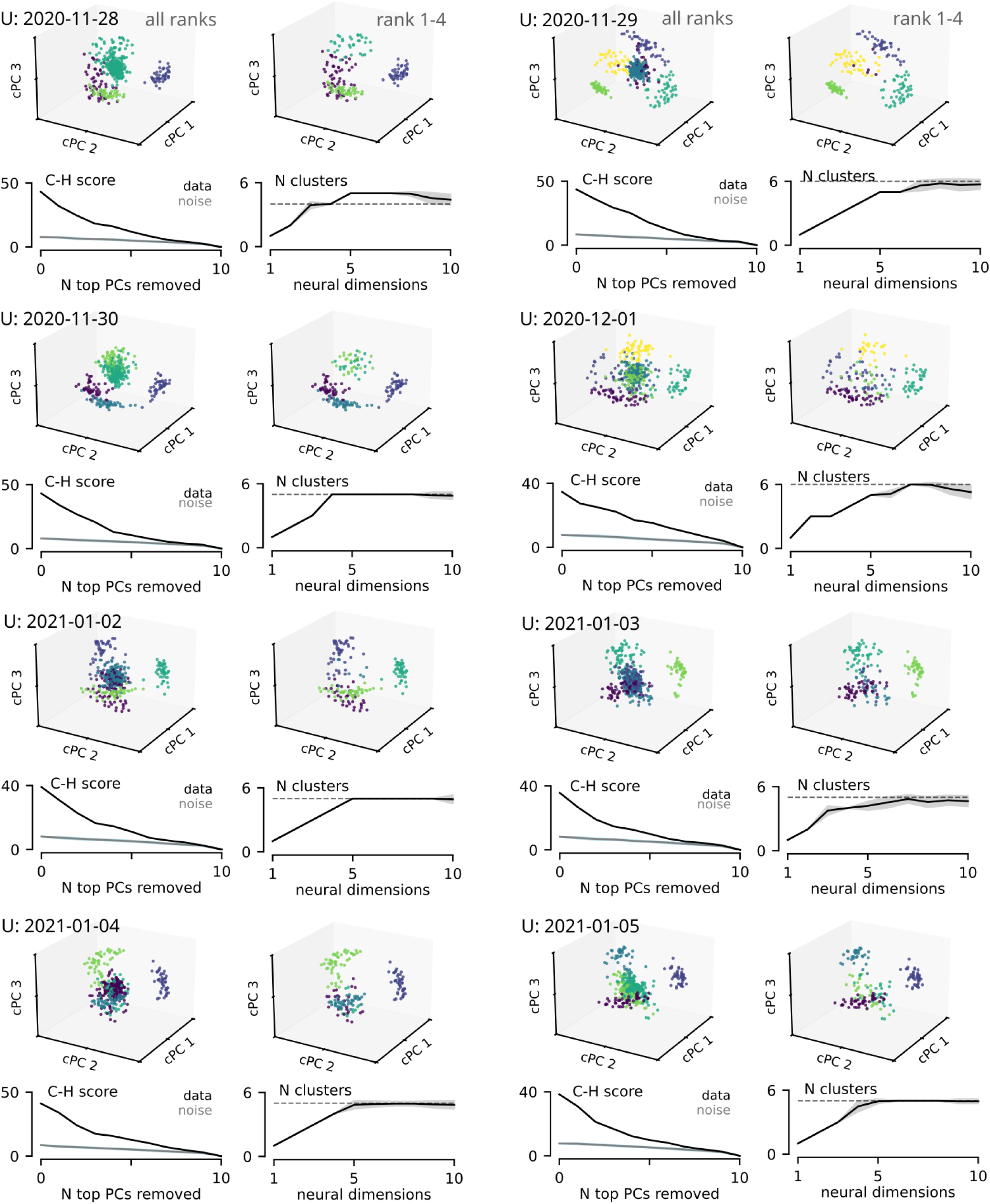
Clustering of population state eigenvectors for Monkey U, sessions 1 through 8.

**Figure S20:**
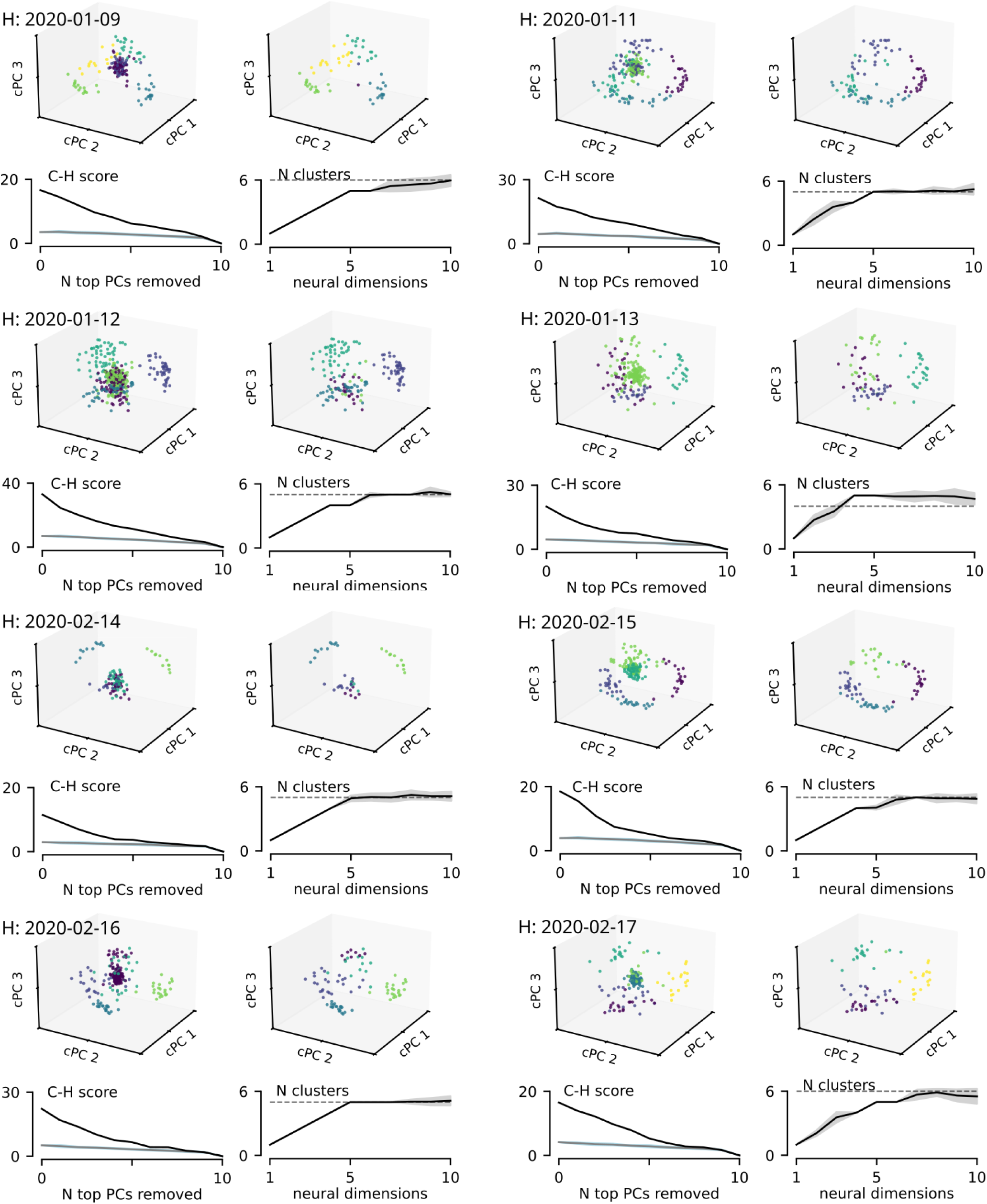
Clustering of population state eigenvectors for Monkey H, sessions 1 through 8.

**Figure S21:**
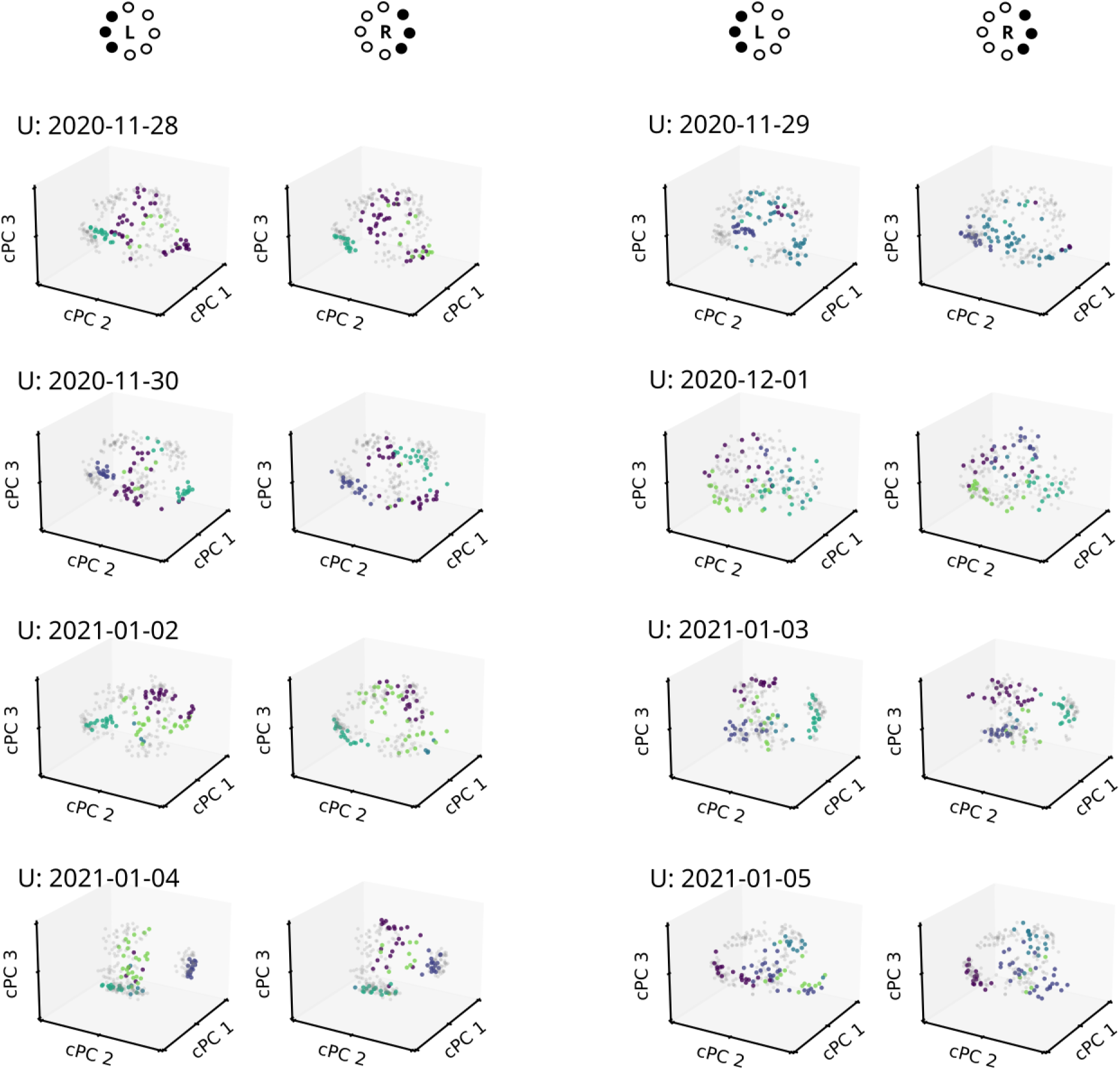
Overlap of state eigenvectors computed on leftward versus rightward sets of targets, Monkey U sessions 1 through 8.

**Figure S22:**
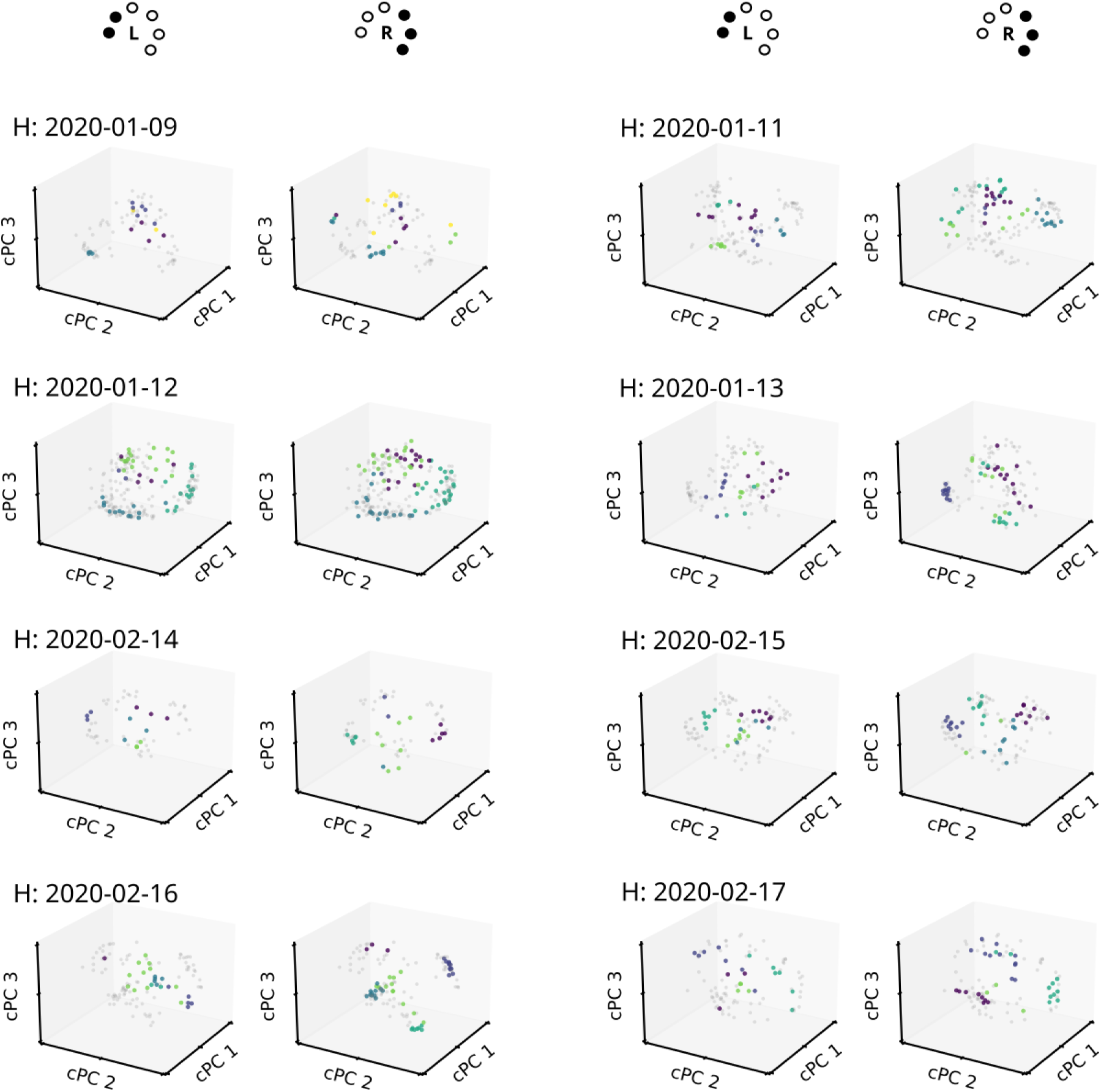
Overlap of state eigenvectors computed on leftward versus rightward sets of targets, Monkey H sessions 1 through 8. Note, the bottom-left target was not presented for Monkey H due to a visual deficit. In this case leftward only contained left and top-left target locations.

**Figure S23:**
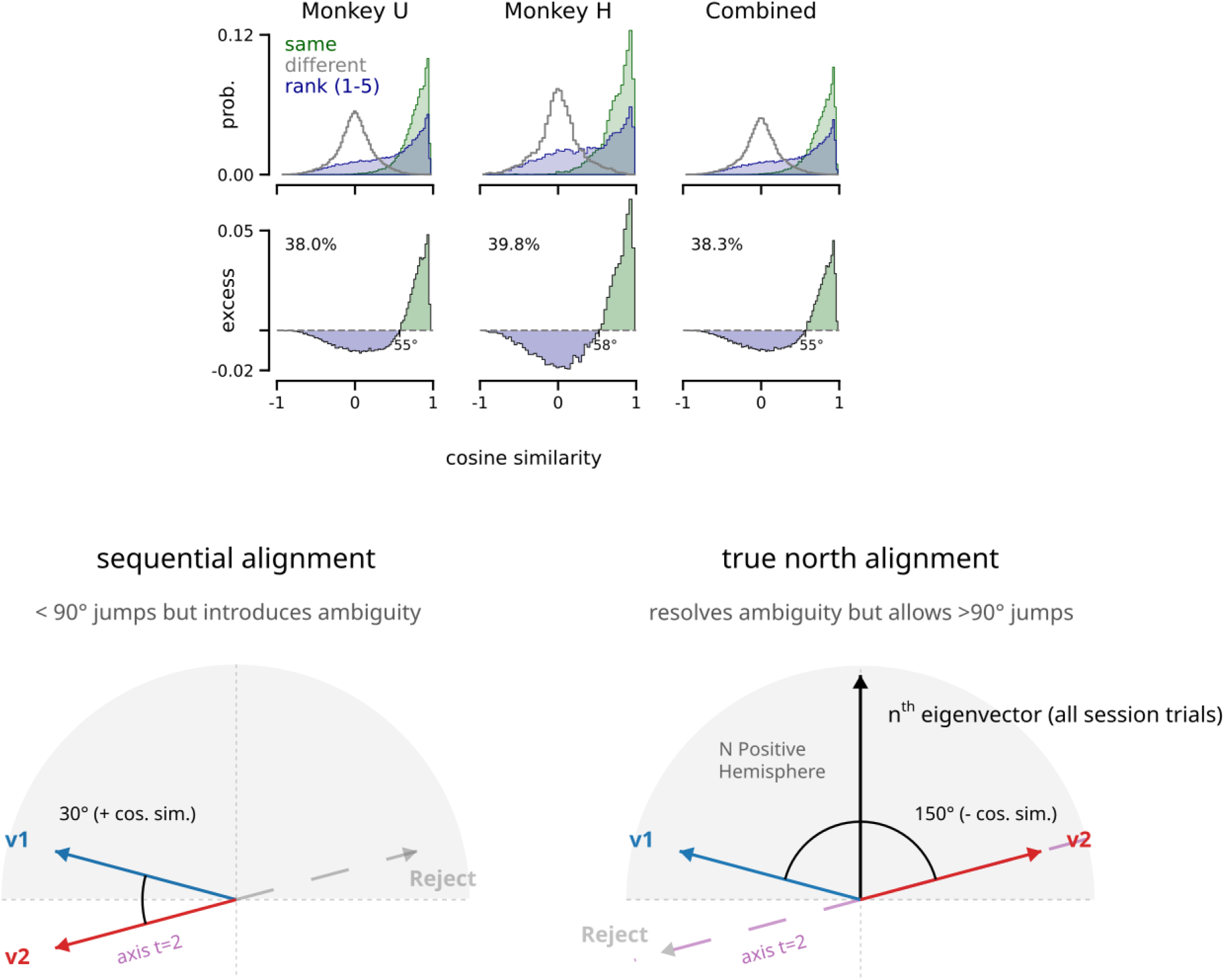
(*Top*) Extension of the distributions shown in Figure 3i of the main text to include ranks 1-5. The inclusion of ranks 4 and 5 led to larger leftward shifts in the rank comparison distribution relative to the within-cluster (same) comparisons. (*Bottom*) Sequential versus global eigenvector alignment. Sign correction of eigenvectors was essential for this analysis since PCA can arbitrarily return sign-flipped axes. (*Left*) Sequential alignment minimizes the angular distance between consecutive time steps by greedily selecting the eigenvector orientation (*v*_2_) closest to the previous block (*v*_1_). While this ensures local smoothness, slow continuous rotations can accumulate over an experimental session, allowing the subspace to fundamentally invert relative to its starting orientation. (*Right*) True north alignment enforces global consistency by requiring all eigenvectors to maintain a positive projection onto fixed, global anchor vectors determined by the unique session-wide eigenvectors associated with the top 10 ranks. This prevents artifactual sign-flipping but exposes the underlying rotation as an abrupt 90^○^ angular jump (cosine similarity < 0) when the continuous rotation crosses the equator of the true north hemisphere.

**Figure S24:**
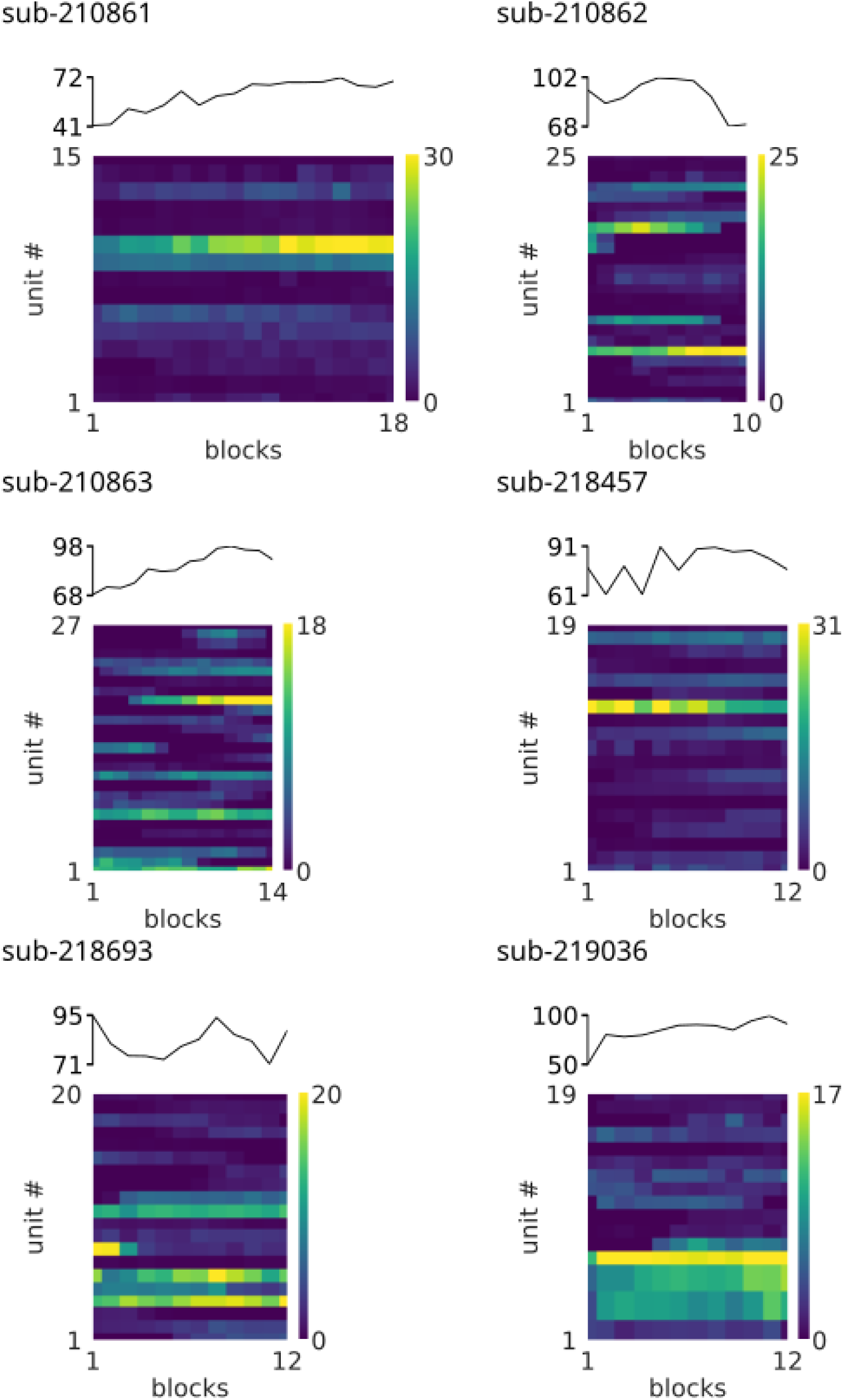
Averaged trial spike counts of individual electrodes and the total population, measured over session trial blocks for 6 mice.

**Figure S25:**
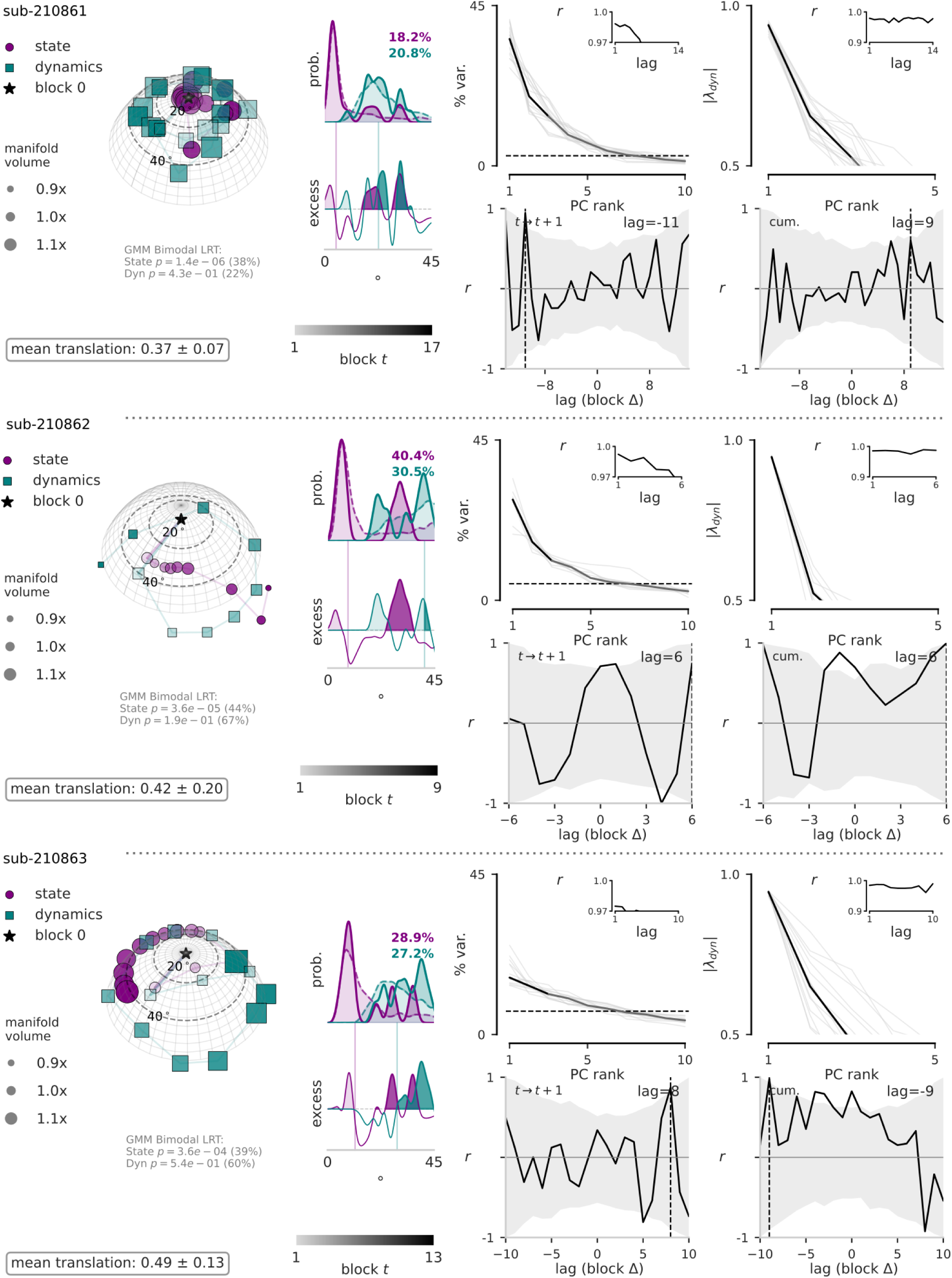
State and dynamics subspace transport for three of six mice.

**Figure S26:**
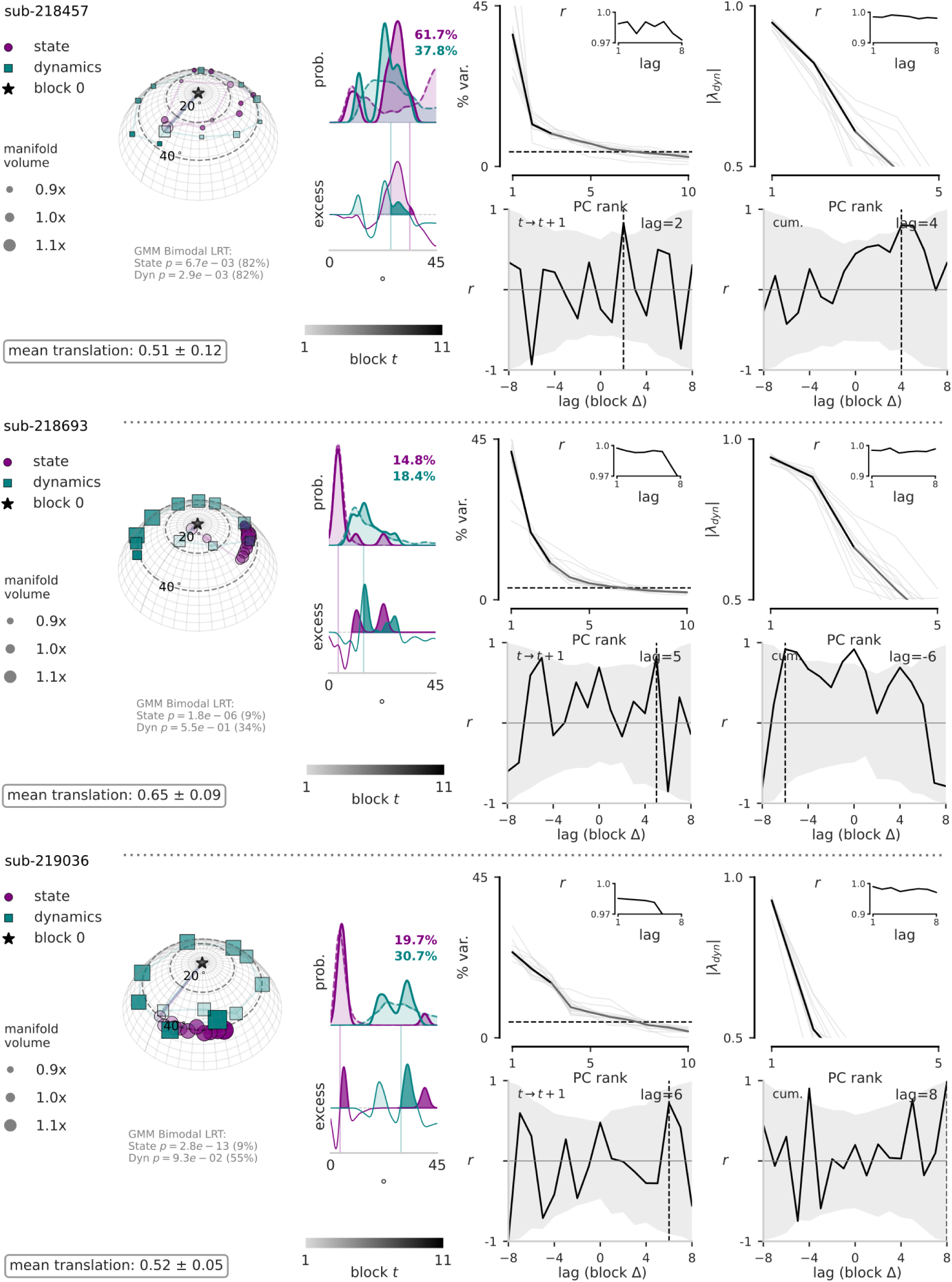
State and dynamics subspace transport for the other three mice.

**Figure S27:**
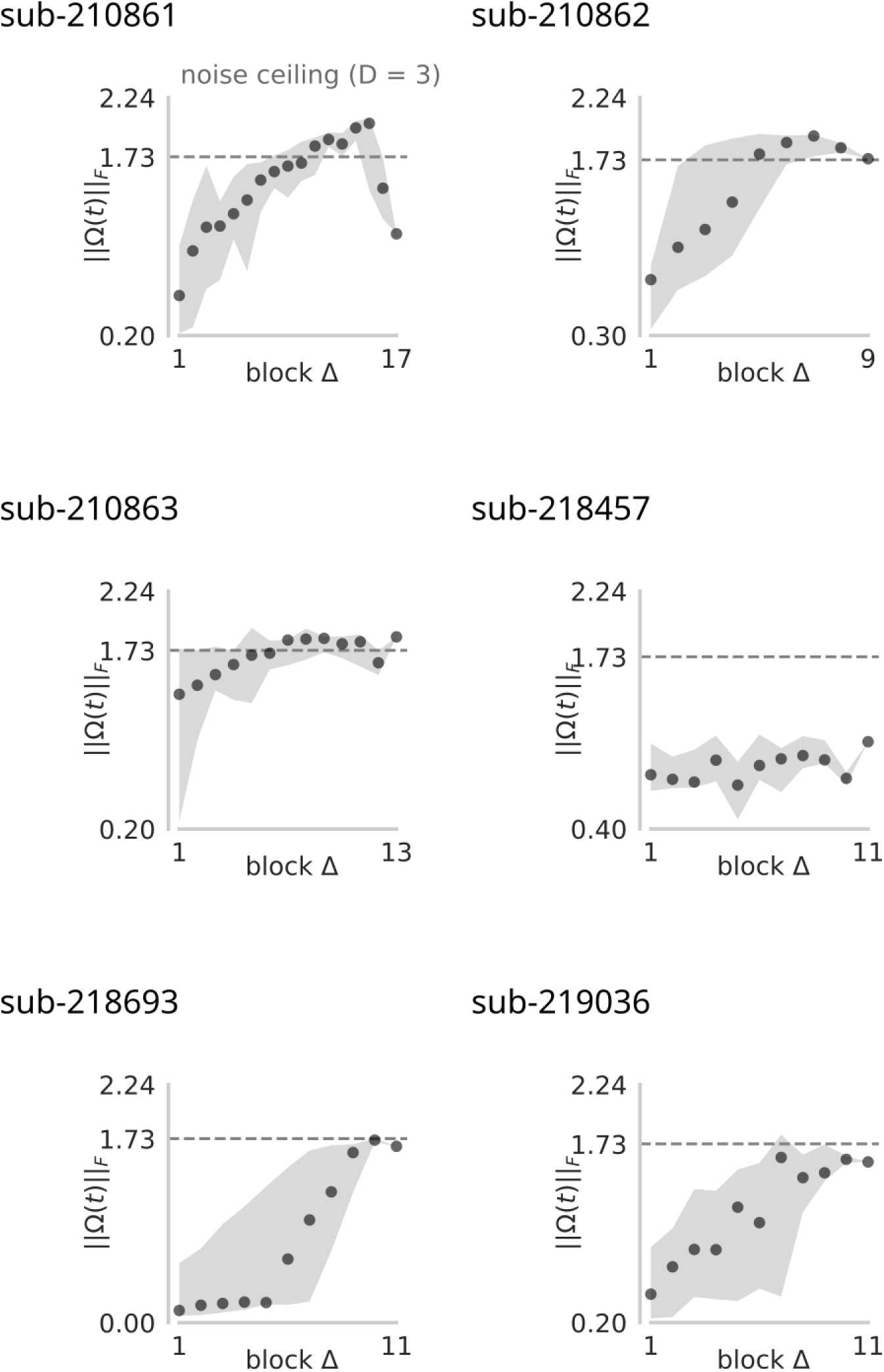
The session transport for each of the six mice.

**Figure S28:**
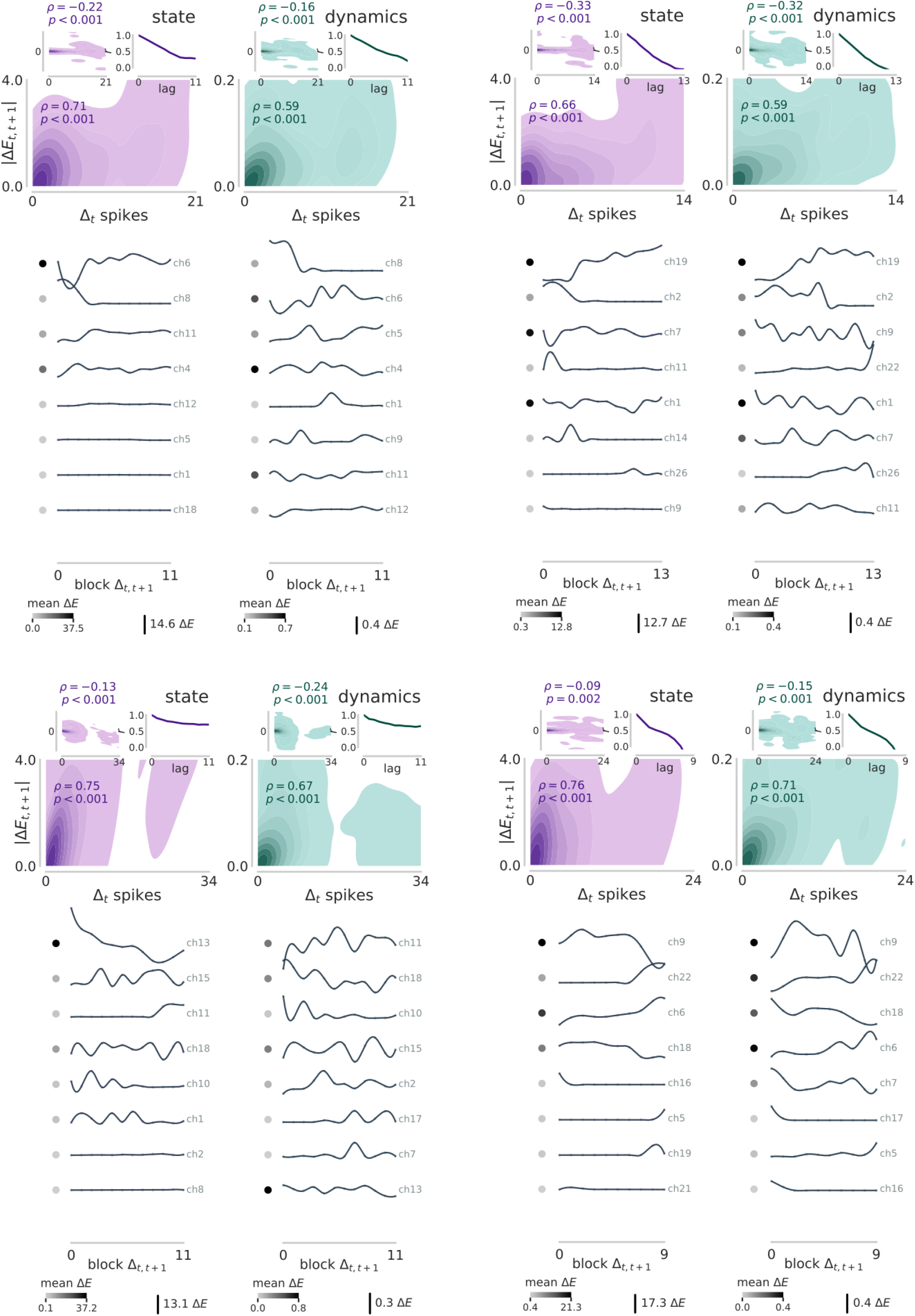
Subspace energy plots for four of the six mice.

**Figure S29:**
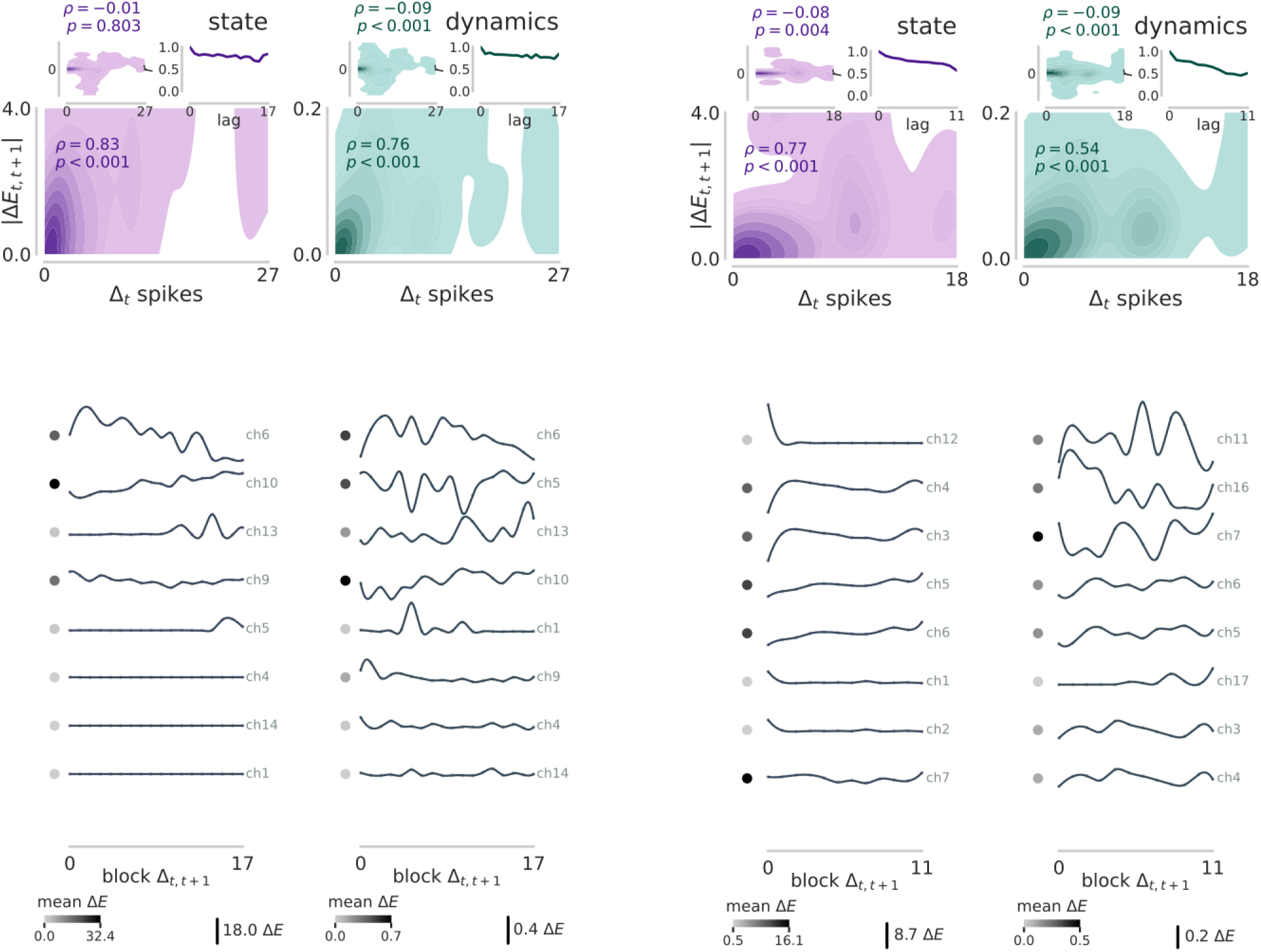
Subspace energy plots for the remaining two mice.

**Figure S30:**
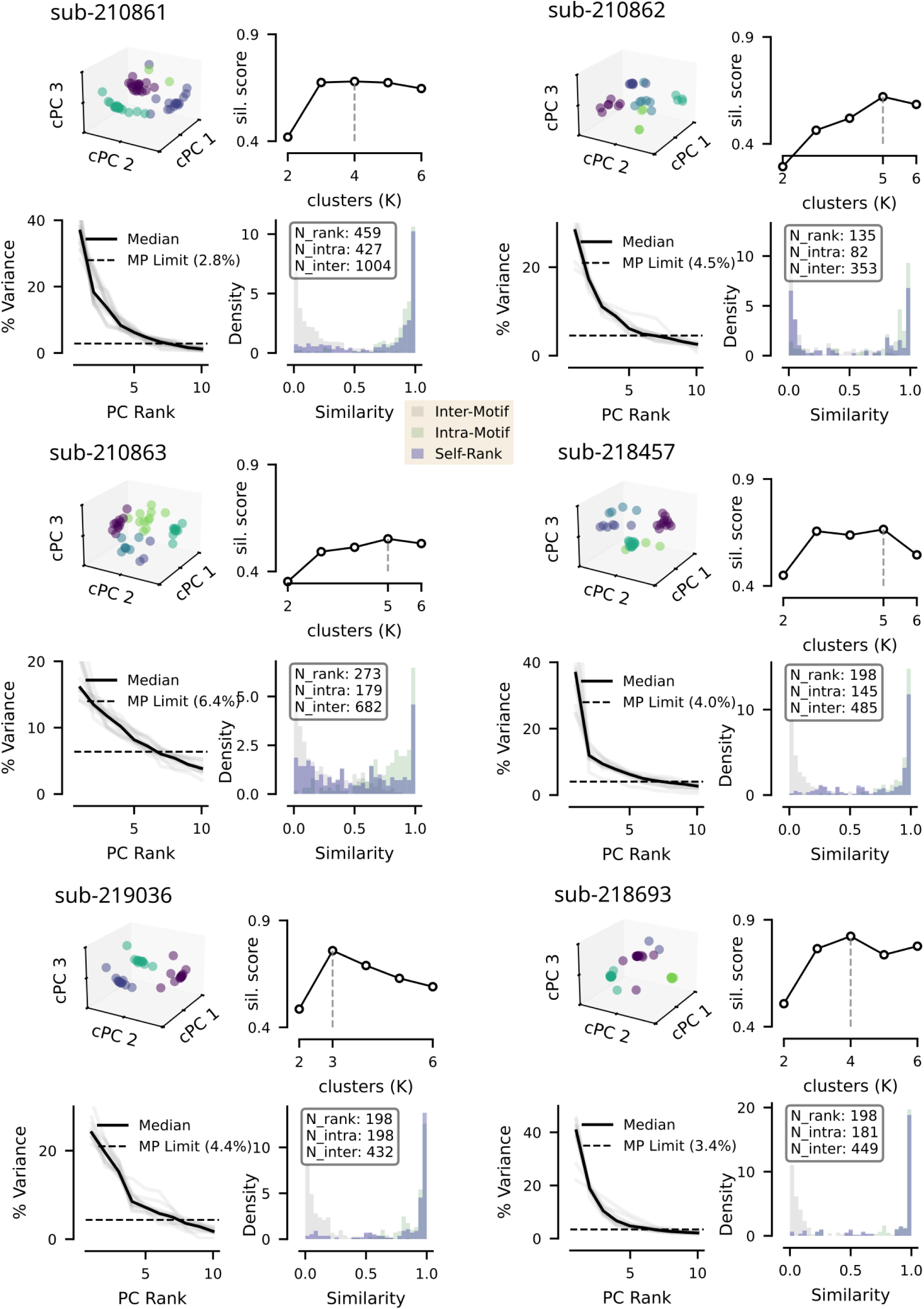
Clustering of population state eigenvectors for all six mice.

**Figure S31:**
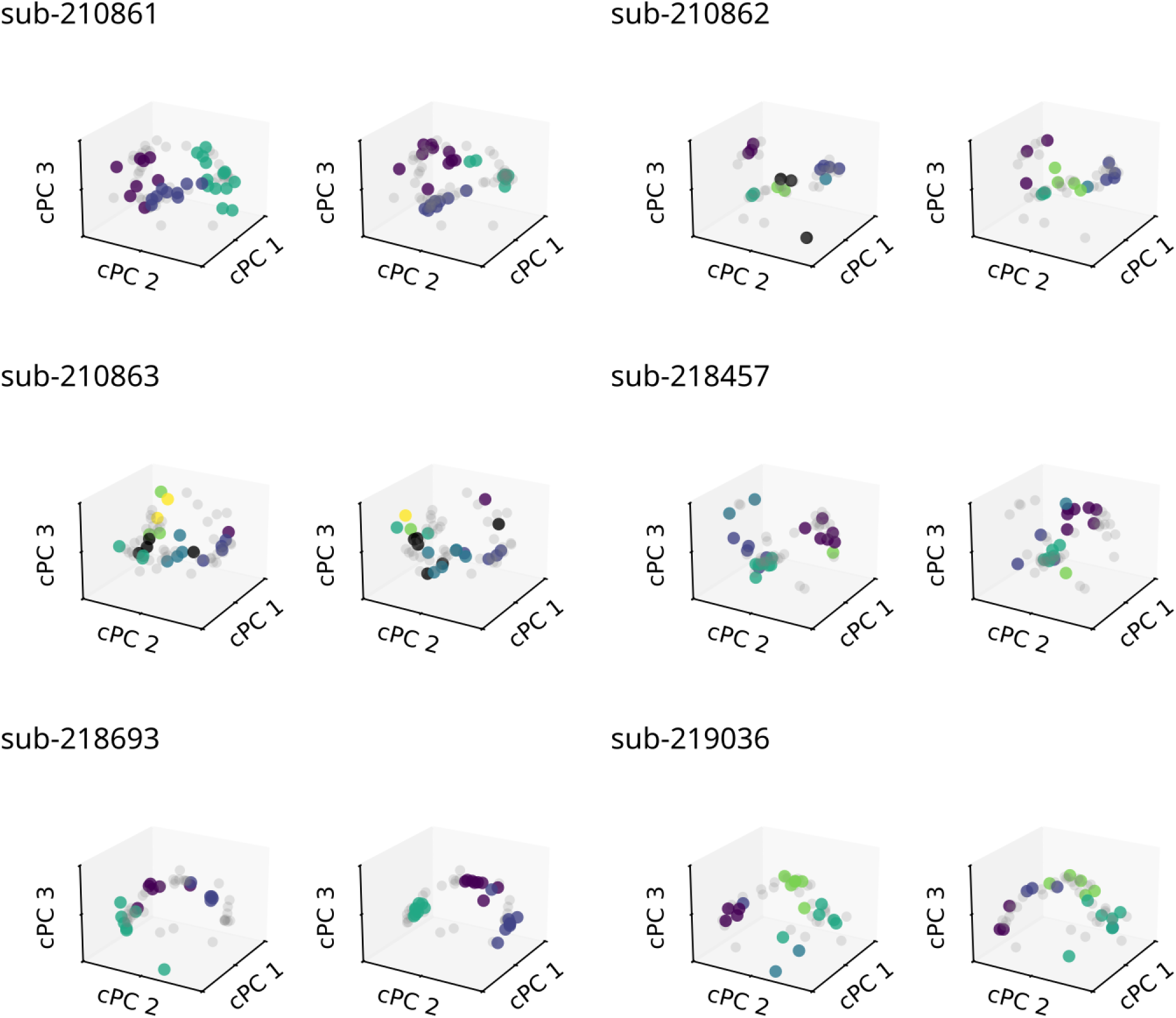
Overlap of state eigenvectors computed on left versus right lick trials for all six mice.

**Figure S32:**
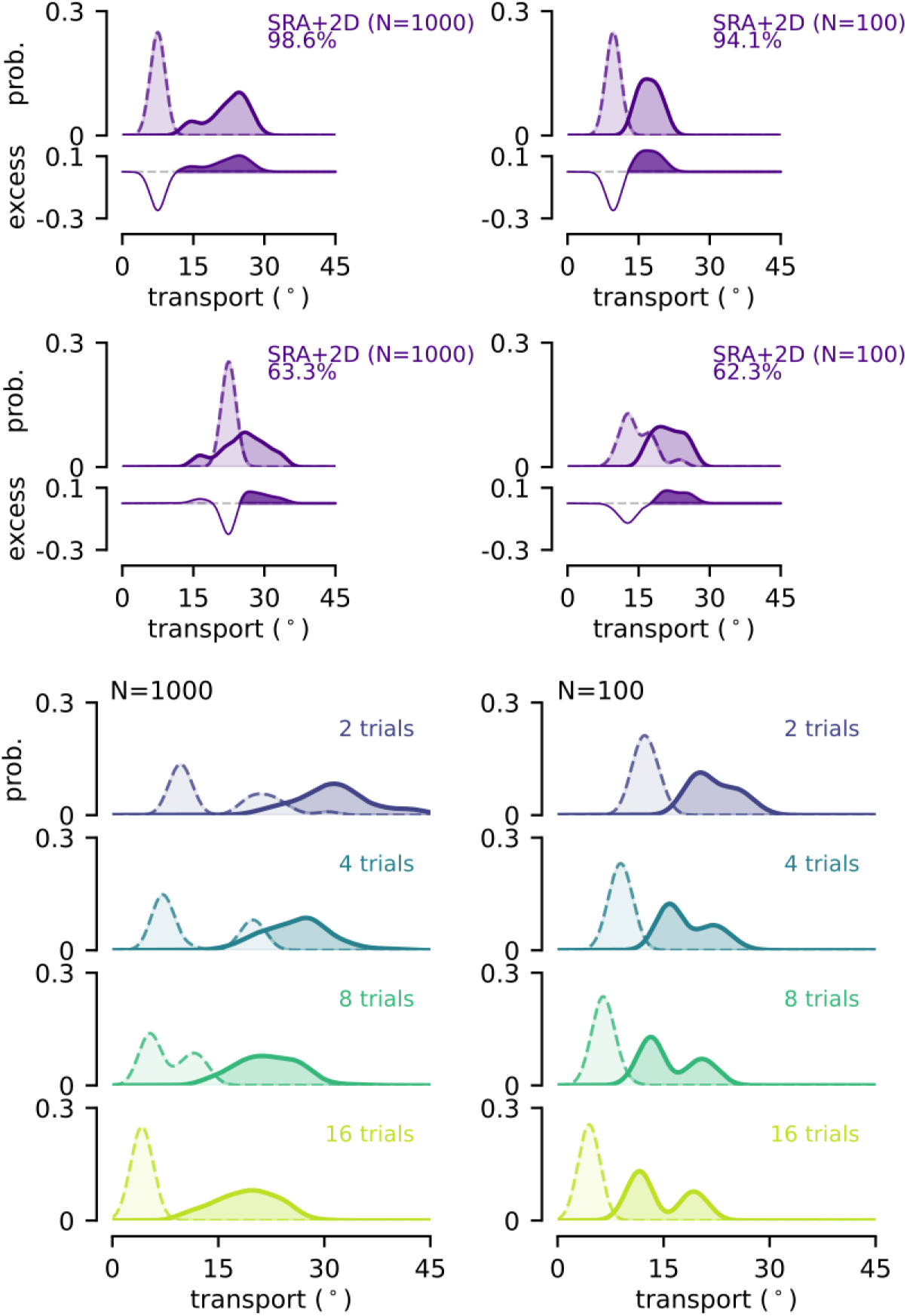
Bimodal Model Null Estimation Boundaries. Four randomly initialized iterations of the SRA+2D-driven network demonstrate varied noise distributions. Beyond demonstrating that the model’s subspace rotations significantly exceed those expected from finite estimation noise, repeated simulations consistently revealed two distinct null distribution peaks. When the network’s neural state remained stable across a trial block, the temporal shuffle control successfully isolated pure finite-sample estimation noise, establishing a theoretical baseline noise floor of 7^○^ 10^○^. Notably, this lower bound was observed *in vivo* for the mouse ALM data, where metastable local dwell times decrease the likelihood of within-block rank switches (Figure 4 and Supplementary Figures S25-S27). However, when the artificial time boundaries of a block happen to straddle an abrupt switch, the resulting temporal mixture inflates the apparent noise estimate to 20^○^ 25^○^, precisely matching both our primate reaching data and session-wide noise estimates in the literature.^62^ This suggests that the observed empirical null distributions in Figure 2 and Supplementary Figures S10,S11 were a signature of geometric undersampling, reflecting both the increased task constraints on redundancy (Figure 5f) and the long experimental block timescales (1-2 minutes) relative to the fast activity timescales of individual adapting neurons (100s of milliseconds).

## Notes

### Competing Interest Statement

The authors have declared no competing interest.

### Summary of Updates

Manuscript revised after further analysis and additional data included.

